# Recapitulating Parkinson’s pathology in human iPSC dopaminergic neurons reveals new mechanistic insights into Lewy body formation and heterogeneity

**DOI:** 10.1101/2025.10.26.684610

**Authors:** Anne-Laure Mahul-Mellier, Lukas van den Heuvel, Maxime Teixeira, Manel L. N. Boussouf, Gaspard Oudinot, Amélie Thonet, Davide Speri, Yllza Jasiqi, Christina Ulrich, Razan Sheta, Walid Idi, Mary Croisier, Stéphanie Clerc-Rosset, Jérôme Blanc, Graham Knott, Abid Oueslati, Hilal A. Lashuel

## Abstract

The accumulation of alpha-synuclein (aSyn) into intraneuronal inclusions of heterogeneous morphology, known as Lewy bodies (LB), is one of the defining diagnostic features of Parkinson’s disease (PD); yet, our understanding of the mechanisms underpinning their formation and heterogeneity remains incomplete. Here, we present a human isogenic iPSC-derived dopaminergic neuron (iDA) model that faithfully recapitulates the diverse biochemical, morphological, and ultrastructural features of LB neuropathology in PD. Unlike other iDA seeding models, our model does not rely on aSyn overexpression, mutations, or genetic engineering, making it a more physiologically relevant system for studying PD. We demonstrate that the iDA model accurately reproduces the temporal relationships between neuritic and cell-body aSyn pathology, recapitulating the full biochemical spectrum, post-translational modifications (PTM), and morphological diversity of aSyn aggregates found in human PD tissue. Moreover, our work provides critical insight into how different pathways to aSyn fibrillization and the complex interaction between aSyn fibrils and membranous organelles influence the morphological diversity of LB-like inclusions. This model represents a versatile platform for investigating the mechanisms of pathology formation, maturation, and neuronal dysfunction, as well as supporting the development of diagnostics that capture the diversity of aSyn pathology in PD and related synucleinopathies.

## Introduction

Parkinson’s disease (PD) is a progressive neurodegenerative disorder for which there are no effective therapies or early diagnostic tools^1^. The primary diagnostic neuropathological hallmarks of PD are the progressive degeneration of dopaminergic (DA) neurons in the substantia nigra (SN) pars compacta and accumulation of misfolded and aggregated forms of the presynaptic protein of alpha-synuclein (aSyn) in the form of the neuritic (Lewy neurites (LNs)) and cytoplasmic aggregates and inclusions, Lewy bodies (LB)^1^. The accumulation of misfolded aSyn aggregates is also observed in the brains of healthy aged individuals as well as in several other neurodegenerative disorders, collectively known as synucleinopathies. However, whether aSyn misfolding, aggregation, and inclusion formation are causes or consequences of the disease process remains a topic of active debate and investigation. This uncertainty is primarily due to the imperfect correlation between the presence and load of pathological aSyn aggregates, such as LB and LNs, and the extent of neurodegeneration or the severity of PD symptoms.

Genetic mutations and multiplications of the SNCA gene, which encodes aSyn, or polymorphism in the promoter regions that regulate aSyn expression, have been linked to familial forms of PD or increased risk of PD^2,3^. Furthermore, the detection of aSyn aggregates in the CSF of PD patients, through the use of aSyn seed amplification assays, enables the diagnosis of PD with greater than 95% accuracy^4,5^. This CSF seeding activity seems to correlate well with the presence or absence of aSyn fibrillar pathology in the brain, suggesting that aSyn fibrillar aggregates in the CSF correlate with brain LB-related pathology^6^. Finally, the presence of aSyn pathology in the brain correlates with cognitive decline in patients with PD and other synucleinopathies, e.g., Alzheimer’s disease^3,7-10^. Despite converging evidence supporting a causative role for aSyn pathology in PD, several observations challenge its central role in the pathogenesis of all PD subtypes. These include the presence of aSyn pathology in the brains of aging healthy individuals^11,12^, the lack of consistent correlations between aSyn pathology and neurodegeneration^13^, and the absence of aSyn pathology in the brains of some patients with LRRK2 mutations who exhibit PD symptoms similar to those of sporadic PD^14^. Two recent papers, using different proximity ligation assays and aSyn antibodies, demonstrated that widespread accumulation of oligomeric forms of aSyn occurs in the LB-negative LRRK2 mutation cases^15,16^.

Several factors may contribute to the discrepancy in understanding the relationship between aSyn aggregation, neurodegeneration, and PD^2,3^. These include: 1) the use of terminology that assumes uniformity among aSyn aggregates, i.e. the use of the terms aggregates or pathology to describe all forms of aSyn aggregates, despite extensive evidence of significant morphological and biochemical heterogeneity in aSyn aggregates across the brains of PD patients and other synucleinopathies; and 2) the lack of cellular and animal models that enable investigating the evolution of LB or accurately replicate the complexity and heterogeneity of aSyn aggregation and pathology formation observed in the human brain^17^.

A closer examination of Lewy pathologies in human brains reveals their morphological diversity and significant differences in the distribution of various molecular markers of LB (e.g., aSyn, ubiquitin, neurofilaments), with the most notable differences observed between brainstem (classical) LB and cortical (ring-like) LB^18-23^. Brainstem LB, typically found in the SN and other regions of the brainstem, are associated with the motor symptoms of PD. In contrast, cortical LB found in the hippocampus and neocortex are more frequently linked to cognitive impairments, such as those seen in dementia with Lewy bodies (DLB)^24^. Whether the association with different functional deficits could be attributed to differences in the biochemical, functional, and ultrastructural properties of aSyn aggregates remains unknown, in large part due to the failure to model aSyn pathological diversity in human neurons. Therefore, developing human neuronal models that recapitulate the formation of LB and LNs over time and enable the dissection of their biochemical and ultrastructural properties is essential for investigating the mechanisms of aSyn pathology formation and dissecting the cell-type-dependent relationship between this process and neurodegeneration in PD and other synucleinopathies.

Towards this goal, we developed a seeding-based isogenic iPSC-derived dopaminergic neuronal (iDA) model of aSyn LB formation that enables the investigation of aSyn fibrillization, pathology progression, and maturation under physiologically relevant conditions, specifically, in the absence of aSyn overexpression. iDAs neurons treated with preformed fibrils (PFF) of aSyn produce LB-like inclusions that closely replicate the diverse biochemical, ultrastructural, and morphological features of human LB pathology observed in idiopathic PD. We demonstrate that the differential distribution of aSyn fibrils and organelles, along with the specific types of membranous organelles that co-accumulate with aSyn inclusions, underlie the morphological diversity of LB inclusions in this model. In-depth characterization of these diverse inclusions suggests distinct mechanisms for the formation of aSyn pathology, including both organelle-dependent and organelle-independent pathways. Our work not only enhances the framework for studying LB maturation but also contributes to the standardization of seeding models, providing a more faithful and reproducible representation of the complex pathology observed in PD. This integrated approach offers a refined working model and a valuable platform for testing therapeutics and hypotheses regarding the intricate interplay between aSyn aggregation, organelle dysfunction, and pathology formation in PD and related synucleinopathies.

## Results

### Internalization, processing, and clearance of aSyn PFF seeds over time in iDA

The aSyn seeding model has been extensively studied and characterized in mammalian cell lines and primary neuronal cultures from rodents. In recent years, advancements in human stem cell technology have enabled the differentiation of iPSCs into iDA^25-49^. However, most available protocols for DA differentiation, until recently, required multistep processes taking several months (40 to 90 days)^50-53^. In recent years, a new protocol combining NGN2 programming and dopaminergic patterning has facilitated the rapid and efficient generation of hiPSC-derived midbrain neurons (iDA)^54,55^ (Figure 1A). In this study, we utilized this advanced cellular approach in combination with the seeding model to develop a robust model of PD in human-derived dopaminergic neurons.

**Figure 1.**
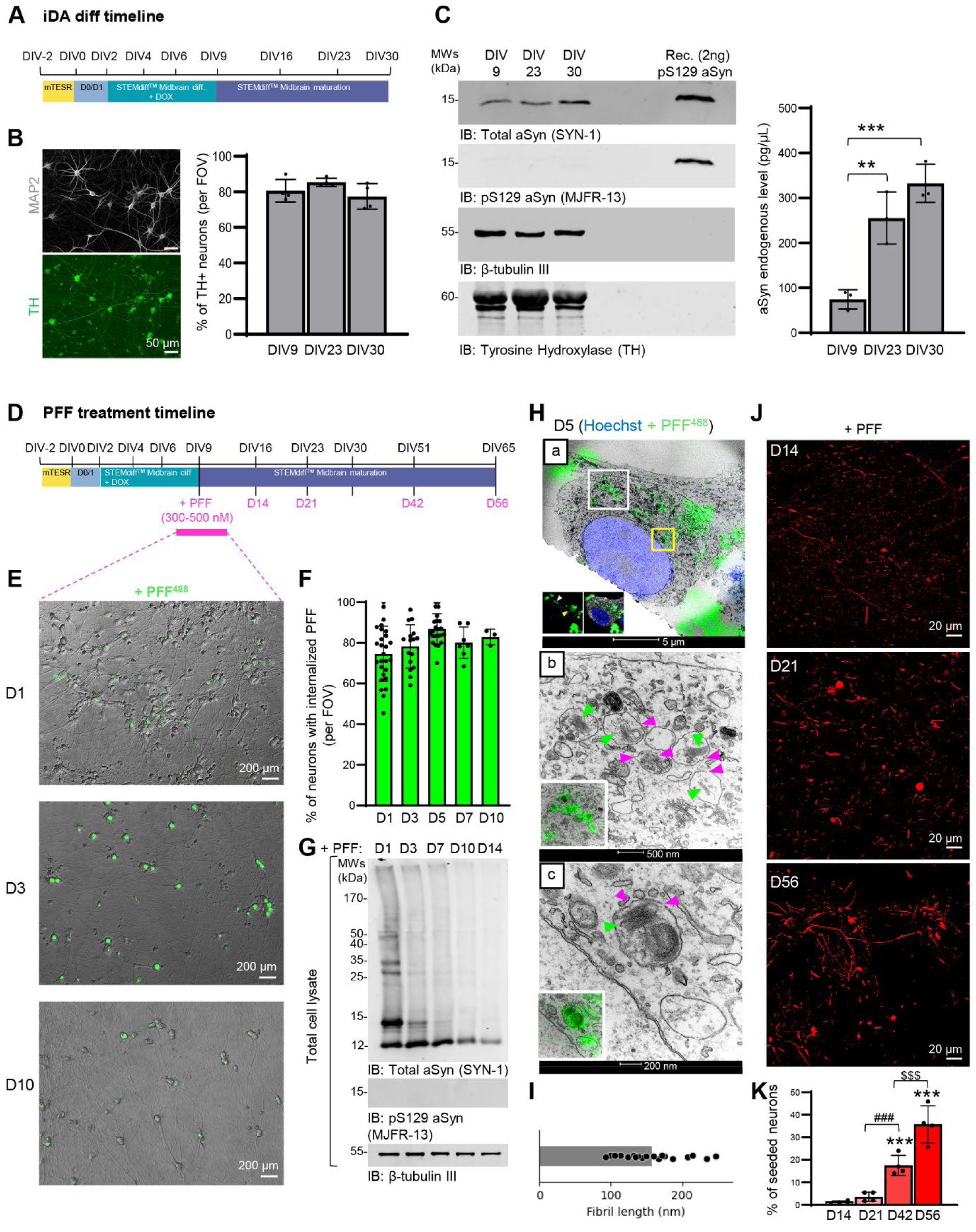
Internalization, processing, and clearance of aSyn PFF seeds over time and subsequent aSyn pathology formation in human iPSC-derived dopaminergic neurons. **A**. Schematic of the iDA differentiation timeline using NGN2 programming for rapid differentiation of human iPSCs into dopaminergic neurons. **B**. ICC confirming the differentiation of iPSCs into dopaminergic neurons at DIV9, with ∼80-90% of neurons expressing TH. Neurons were stained with microtubule-associated protein (MAP2) antibody, and the nuclei were counterstained with DAPI staining. Scale bars = 50 µm. The graph quantifies the percentage of TH-positive neurons at DIV9, DIV23, and DIV30. **C**. WB analysis of total aSyn (SYN-1 antibody) and pS129 aSyn (MJF-R13 antibody) protein levels in iDA cultures at DIV9, DIV23, and DIV30, showing a significant increase in total aSyn levels over time. The immunoblot was probed with β-tubulin III and TH antibodies as loading controls. aSyn concentration (µg/µL) was quantified by densitometric analysis, using the signal intensity of 2 ng of recombinant aSyn protein loaded on the same blot as a reference standard. **D**. Seeding model in iDA. 500 nM of PFF were added at DIV9. Control iDA neurons were treated with PBS buffer, which was used to prepare PFF. **E**. Representative images showing fluorescently labeled PFF (PFF^488^) internalized by iDA neurons at D1, D3, and D10 post-treatment. Scale bars = 200 µm. **F**. Quantification of the percentage of neurons with internalized PFF^488^ at different time points (D1, D3, D5, D7, and D10). **G**. WB analysis showing the processing of PFF over time. Internalized PFF detected by SYN-1 antibody are cleaved into a 12 kDa fragment within 24 hours of uptake. The PFF were not detected by pS129 (MJF-R13) antibody. The immunoblot was probed with β-tubulin III antibody as a loading control. **H**. CLEM images showing PFF within lysosomal-like vesicles at D5, confirming PFF internalization via the endolysosomal pathway. Panel a displays the overlay of the selected cell imaged by confocal microscopy and electron microscopy (EM). Panel b provides a higher magnification of the region highlighted by the white rectangle in panel a. Panel c shows an even higher magnification of the area marked by the yellow rectangle in panel a. Green arrows indicate internalized PFF within intact vesicles, while pink arrows indicate vesicles containing PFF that appear ruptured. Scale bars = 50 µm (panel a); 500 nm (panel b); and 200 nm (panel c). **I**. Size of PFF within vesicles (∼156 nm). **J**. Representative images illustrating the development of pS129 pathology at D14, D21, and D56 after PFF treatment. pS129 aggregates (81A antibody) are primarily detected in neurites at D14, with somatic inclusions emerging at D21 and increasing by D56. Additional staining for MAP2, TH, and DAPI is shown in Figure S5A. **K**. Quantification of pS129 pathology in neurons at different time points. **C, K**. The graphs represent the mean ± SD of 3 independent experiments. C, **p<0.001, ***p<0.0001 (ANOVA followed by Tukey HSD *post-hoc* test, DIV9 vs. other time-points). K. ***p<0.0001 (ANOVA followed by Tukey HSD *post-hoc* test, D14 vs. other time-points); ###p<0.0001 (ANOVA followed by Tukey HSD *post-hoc* test, D21 vs. D42); $$$p<0.0001 (ANOVA followed by Tukey HSD *post-hoc* test, D42 vs. D56).

First, we validated the effectiveness and reproducibility of the rapid protocol developed by Sheta et al. (Figure 1A-C). Immunocytochemistry (ICC) combined with confocal imaging confirmed the differentiation of isogenic iPSCs into iDA (Figures 1B and S1). Among the neuronal population (MAP2-positive), approximately 80% of the neurons expressed tyrosine hydroxylase (TH), the rate-limiting enzyme in dopamine synthesis and a marker of dopaminergic neurons, after 9 days in vitro (DIV) (Figure 1B). This differentiation efficiency was maintained over time, with a consistent ∼80% of the neurons expressing TH at DIV30. Western blot analysis of total cell lysates from these iDA cultures confirmed the expression of the TH protein. Additionally, we measured the concentration of endogenous aSyn in these cultures over time. Using recombinant monomeric aSyn as a standard, we estimated that endogenous aSyn was expressed at a concentration of 84 pg/μl at DIV9, which increased 3.5-fold by DIV30 to reach ∼300 pg/μl (Figure 1C). At this protein expression level, we could not detect any signal for S129-phosphorylated aSyn (pS129) in iDA neurons (Figure 1C).

Next, the iDA cultures were treated at DIV9 with aSyn PFF (Figures 1D and S2) to induce seeding and formation of aSyn pathology as previously described^40,56-58^. To assess the internalization of seeds by iDA neurons over time, we used WT human PFFs fluorescently labeled with Atto 488 (PFF^488^) (Figure S2C). A concentration of 300 nM was chosen to allow reliable quantification of uptake without saturating the fluorescent signal.

One day after the addition of PFF^488^ to the extracellular media, we observed that approximately ∼70-75 % of the neurons had internalized the PFF seeds (Figures 1E-F and S3A). This uptake and accumulation, observed mainly as puncta in the cytosol, increased to approximately 80% of the neurons with internalized PFF by Day 3 (D3) post-treatment. The presence of the PFF remained detectable at the same level 10 days (D10) post-treatment (Figures 1E-F and S3A). This aligns with previous studies showing that PFF are rapidly and efficiently internalized across various cellular models, including mammalian cell lines^25,40,59-61^, primary neurons^25,28,40,57,60,61^, and iPSCs^25,28,31,34,38,40,41,46,60,62-65^, as well as in vivo following brain injection^40,60,66,67^.

Biochemical analysis of total cell lysates from PFF-treated iDA cultures (Figures 1G and S3B) showed that within 24 hours of treatment, the internalized PFF were C-terminally cleaved, as indicated by the appearance of a 12 kDa band. By 7 days (D7) post-treatment, the full-length aSyn (∼ 15 kDa band) was barely detectable. Although the truncated species were gradually cleared over time, a residual 12 kDa band remained detectable 14 days (D14) after PFF internalization. Additionally, the detection of high-molecular-weight (HMW) aggregates decreased progressively over time. As expected, WB analysis also confirmed that the PFF are not phosphorylated at serine S129. Similar observations have been reported by our group and others in primary neurons, iDAs, and mammalian cell lines treated with PFF, as well as in mice injected with PFF^40,60^. Live imaging also showed that WT human PFFs fluorescently labeled with Atto 647 (PFF^647^, Figure S2D) are rapidly internalized by iDA cells through the endolysosomal pathway within minutes after adding PFF to the extracellular medium (Figure S3C-E) as previously described^25,40,60-62^. Consistent with these results, CLEM combined with confocal microscopy confirmed the accumulation of the PFF^488^ in lysosomal-like vesicles (Figures 1H-I and S4). aSyn PFF inside these vesicles measured approximately ∼156 nm in length (Figure 1I), consistent with the size distribution observed by electron microscopy (EM) during the characterization of the sonicated PFF preparation, which ranged from ∼50 nm to 250 nm (Figure S2A). We also observed that the membranes of the vesicles containing the PFF appeared to be ruptured (pink arrows) or exhibited a less-defined, diffuse membrane structure (Figures 1H, panel c and S4). This is in line with a recent study showing that internalized PFF trigger the perforation of the endolysosomal vesicles^26,62^. Altogether, our results demonstrate that PFF are internalized and processed similarly in iDA cultures, as in other neuronal seeding models, such as PFF-treated primary WT or KO neurons.

### aSyn PFF induce pS129 pathology in iDA

Next, we monitored the formation, maturation, and subcellular localization of pS129 pathology in iDA cells over time. We conducted ICC at D7, D14, D21, and D56 on iDA cells treated with 500 nM of unlabelled human aSyn PFF. No pS129 pathology was observed after D7 (Figure S5A). At D14, pS129 pathology was detected primarily in the neurites, with less than 1% of neurons showing cytosolic seeded aggregates (Figures 1J-K and S5A). Interestingly, while neuritic pathology in primary neurons was mainly observed in dendrites (MAP2+)^57^, in iDA neurons, pS129 aggregates were barely seen in MAP2+ dendrites (Figure S5A). Instead, the neuritic aggregates were mostly located in the axons (β-tubulin+; MAP2-) of the iDA cells (Figure S5B). This aligns with human pathology, where axonal projections are more frequently and consistently affected by pS129 aggregation than dendrites^68^. By D21, both neuritic and somatic pS129 pathology were observed; however, somatic pathology was present in fewer than 3% of the iDA neurons (Figures 1J-K and S5A). At D56, approximately 30-45% of the iDA neuronal cell bodies exhibited seeded aggregates (Figures 1J-K and S5A). It is worth noting that while 70 nM (∼1 µg/mL) of human seeds were sufficient to induce robust pathology in mouse primary neurons^57,58^, a minimum concentration of 500 nM (∼7 µg/mL) of human PFF was required to achieve a significant level of pathology over time in iDA (Figures 1J-K and S5A). This finding aligns with other studies that employed similar concentrations of PFF in iPSC-derived seeding models, where the reported effective range of PFF concentrations spans from ≤1 µg/mL^25,26,36,46,63^ to 2-5 µg/mL^31,34,38,40,41,60,64^ or 10-15 µg/mL ^28,38,45,47,69^ and up to 60µg/mL^35^. Even in lines with aSyn gene multiplication, a minimum of 5 µg/mL and up to 10 µg/mL was used^38,45,69^. It is worth noting that under these conditions, mouse PFF were unable to induce any pathology over time in our hands (Figure S5C). A likely explanation for the greater pS129 pathology observed in mouse primary neurons compared to iDA cultures is the fact that the sequence of the mouse aSyn-PFF and the substrate is the same, together with markedly higher endogenous mouse aSyn levels, estimated to be 7–10 times higher at early differentiation stages (DIV7–9) and around 4-fold higher in more mature neurons (DIV30– 35) (Figure S6A-B). These findings underscore the importance of endogenous aSyn concentration as a key determinant of the initiation and the extent of pathological burden in neurons.

We were also intrigued by the fact that, despite ∼80% of the iDA neurons internalizing PFF seeds, only 3% exhibited pathology at D21. This was significantly lower than the 40% observed in primary mouse neurons under similar conditions^57^. We hypothesized that variations in endogenous aSyn levels among neurons at DIV9, when seeds were added, or other differences in cellular factors that influence the initiation of seeding might explain why pathology developed in only a minor population at early stages (Figure 1J). To estimate the local concentration of aSyn at the single-cell level, we performed ICC on untreated iDA cultures at DIV9. Neurons were co-stained with β-tubulin III (for the total neuronal population) and TH (for dopaminergic neurons) antibodies. Total aSyn levels in each cell were assessed using SYN-1 (epitope: 91-99) antibody. High-content imaging analysis (HCA) revealed that the concentration of endogenous aSyn was similar across non-dopaminergic (β-tubulin III-) and dopaminergic (TH+) neurons, regardless of whether TH expression levels were high or moderate (Figure S6C-D). However, among TH+ neurons, 7% exhibited aSyn expression levels 2–4 times higher than the population average (Figure S6E). It is likely that these neurons with elevated aSyn levels are where seeding predominantly occurs, which may explain why, at D21, pathology was observed in only ∼3-5% of the total neuronal population (Figure 1J). Overall, our findings highlight that although more than 80% of iDA cells internalized PFF (Figure 1E-F), seeding initiation is a slow stochastic process occurring in only a few neurons at early stages.

### Seeded aggregates in iDA neurons exhibit LB pathological markers and a broad morphological spectrum similar to human brain pathology

Next, we conducted an in-depth characterization of pS129 pathology in both neurites and cell bodies, focusing on how our seeded aggregates reproduce PD-specific features observed in patient brain tissues. We compared the LB inclusions formed in PFF-treated iDA neurons to those found in human PD brain tissues, first assessing whether our seeded aggregates could replicate the full spectrum of LB morphologies observed in patients (Figure 2). Additionally, we examined key pathological markers (p62 and ubiquitin) (Figure 3), PTM signatures (Figure 4), and the ultrastructural organization and morphology characteristic of *bona fide* LB inclusions (Figures 5-7), as well as how organelle sequestration and dysfunction contribute to LB formation and maturation(Figures 8-9). First, we observed a broad morphological spectrum of aggregates formed over time (Figure 2). In addition to neuritic pathology, we classified the somatic-seeded aggregates into six categories: filamentous-like aggregates, tiny dot-like aggregates, speckled-like aggregates, and three types of LB, including dense-like LB, ring-like LB, and LB (dense- or ring-like) surrounded by speckled aggregates (Figure 2A). This morphological diversity was evident in both D21 (Figure 2B) and D56 (Figure 2C).

**Figure 2.**
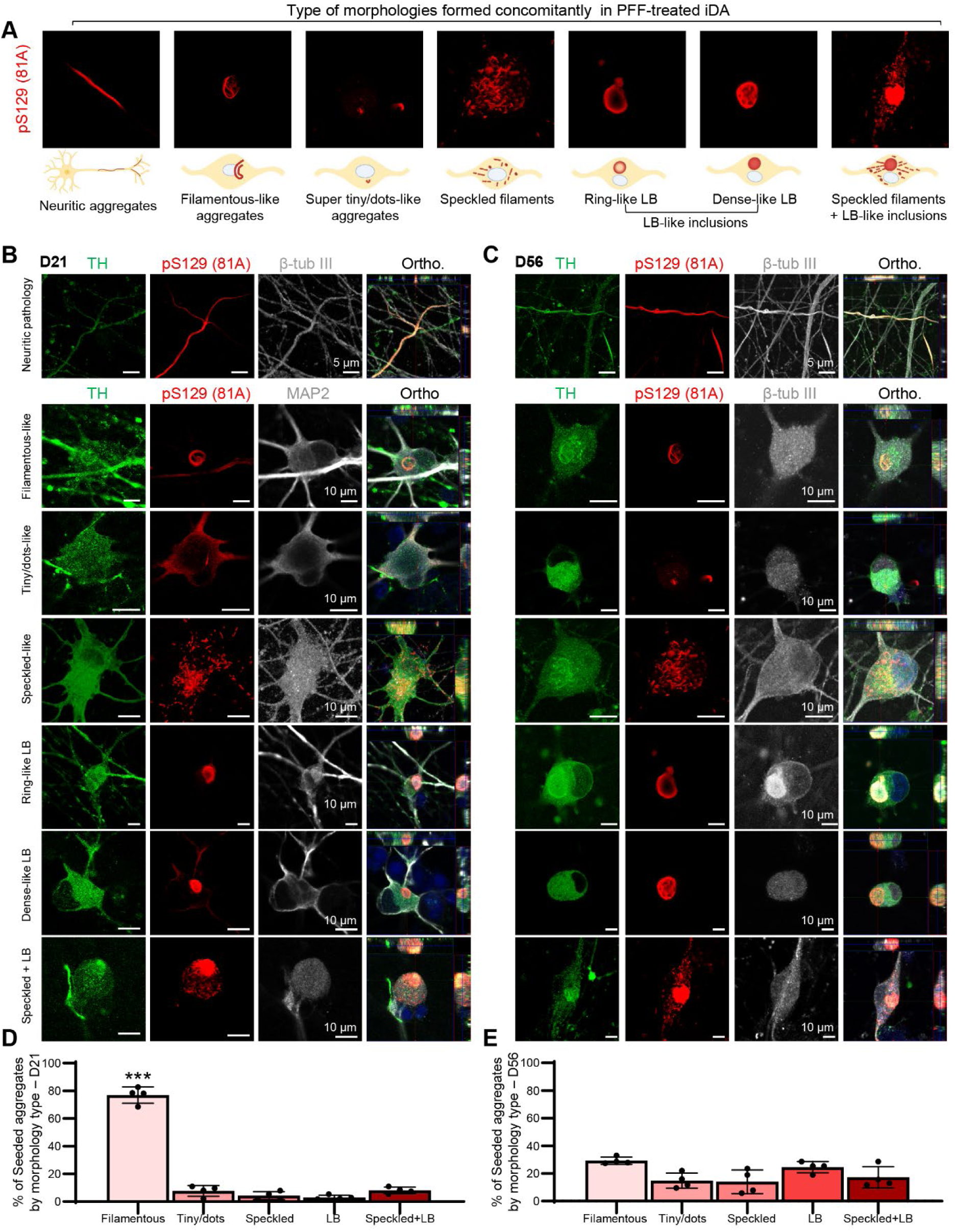
Seeded aggregates in iDA neurons exhibit a wide morphological spectrum similar to human brain pathology. **A**. Schematic representation of the different types of morphological aggregates formed in PFF-treated iDA neurons: neuritic aggregates, filamentous-like aggregates, super tiny/dot-like aggregates, speckled-like aggregates, ring-like LB, dense-like LB, and LB-like inclusions. Representative images shown in panel A are presented in full, with an additional neuronal marker, in Figure 2 or Figure S9. **B-C**. Representative images of iDA neurons at D21 (B) and D56 (C) showing the different aggregate morphologies positively stained with pS129 antibody (81A, red). Neurons were counterstained with TH (green), β-tubulin III or MAP2 (gray) antibodies. Orthogonal projections (Ortho.). **B-C**. Scale bars **=** 5 µm (top row of panels B and C), 10 µm (bottom row of panels B and C). **D-E**. Quantification of the percentage of each type of seeded aggregate at D21 (D) and D56 (E), showing a predominance of filamentous-like aggregates at D21 and a significant increase in LB-like inclusions by D56. The graphs represent the mean ± SD of 3 independent experiments. ***p<0.0001 (ANOVA followed by Tukey HSD *post-hoc* test, filamentous vs. other morphologies).

**Figure 3.**
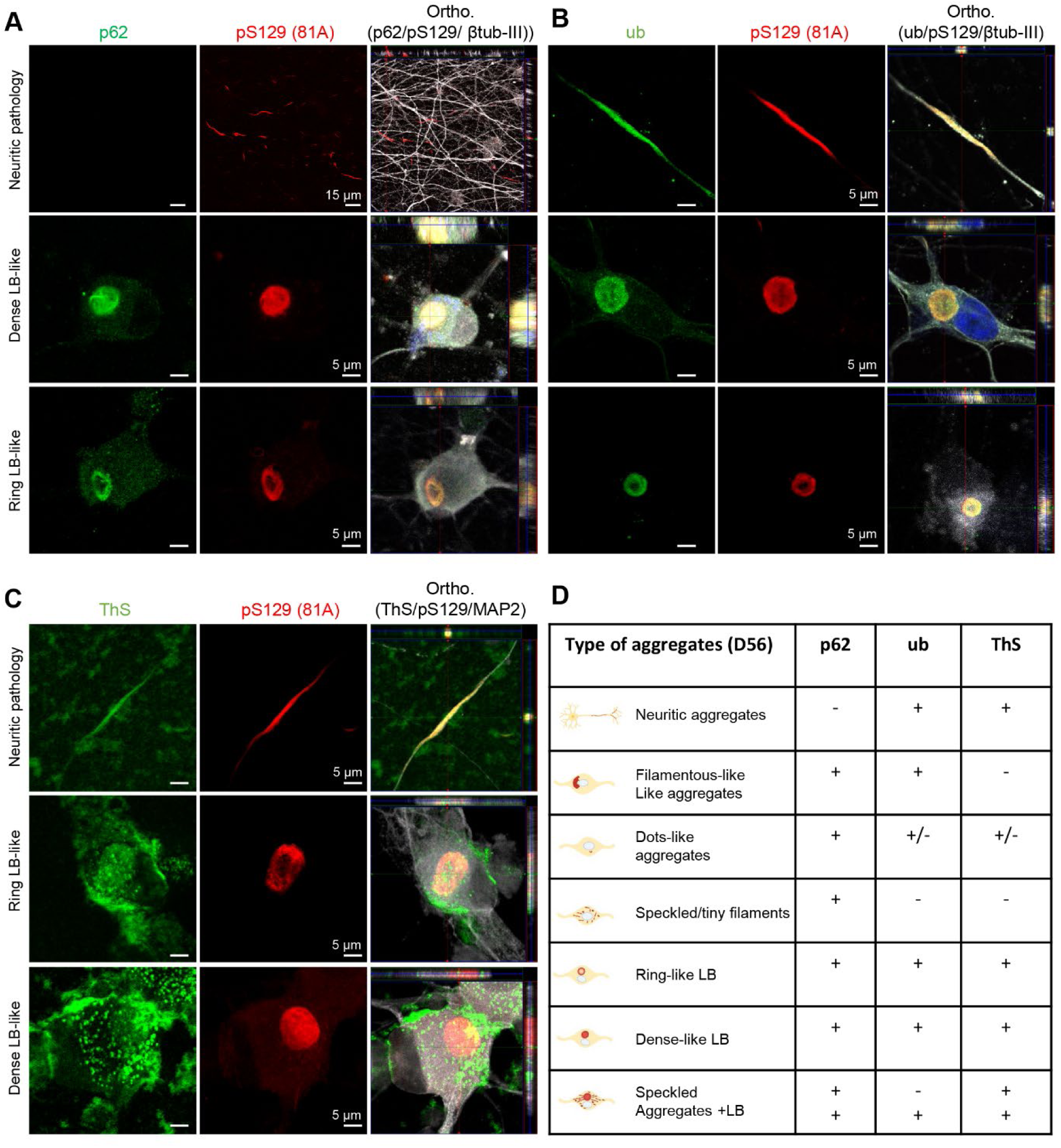
Seeded aggregates in iDA neurons exhibit the LB pathological markers similar to human brain pathology. **A**. Representative images showing the differential sequestration of p62 and pS129 pathology in somatic and neuritic pathology at D56. p62 staining colocalized with the pS129-positive dense-like and ring-like LB-like inclusions, but not with the neuritic pathology. **B-C**. Ubiquitin (ub, B) and Thioflavin S (ThS, C) staining confirming β-sheet amyloid-like species colocalized with both neuritic pS129 pathology and LB-like inclusions at D56. **A-C.** Neurons were counterstained with β-tubulin III or MAP2 antibody. **D**. Summary table showing the differential sequestration of p62, ubiquitin, and ThS in different types of seeded aggregates at D56 (see additional staining of the different types of seeded aggregates in Figure S8). Orthogonal projections (Ortho.). Scale bars = 5 µm (**A-C**).

**Figure 4.**
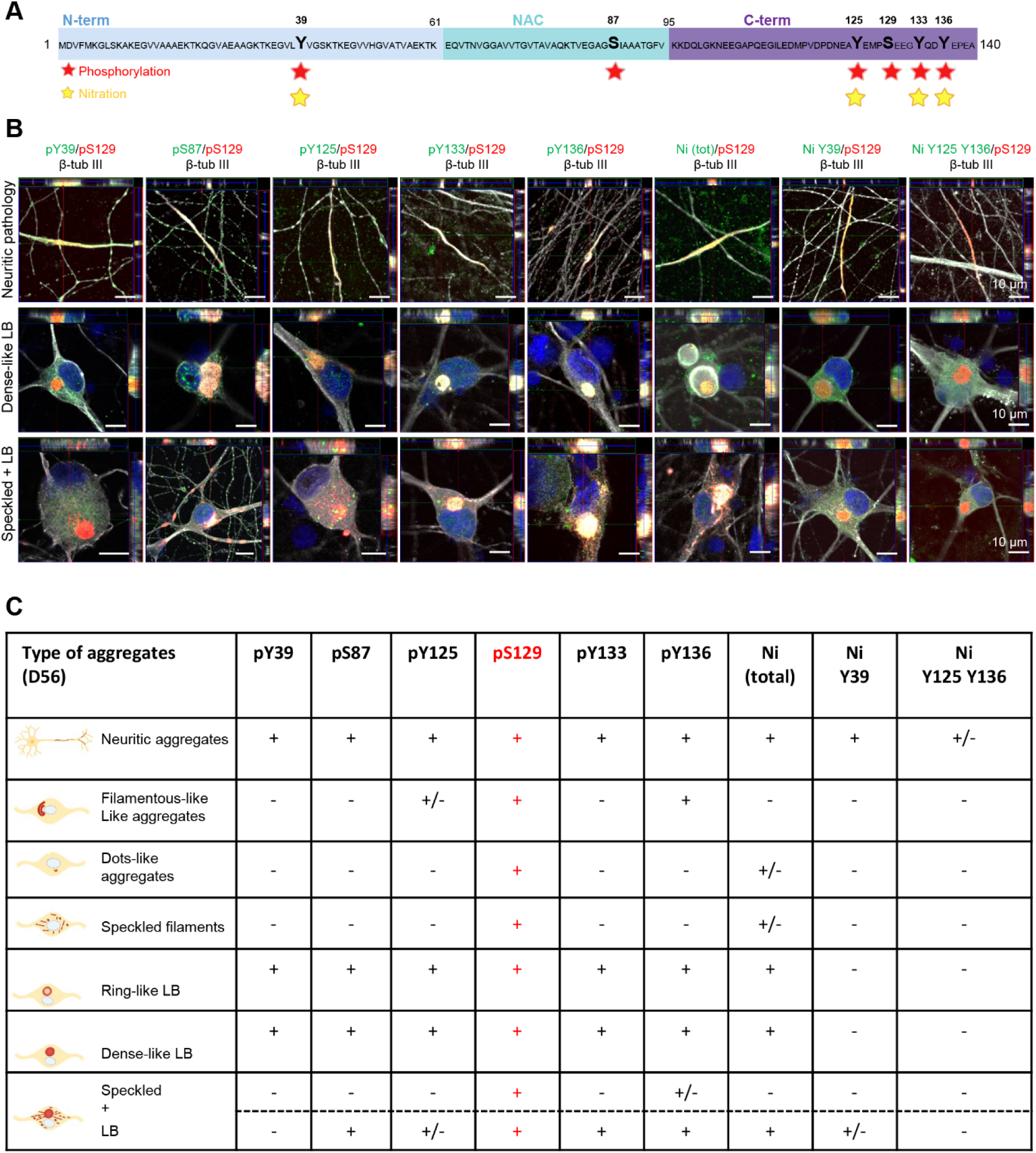
The iDA seeding model replicates the PTM signature associated with human pathology. **A.** Schematic representation of PTM analyzed in this study. These include phosphorylation at residue Y39, S87, Y125, S129, Y133 and Y136, and nitration at Tyrosine residues (nY39, nY125, nY133, nY136). All of these PTM have been previously identified in aSyn pathology in post-mortem brain tissues from patients with PD and MSA. **B.** Representative confocal images at D56 illustrate the colocalization of pS129 with the PTM described in (**A**) across distinct aSyn aggregate morphologies within neuronal cell bodies. Images are shown as orthogonal projections of merged channels, including pS129, βIII-tubulin, and the respective PTM-specific antibodies. Scale bar = 10 µm. Individual channels and additional aggregate morphologies, including dot-like, ring-like, filamentous, speckled, and neuritic inclusions, are shown in **Figure S9**. **C.** Summary table displaying the distribution of specific PTM across the different aSyn aggregate types observed at D56, highlighting the diversity and selectivity of PTM profiles associated with each morphology.

**Figure 5.**
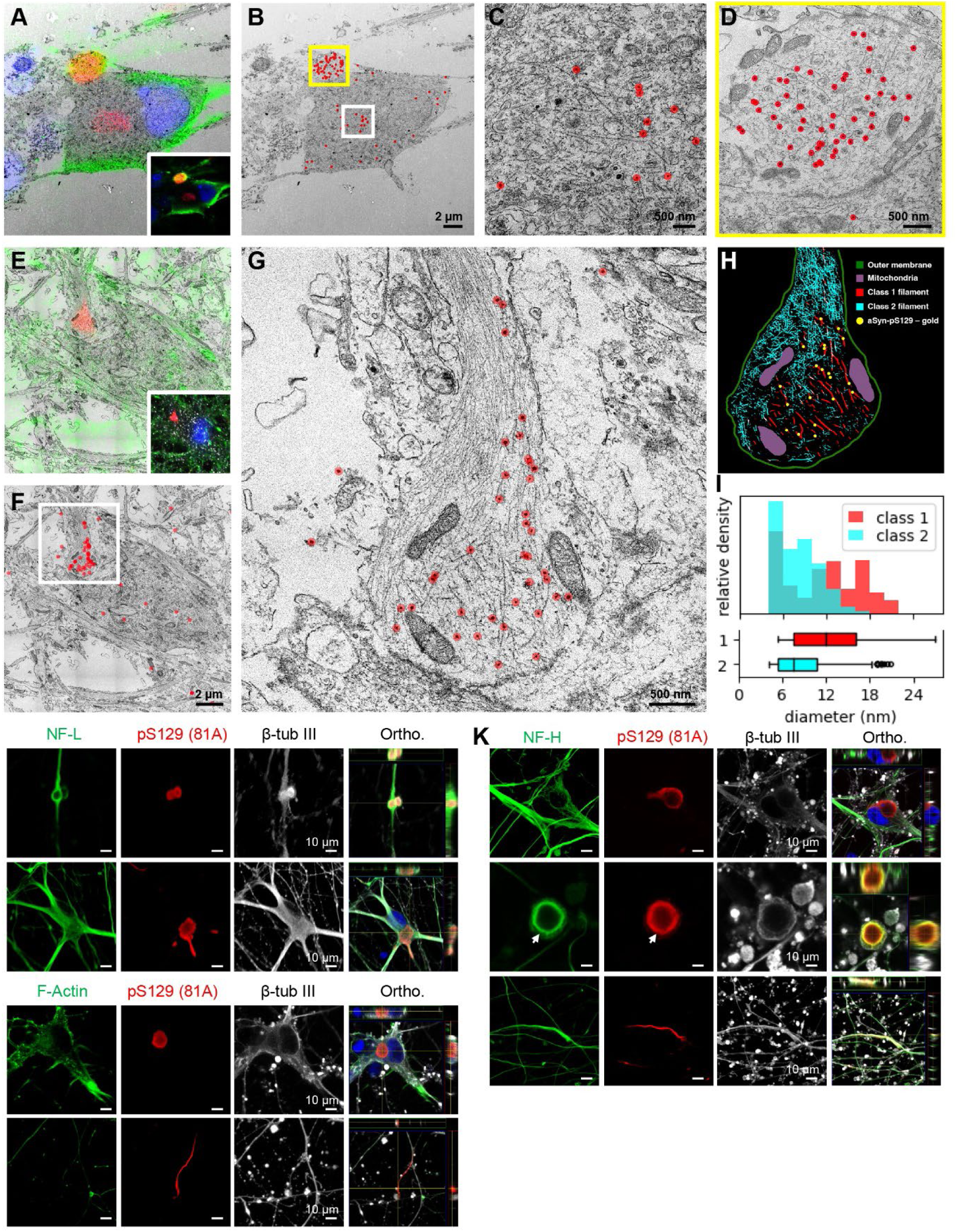
Seeded aggregates in iDAs are rich in fibrils that label positively for pS129 aSyn and that are distinct from cytoskeletal filaments. **A.** ICC and EM overlay of a cell stained with fluoro-gold conjugated antibodies binding to pS129 aSyn. The cell has an aggregate in its center and is attached to a small, dead cell (top) that is also immunopositive for pS129 aSyn. Inset: immunofluorescence only. **B.** Electron micrograph of the same cell, with gold particle detections shown as red dots. **C-D.** Zoomed insets of the aggregate inside the cell cytoplasm (**C**) and the dead cell (**D**) show the association of gold particles with filaments. **E.** IF and EM overlay of another cell stained with fluoro-gold conjugated antibodies binding to pS129 aSyn. **F-G.** Gold particle detections (red dots) agglomerate in the neurite of this cell, where two filament types can be distinguished: thick filaments associated with gold particles and thinner filaments that do not associate with gold particles. **H.** Tomographic, neural-network-based segmentations of gold-associated filaments (class 1) and thin filaments (class 2), together with manual segmentations of mitochondria and the neurite outer membrane. Gold particle detections are shown in yellow. **I.** Thickness measurements of the segmented filaments in class 1 (mean=12.0 nm, std=5.2 nm) and class 2 (mean=7.6 nm, std=3.0 nm) **J, K-L.** Immunolabeling of the cytoskeletal proteins NF-L, NF-H, F-Actin and β-tubulin III revealed no co-localization with pS129 staining in the soma of live cells. Scale bars = 2 µm (**B**, **F**), 500 nm (**C**,**D** and **G**), 10 µm (**K**-**L**).

**Figure 6.**
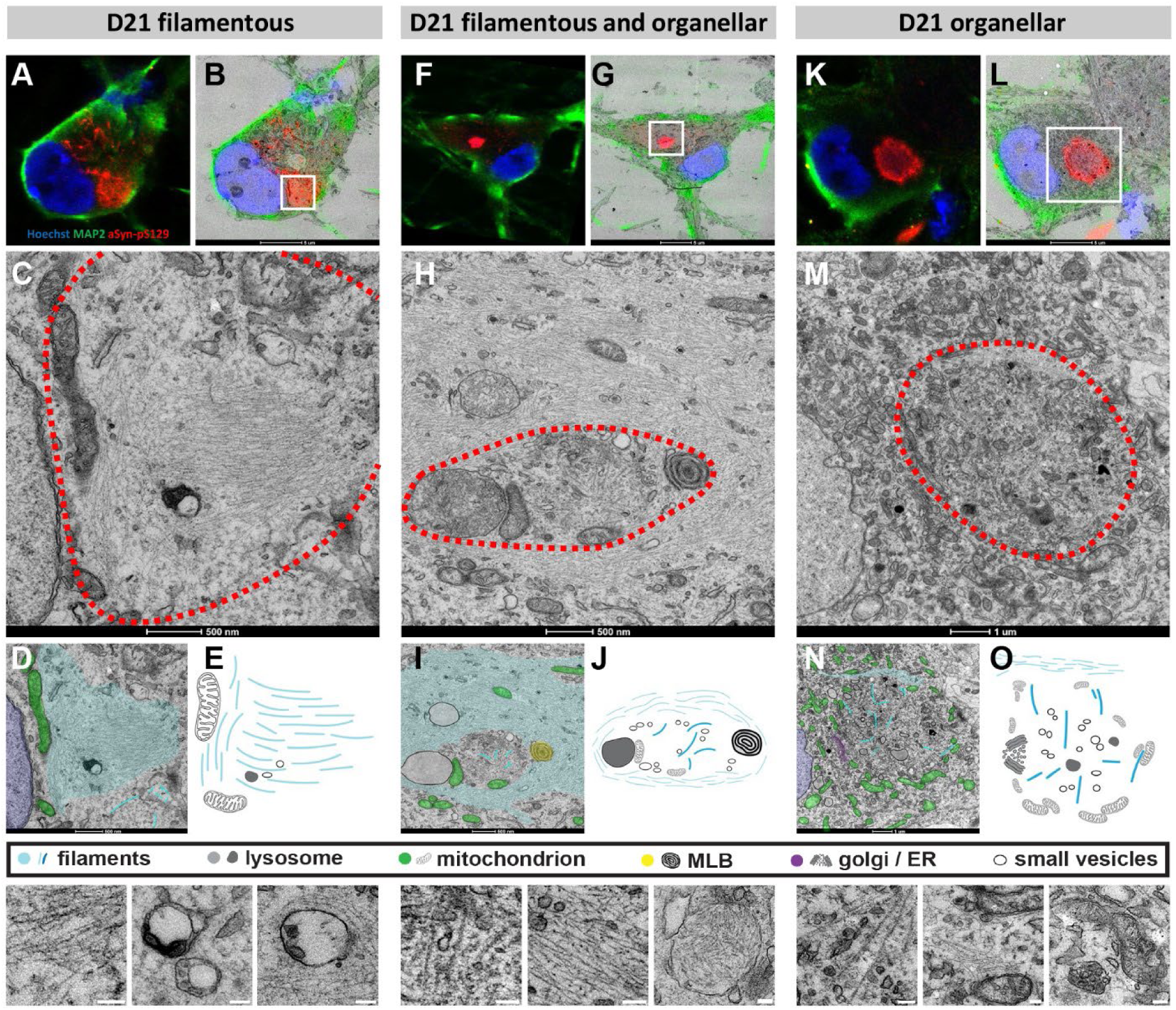
CLEM of pS129 aSyn aggregates at D21 in PFF-treated iDA. Neurons were fixed in 4% PFA, permeabilized by rapid freezing and thawing in the absence of detergent to keep the ultrastructure intact, and immunolabeled with pS129 (81A) antibody. After confocal immunofluorescence imaging, the same cells were subjected to heavy-metal staining, resin embedding, ultrathin sectioning, and imaged with a transmission electron microscope. Aggregates were classified into three groups based on their ultrastructure. **A-E.** A predominantly filamentous ultrastructure, where pS129 immunofluorescence correlates with a dense mesh of thin, parallel filaments. Small vesicles and electron-dense lysosomes are sparsely mixed with the filaments. **F-J.** A filamentous mesh surrounds a dense aggregate of small vesicles, membrane-bound organelles and thick (13 ± 3.5 nm) fibrils, which are sparsely and randomly distributed. The membranous aggregate correlates with bright pS129 immunofluorescence (**F-G**), while the immunofluorescence in the filamentous mesh is less bright and more punctate. **K-O**. pS129 immunofluorescence correlates with an ultrastructure dense in membranes, thick (13 ± 3.5 nm) fibrils and membrane-bound organelles, in particular mitochondria, lysosomes, and small vesicles. Scale bars = 5 μm (**B, G-L**) or 500 nm (**C, H-M**), insets 100 nm.

**Figure 7.**
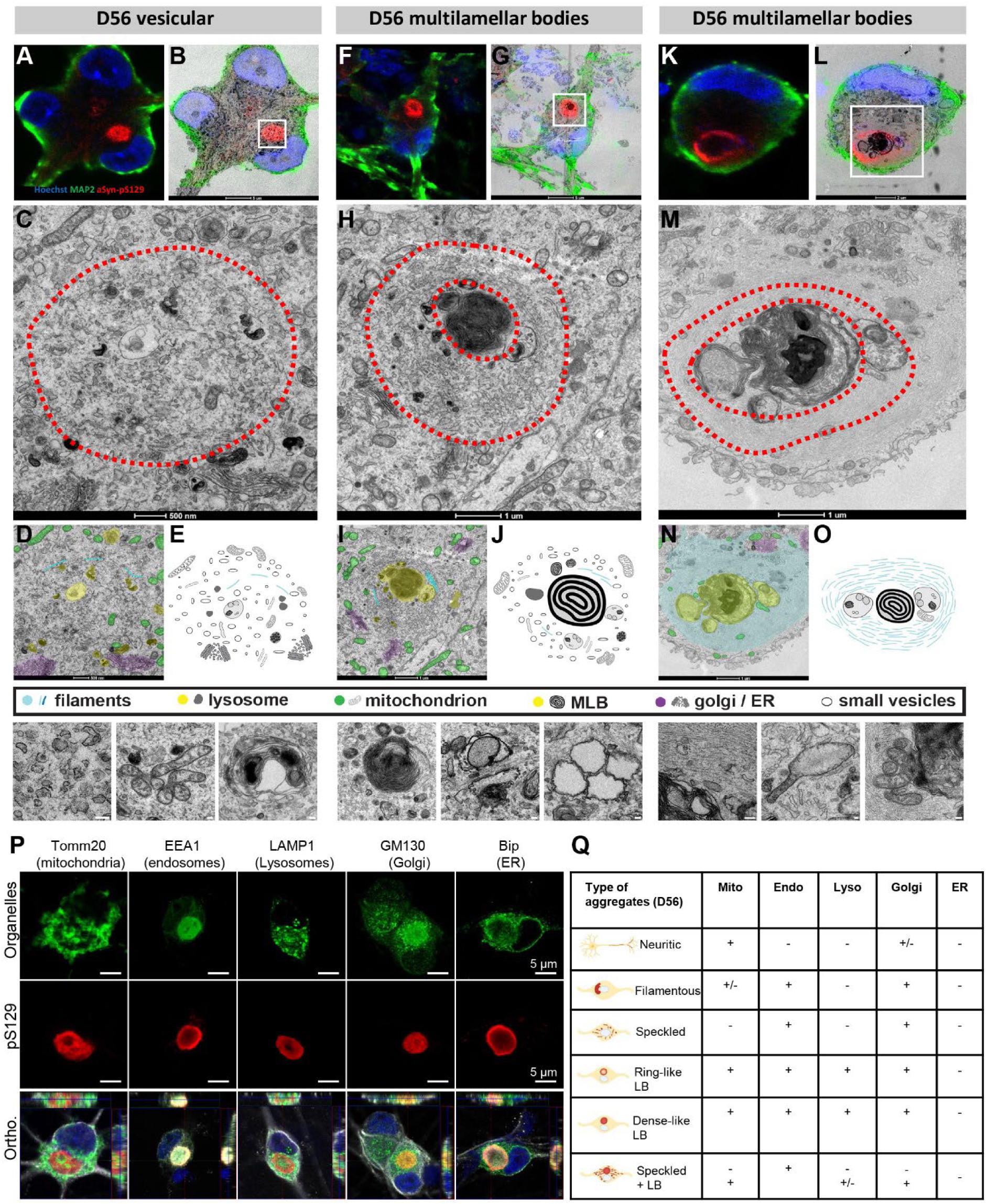
CLEM of pS129 aSyn aggregates at D56 in PFF-treated iDA. Aggregates were classified into three groups based on their ultrastructure. **A-E.** A uniform pS129 immunofluorescence, which correlates with an ultrastructure rich in small vesicles, lysosomes, clustered mitochondria, and long, thick filaments (13 ± 3.5 nm). **F-J.** A ring-shaped pS129 immunofluorescence, of which the centre correlates with the presence of membrane-rich multilamellar bodies (MLB). Surrounding the MLB are small vesicles and sparse, thick filaments (13 ± 3.5 nm). **K-O.** A similar ring-shaped pattern of pS129 immunoreactivity with MLB in its centre, but here surrounded by a dense mesh of parallel, thin (8 ± 2 nm) filaments. Scale bars = 5 μm (**B, G-L**) or 500 nm (**C, H-M**), insets 100 nm. **P.** Representative images show pS129 pathology (red, 81A antibody) alongside markers for key organelles: GM130 for the Golgi apparatus, BiP for the ER, EEA1 for endosomes, LAMP1 for lysosomes, and Tomm20 for mitochondria (all in green). Orthogonal (Ortho.) projections confirm that all these organelles except the ER are sequestered into pS129 LB-like inclusions at D56. Neurons were stained with the β-tubulin III antibody, while nuclei were counterstained with DAPI. Scale bars = 5 µm. **Q.** Summary table depicting the sequestration of cellular organelles across the distinct aSyn aggregate morphologies observed at D56.

**Figure 8.**
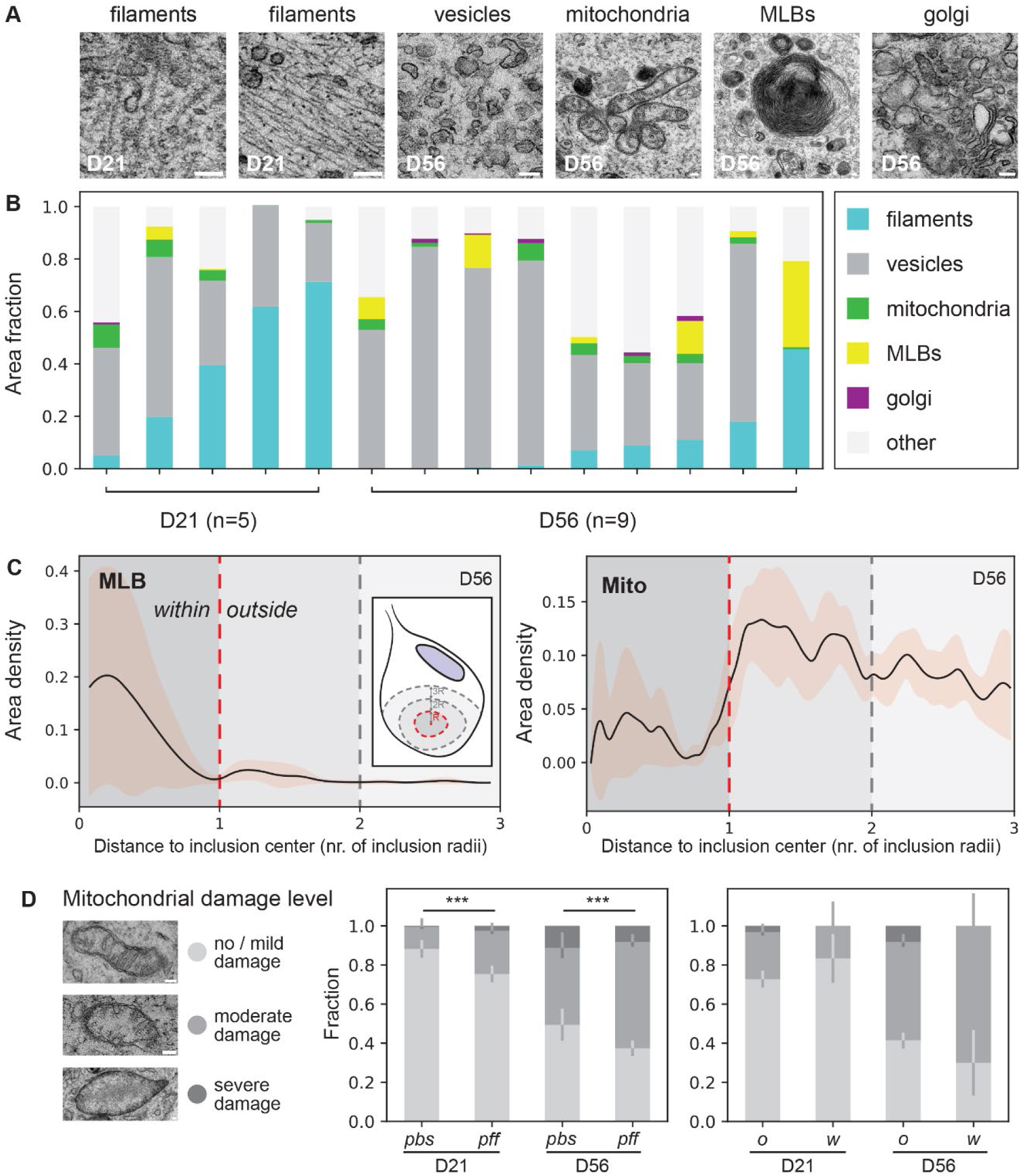
Inclusion composition and mitochondrial damage of seeded aggregates in PFF-treated iDAs analyzed by CLEM. **A.** Representative images of components within aggregates at D21 and D56, notably filamentous ultrastructure (thick filaments of ∼13 nm on the left, thin filaments of ∼8 nm on the right), small vesicles, mitochondria, MLB, and the Golgi apparatus. Scale bars = 100 nm. **B**. Area proportion of the above-mentioned ultrastructural components in electron micrographs through the central slice of five D21 aggregates and nine D56 aggregates. **C.** Area of MLB (left) and mitochondria (right) divided by the total area of the pS129-positive region at different positions within and outside D56 aggregates. A distance of 0 corresponds to the center of the aggregate, 1 to the periphery, 2 and 3 to one and two aggregate radii away from the inclusion, respectively. The black line represents the mean across 9 aggregates in 9 different cells, while the shaded area indicates the 95% confidence interval obtained through bootstrapping. **D.** Three representative images show the classification of mitochondria into three classes according to their level of damage. Scale bars = 100 nm. The left plot shows the mean fraction of each class in PBS control cells or PFF-treated cells with aggregates, at D21 and D56. The right plot shows the fraction of each class inside and outside the inclusions of PFF-treated cells with aggregates, at D21 and D56. Error bars indicate one-half of the 95% confidence intervals for each class, calculated using bootstrapping. A Chi-squared test indicates significance between PBS- and PFF-treated cells with ***p < 0.0001 for D21 and D56.

**Figure 9.**
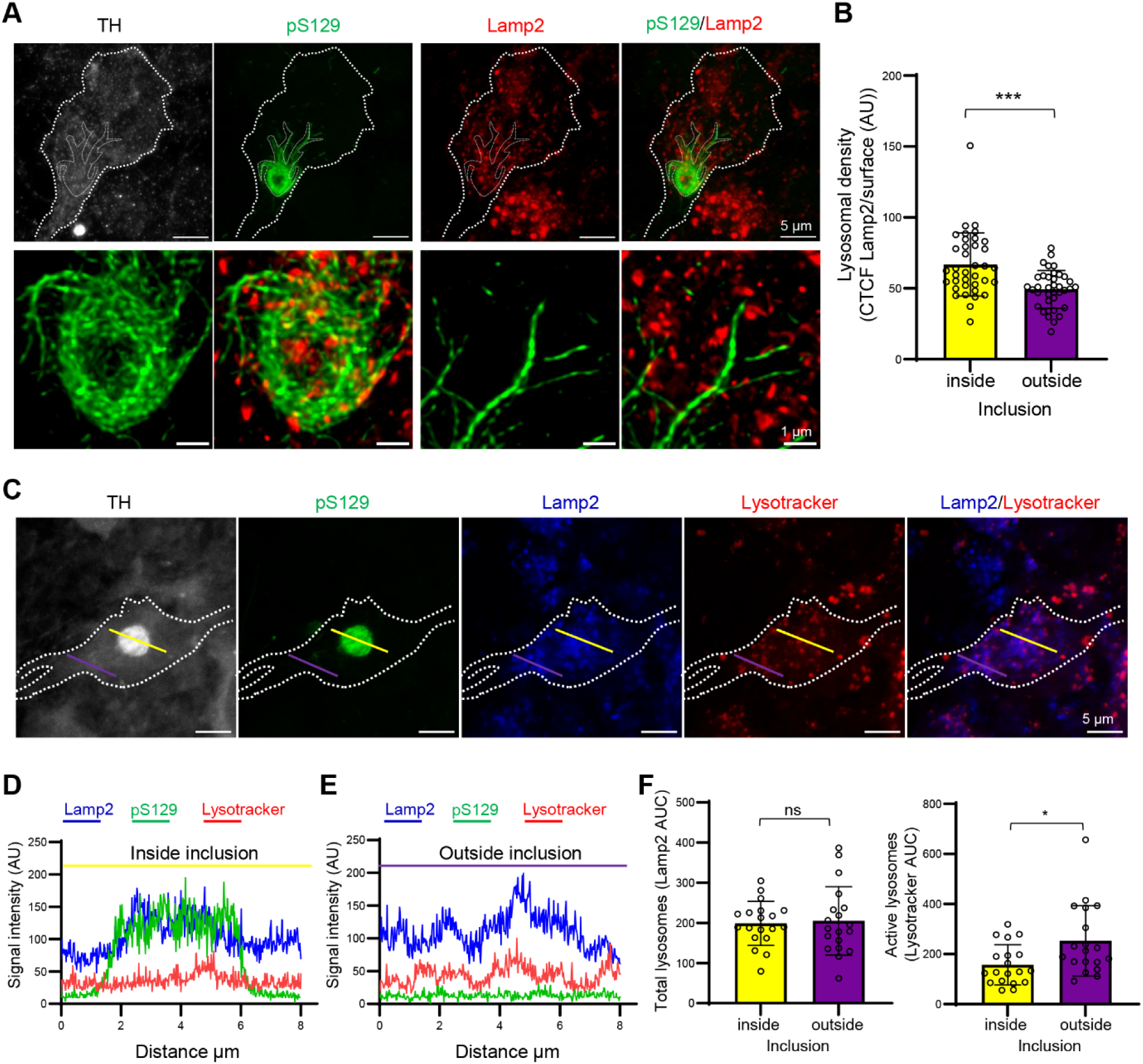
aSyn pathology disrupts lysosomal function in iDA. **A.** Confocal microscopy images (top panel) of iDA neurons (stained for TH, grey), 42 days post-addition of aSyn PFF, showing a radiating aSyn inclusion (stained for pS129, green) highly interacting with lysosomes (stained for LAMP2, red). Scale bar = 5 µm. STED microscopy images (bottom panel) highlight a high accumulation of lysosomes inside and around the core of the inclusion, as well as with the radiating fibers. Scale bar = 1 µm. **B.** Comparison of fluorescence intensity relative to the lysosomes (red) has been measured inside and outside the inclusion (green) for the same TH^+^ neuron to evaluate the spatial distribution of lysosomes inside the cell. Lysosomes are significantly increased inside the inclusions compared to those free in the cytosol (***p<0.0001, unpaired t-test, n=37). **C.** Confocal microscopy images of iDA neurons (stained for TH, grey) showing a dense aSyn inclusion (stained for pS129, green), total lysosomes (stained for LAMP2, blue) and active lysosomes (stained with lysotracker, red). Scale bar = 5 µm. **D.** Fluorescence intensity profiles show that lysosomes (blue) are less active (red) inside the inclusion (yellow line), with few spikes of lysotracker signal compared to LAMP2 (dark arrow). **E.** In comparison, lysosomes outside the inclusion (purple line) show synchronized lysotracker (dark arrows) signals with LAMP2, indicating they are functional. **F.** Quantification of the area under the curve (AUC) for all the lysosomes does not show any significant difference between outside and inside the inclusion, showing that the total level of LAMP2 protein is stable inside and outside the inclusion. The unpaired t-test showed no significant (n.s) difference between the two groups (n = 19). Quantification of AUC for the lysotracker signal shows a significant reduction (*p<0.05, unpaired t-test, n=19) in the fluorescence intensity inside the aSyn inclusion, meaning that the lysosomes inside are less functional.

At D21, filamentous-seeded aggregates were the predominant species (∼60%), with barely any LB-like structures detected (<5%) (Figure 2B, D). By D56, approximately 45% of the seeded aggregates were classified as LB-like inclusions (Figure 2C, E). These findings suggest that aSyn fibrillization is required for LB formation.

Immunostaining showed that TH was absent from early filamentous or speckled aggregates but frequently colocalized with more mature LB-like inclusions (Figures 2 and S7). This relocalization was observed in various inclusion morphologies (Figure S7). In some neurons, TH was diffusely distributed within the inclusion, while in others it was sharply concentrated at the periphery or appeared fully sequestered. Interestingly, where TH was strongly enriched within aggregates, the surrounding cytoplasm often appeared nearly devoid of TH staining, suggesting a progressive sequestration of this enzyme from the cytoplasm into the inclusions over time. Notably, this pattern mirrors observations in the putamen and substantia nigra of patients with PD and MSA, where TH has been shown to accumulate within LB inclusions while its levels in the surrounding cytoplasm are reduced^70^. This suggests that the formation and maturation of LB inclusions may disrupt the production of L-dopa and dopamine by reducing the availability of the rate-limiting enzymes necessary for dopamine synthesis^71-73^. Thus, the presence of TH within pS129-positive aggregates in our model supports the idea that aSyn pathology can disrupt dopaminergic neuron function through altered TH localization and enzymatic activity, in direct agreement with observations from human brain tissue.

In our model, neuritic aSyn pathology consistently appears by D14, preceding the formation of somatic inclusions. Interestingly, we also observed that different types of seeded aggregates within neurons exhibit differential immunoreactivity of key LB pathological markers, including p62, ubiquitin, and β-sheet dyes such as Thioflavin S (ThS), depending on their type and localization (Figures 3 and S8). p62 staining colocalized with all somatic pS129-positive aggregates, regardless of morphology, but not with neuritic pathology (Figures 3A and S8A), consistent with previous reports in mouse neuronal seeding models^40,56,57,60^ and human brain tissues^18^. Conversely, ubiquitin strongly labeled both pS129-positive neuritic aggregates and somatic inclusions, except for the speckled species (Figures 3B and S8B). Neuritic pS129 pathology and mature LB-like inclusions were also strongly positive for ThS, confirming the presence of β-sheet amyloid-like species, while filamentous and speckled aggregates did not colocalize with ThS, although the dye was often found in close proximity (Figures 3C and S8C). Notably, these neuritic aggregates remained p62-negative (Figure 3A), consistent with findings in human PD brain^18^, where such inclusions are considered to reflect an earlier or distinct aggregation state.

This compartment-specific difference suggests that neuritic and somatic aggregates follow distinct molecular pathways or exhibit different kinetics of formation and maturation. Kuusisto et al. first described these differences^18^, and more recently Lewis et al. showed that neuritic aggregates often contain a mix of fibrillar and membranous material^19^, which may provide a microenvironment for initial seeding that later propagates to the soma. Together, these findings indicate that seeded aggregates differ in composition and binding properties, and that amyloid dyes such as ThS may not detect all forms of aSyn pathology, as recently noted by De Giorgi et al. ^74^.

### The iDA seeding model replicates the PTM signature associated with the human pathology

Various post-translational modifications (PTM), including phosphorylation at Serine 129 (pS129) and Serine 87 (pS87), ubiquitination, and nitration (nitration at Tyrosine 39 (nY39) or the C-terminal tyrosine residues (nY125/nY133/nY136)), have been identified in aSyn aggregates and LB within the post-mortem brains of PD or MSA patients^22,23,40^ (Figure 4A). These modifications are believed to influence aSyn aggregation, aSyn fibril interactome and inclusion formation and maturation, as well as toxicity and disease progression^40,56,57,60,75^. Therefore, we sought to determine if the iDA seeding model recapitulates human pathology at the level of PTM.

Toward this goal, we utilized an expanded antibody toolset targeting different sequences within different domains of aSyn and the majority of disease-associated PTM, and it was recently validated in human brain tissues from PD, MSA, and LBD patients^22,23,76^. We analyzed the PTM signatures of aSyn neuritic and somatic pathology that developed over time in PFF-treated iDA neurons. Since only 3% of neurons showed somatic aggregates at D21, we chose to assess PTM signatures at D56, when ∼35% of neurons displayed seeded aggregates encompassing a wide spectrum of morphologies.

Neuritic pathology, which forms early in the aggregation process, consistently exhibited all major disease-associated PTM, including pS129, pS87, pY39, pY125, pY133, and pY136. Strong colocalization of these PTM with pS129-positive neuritic aggregates was observed (Figure S9B), reflecting an early, PTM-rich fibrillar state. This pattern extended to residue-specific nitration, with neurites staining positive for nY39, nY125, and nY136, as well as for total nitrated aSyn (Figure S9C). It is not clear whether these modifications co-occur on the same fibril and what percentage of the fibrils carry these modifications.

By contrast, PTM incorporation in somatic aggregates was highly morphology-dependent. Among the various aggregate types, only LB-like inclusions, either dense or ring-like, showed strong and consistent colocalization with multiple PTM, including pS87, pY125, pY133, pY136, and residue-specific nitration (Figures 4 and S9D-G). In neurons containing both LB-like inclusions and surrounding speckled aggregates, the LB-like structures were PTM-positive while the adjacent speckled pathology remained negative, indicating spatially segregated PTM patterns within the same cell (Figures 4 and S9H-I). In smaller somatic aggregates, such as dots, filamentous, or speckled forms, PTM signals were often restricted to surrounding granules (e.g., pS87 or pY125), rather than fully colocalizing with the aggregates themselves (Figures S9J-M, white arrows). Dot-like and speckled aggregates consistently lacked pY136 staining.

Total nitrated aSyn was detected in most seeded aggregates except filamentous ones; however, only neuritic pathology and LB-like inclusions showed clear labeling with nY39 and nY125/nY136 antibodies. An exception was the LB-like inclusion surrounded by speckled pathology, which also stained positively for nY39. These findings are consistent with recent studies from our laboratory suggesting that nitration may reflect a late-stage modification to neutralize the pathogenic activity of aSyn fibrils and LB^77^.

Altogether, this represents the most comprehensive PTM characterization of seeded aSyn pathology in human-derived neurons to date. Our findings demonstrate that PTM pattern is tightly linked to both the morphology and subcellular localization of aSyn aggregates. Neuritic pathology rapidly accumulates disease-relevant PTM, while somatic aggregates require higher-order organization, specifically LB-like architecture, to exhibit full PTM signatures. These results highlight the complex biochemical landscape of seeded aSyn pathology and support the notion that distinct PTM patterns may reflect maturation stages, structural classes, or divergent aggregation pathways. Whether these modifications precede or follow fibrillization remains to be determined.

### CLEM reveals a large diversity of pS129-positive inclusions at the ultrastructural level

The morphological and biochemical similarities between seeded aggregates in iDA neurons and *bona fide* LB in the human PD brain, including pathological markers, a broad morphological spectrum, and PTM signatures, motivated us to proceed with characterizing the ultrastructure of aSyn pathology formed in iDAs using CLEM. We hypothesized that such significant differences in their biochemical signatures should be reflected in the composition and ultrastructural properties of these different types of aggregates. Given that LB-like aggregates are enriched in membranous structures and organelles, it is crucial to preserve membrane integrity and avoid the damage caused by detergents like Triton. To achieve this goal, we developed a new ICC protocol that eliminates the need for permeabilization (see materials and methods section). In this approach, iDA neurons were fixed in 4% PFA at D21 or D56 and permeabilized by rapid freezing and thawing without detergent, thus preserving the ultrastructure of neurons, inclusions, and organelles. Finally, to validate the presence of aSyn fibrils, they were immunolabeled with a pS129 antibody (81A) (Figures 5-8).

Neurons with pS129 pathology were identified with confocal immunofluorescence imaging, and the same cells were heavy-metal stained, resin-embedded, ultrathin sectioned, and imaged with a transmission electron microscope (TEM) as previously described^40,57,60^. In most inclusions, TEM imaging revealed the presence of fibrils, which colocalized with pS129 aSyn fluorescence immunopositivity (Figures 5-8).

Neuritic aggregates were composed of aSyn filaments and thinner filaments, likely corresponding to neurofilaments, surrounding mitochondria, structures similar to those described by Lam et al^38^. Pre-embedding immunogold labelling confirmed that these structures were indeed aSyn fibrils (Figure 5A-I) rather than cytoskeletal proteins of similar diameter (Figure 5J-K). Interestingly, our data revealed two distinct types of aSyn fibrils, with diameters of ∼7.6 nm and 12.0 nm (Figure 5G-I). ICC also confirmed the absence of colocalization of the pS129 somatic or neuritic pathology with the filamentous cytoskeletal proteins, including the three main subunits of the neurofilaments, namely NF-L, NF-M and NF-H, β-tubulin-III or actin (Figures 5J-L and S10). Interestingly, despite this general lack of association, in this model NF-H recruitment at the periphery of the LB-like inclusion was observed only in dead neurons (Figure 5K, white arrow).

At D21 (Figures 6 and S11), the number and density/distribution of fibrils present in each inclusion varied greatly, with some inclusions predominantly composed of long fibrils.

In contrast, others mainly consisted of small vesicles intermixed with organelles and a sparse number of randomly oriented fibrils. The ultrastructure of a speckled aggregate consisted of long filaments that were organized in a parallel fashion and devoid of membranous organelles (Figure 6A-E). In contrast, the dense inclusions were consistently associated with a mixed ultrastructure of small vesicles, lysosomes, mitochondria and fibrils (Figures 6F-O and S11). When the dense LB was surrounded by speckled pS129 staining, these speckles colocalized with patches of predominantly long, parallel-associated filaments curving through the cytosol and sometimes enclosing the dense aggregate (Figure 6H). Another example of a large, dense LB also consisted of membranes intermixed with fibrils but was surrounded by a thin layer of filaments that presumably belonged to the intermediate filament family (Figure 6M). Interestingly, the condensation of pS129 immunostaining at the periphery of this aggregate correlates with an ultrastructure enriched in mitochondria.

At D56, the CLEM analysis confirmed the ultrastructure of three more dense LB to be a mix of membranous organelles and fibrils, although fewer fibrils and more vesicular structures were present compared to D21 (Figures 7A-O, 8A-B and S12). In total, we identified five ring-like LB-like inclusions and characterized their ultrastructure (Figure S12). The ultrastructure of the pS129-immunopositive halo was found to be heterogeneous: in three examples, it consisted of parallel-associated filaments (Figure 7M), whereas in the remaining two examples, it had an ultrastructure that was rich in small vesicles and sparse fibrils (Figure 7C, H). Using specific antibodies for each organelle, we confirmed that mitochondria, lysosomes, endosomes, the Golgi apparatus, and the endoplasmic reticulum (ER) are differentially sequestered based on the type and subcellular localization (neurites vs. soma) of aSyn-seeded aggregates, indicating distinct organelle involvement at various stages of aggregate formation and maturation within iDA neurons (Figures 7P, 7Q and S13). Despite their ultrastructural heterogeneity (Figure 8B), a common feature of almost all inclusions observed by CLEM was an enriched density of mitochondria at the periphery of the pS129-immunopositive region (Figure 8C), which is consistent with human brain LB pathology.

Mitochondria were consistently associated with nearly all types of aggregates, including neuritic aggregates, filamentous-like aggregates, ring-like LB, dense-like LB, but not with the speckled LB. Lysosomes, on the other hand, were specifically recruited only to LB-like inclusions, such as ring-like LB and dense-like LB. This restricted involvement suggests that lysosomes play a significant role in the processing and maturation of advanced aggregates. Endosomes, however, were found in all aggregate types except neuritic aggregates, suggesting a more specialized role in the progression and processing of more developed aggregates. The Golgi apparatus was consistently recruited to all aggregate types, including neuritic aggregates, filamentous-like aggregates, and all LB-like inclusions. This indicates that Golgi-related processes are implicated across all stages of aggregation, from early to mature inclusions. Although EM clearly revealed the presence of fragmented and damaged ER within LB–like inclusions, the ER was difficult to detect by immunocytochemistry (Figure S13C). BiP showed no colocalization, whereas calreticulin labeling revealed only partial ER signal, suggesting that the limited ICC detection may result from epitope masking or structural disruption of the ER within inclusions. These findings collectively highlight the organelle-specific roles in aSyn aggregate formation and maturation, with mitochondria, lysosomes, the Golgi apparatus, and endosomes contributing distinctively to different stages of the aggregation pathway.

Strikingly, in all five examples, the centre of the ring, where the pS129 immunopositivity is weak, consistently correlated with the presence of electron-dense multilamellar bodies (MLB). Interestingly, these MLB were only observed within LB-like inclusions and were never detected in the surrounding cytoplasm (Figure 8C). These MLB exhibited strong immunoreactivity for LAMP1, a lysosomal marker, confirming their lysosomal identity^78-80^ (Figure 7H-M). These LAMP1-positive structures reflect a specific stage of lysosomal and autophagy failure^78^, where impaired degradation of membranes, organelles, and protein aggregates creates a microenvironment prone to membrane stacking, lipid peroxidation, and progressive MLB formation. In addition, the centre of ring-like inclusions showed enrichment of Tomm20, a mitochondrial outer membrane marker, while intact mitochondria were notably absent on the EM micrographs (Figure 7P). This spatial and molecular profile suggests that MLB may form through lysosomal processing of mitochondrial components. MLB are membrane-bound organelles composed of concentric lamellae, typically associated with lysosomes and known to accumulate when lysosomal degradation is impaired. Their presence within LB-like inclusions, combined with LAMP1 and Tomm20 positivity, points to a convergence of mitochondrial turnover and lysosomal activity under stress conditions. Although such features remain rare in human pathology, dense and lamellated bodies were already described by Forno et al., in axonal swellings associated with LB pathology in Parkinsonism cases^81^, and rosette-like lamellae have occasionally been noted in neuritic inclusions in PD^21^.

In our study, by contrast, these structures were consistently observed within somatic LB-like inclusions. Consistent with this, a recent study demonstrated that PFF-injected mice develop MLB in perinuclear regions of dopaminergic neurons in the SN (similar to Figure 7H-M), where they co-occur with autophagosomes, endolysosomes, and oxidized mitochondria, reflecting active organelle remodeling and stress responses^66^. MLB have also been described in post-mortem Alzheimer’s disease brain tissue^82,83^, where they localize within amyloid plaques and tau tangles and are associated with autophagic stress and impaired proteostasis^78,83^. Together, these observations suggest that MLB within LB-like inclusions represent a pathological interface between mitochondrial degradation and lysosomal dysfunction, marking them as indicators of disrupted organelle homeostasis in aSyn-driven pathology formation and neurodegeneration.

Next, we employed EM to evaluate mitochondrial health in the iDA seeding model. Mitochondria in PFF-treated cells with inclusions, as well as those in PBS-treated control cells, were classified in three groups based on their level of damage (Figure 8D): no / mild damage, moderate damage (light swelling and/or decreased number of cristae) and severe damage (almost complete loss of cristae). In both D21 and D56, the fraction of damaged (moderately and severely) mitochondria was significantly increased in cells with inclusions compared with PBS-treated control cells, indicating that inclusion formation negatively impacts mitochondrial health. Interestingly, the majority of severely damaged mitochondria was found outside and in close proximity to the pS129-immunopositive inclusions, not within (Figure 8D), possibly because severely damaged mitochondria within inclusions might give rise to MLB in the process of inclusion formation.

Altogether, our results at D21 and D56 suggest that the evolving morphological diversity of aSyn aggregates is likely driven by multiple, interconnected or interdependent factors. These include differences in fibril abundance, the spatial distribution and ultrastructural organization of fibrils and membranous organelles within inclusions, as well as variations in their biochemical composition and PTM. In addition, the selective recruitment and sequestration of specific organelles, such as mitochondria, lysosomes, endosomes, and the Golgi apparatus, further contribute to the heterogeneity of aggregate structure and likely reflect stage-dependent remodeling events and cellular stress responses. Together, these elements shape not only the maturation and architecture of aSyn inclusions but also their impact on organelle function and neuronal integrity over time.

### aSyn pathology disrupts lysosomal function in iDA

Previous studies employing advanced techniques such as immunohistochemistry^84^, proteomics^85-87^, STED microscopy^88^, and CLEM^19,21,89,90^ have provided robust evidence that endolysosomal organelles are sequestered within LB inclusions in post-mortem human brain tissues. Proteomic analyses have identified numerous endolysosomal proteins, including those involved in vesicular trafficking and lysosomal function, reinforcing the role of endolysosomal pathways in LB pathology. Lysosomal markers, such as LAMP1, colocalize with aSyn aggregates, while ultrastructural analyses reveal lysosomal vesicles integrated within LB inclusions. These findings suggest a direct sequestration of endolysosomal compartments into LB, indicating their active recruitment into these pathological structures. This sequestration is believed to impair endolysosomal trafficking, resulting in lysosomal dysfunction that leads to cellular stress and disrupted neuronal homeostasis. To determine whether our cellular model also recapitulates this aspect of the pathology, we investigated how the formation and maturation of LB-like inclusions in PFF-seeded iDA neurons impact lysosomal distribution and function.

First, we assessed the extent to which lysosomes were sequestered within LB-like inclusions at D42 (Figure 9A-B). We immunostained lysosomes using LAMP2 and observed their accumulation in contact with, and within, pS129-positive (pS129^+^) aSyn inclusions, suggesting that lysosomes tend to cluster and accumulate in the core of the LB-like inclusion (Figure 9A-B).

To test whether sequestration affected lysosomal function, we used LysoTracker to label active acidic vesicle^25,91,92^ and analysed inside (Figure 9B-C, yellow line) and the fluorescence intensity distribution of LAMP2 and LysoTracker outside (Figure 9B-D, purple line) the pS129+ inclusions. Inside inclusions, LysoTracker did not spike with LAMP2, whereas outside, both signals peaked together (Figure 9C-D). This indicates that lysosomes trapped in inclusions are functionally impaired. Such disruption may reflect an overloaded degradation system but could also represent a protective mechanism by sequestering damaged components. Rupture of lysosomal membranes may then release enzymes that fragment or degrade fibrils, explaining the scarcity of long fibrils in LB cores.

We next quantified fluorescence (AUC) for LAMP2 and LysoTracker across iDA neurons. Outside the inclusions, lysosomes were more active, with LysoTracker intensity peaking simultaneously with LAMP2 (Figure 9D). While total lysosome content (LAMP2 levels) was unchanged, LysoTracker intensity was significantly reduced inside inclusions and elevated outside (Figure 9F). These results indicate that lysosomes sequestered into aSyn inclusions become functionally impaired or inactive.

Consequently, lysosomes accumulate in the core of inclusions but lose activity, resulting in impaired clearance capacity, heightened neuronal stress, and accelerated neurodegeneration. Altogether, these findings underscore the importance of preserving lysosomal integrity and highlight it as a potential therapeutic target in synucleinopathies. Using STED microscopy, we observed a significantly higher lysosomal density within the LB inclusions than in the surrounding cytoplasm, suggesting a concentration of lysosomal vesicles within the inclusions. Moreover, lysosomes sequestered within the LB inclusions exhibited reduced activity, indicating a significant functional impairment (Figure 9C-F). This reduction in lysosomal activity within LB inclusions implies that lysosomal dysfunction may disrupt cellular clearance processes, increasing neuronal stress and contributing to progressive neuronal dysfunction. Such impairment likely accelerates neurodegenerative progression, underscoring the critical role of lysosomal integrity in maintaining neuronal health and highlighting potential therapeutic avenues to target lysosomal preservation in synucleinopathies.

## Discussion

In this study, we employed a human isogenic iPSC-derived dopaminergic neuron iDA model, combined with a PFF seeding approach, to resolve the sequential stages of LB development and maturation in a context that recapitulates human brain pathology without relying on aSyn overexpression or genetic manipulation. We show that this physiologically relevant system captures the morphological, biochemical, and ultrastructural spectrum of aSyn aggregates observed in PD^1,18,19,21,84,93^.

One key advantage of our iDA model is that it consistently generates both neuritic and somatic inclusions, including dense and ring-like LB-like structures. This dual-compartment resolution enables direct, spatially resolved analysis of aggregate formation and progression within the same human neuron system. This is in contrast to most published iPSC-derived seeding model studies, in which filamentous or ribbon-like somatic aggregates, in addition to neuritic pathology, have been observed in only a small subset of studies.^35,36,45,62^

Furthermore, in this model, we observed a wide spectrum of seeded aSyn pathology, ranging from early neuritic filamentous aggregates to dot-like, speckled, and filamentous inclusions, as well as dense and ring-like LB-like inclusions within the soma. This broad phenotypic range mirrors the diversity of Lewy pathology observed in postmortem PD, LBD and MSA brain tissue, and allows the study of both early seeding events and late-stage inclusion maturation within the same human neuron model.

Beyond confirming known features of pathology, the model uncovers new mechanistic insights into the mechanisms underpinning the diversity of aSyn-induced neurodegeneration, pathology formation, and propagation. Using a combination of imaging approaches and molecular profiling, we followed the development of aSyn pathology from early seed uptake to the formation of mature LB-like structures. While relying on pS129 antibodies, which have been the main pathology detection tools in all previous iPSC-derived seeding models, facilitates the detection of aSyn pathological diversity ^25,26,31,34,46-48,62,69^, it allows only limited insight into the biochemical diversity of aSyn aggregates across different inclusions and at different stages of LB formation. To address this, we performed systematic multiparametric immunostaining using additional hallmark markers of human LB pathology, including p62, ubiquitin, ThS, and a broad panel of antibodies against disease-associated PTM and different organelles (Figure S14). This expanded molecular profiling allowed us to define distinct PTM, organelle and ultrastructural signatures associated with specific inclusion morphologies and maturation stages. Altogether, our work shows that LB formation is not the result of a passive accumulation of fibrils, but rather a dynamic, progressive process driven by tightly coordinated events, in which the pattern of aSyn PTM, together with fibril interactions with organelles and other intracellular components, drives the morphological and biochemical heterogeneity of aSyn pathology. Longitudinal analysis at D21 and D56 revealed a reproducible progression from simple filamentous structures to more complex cortical-like, ring-like, and compact LB-like morphologies, accompanied by the accumulation of disease-associated PTM and pathological markers (Figure 10). Although we did not follow individual inclusions over time, the consistency of these patterns supports a stepwise maturation process reminiscent of LB development in vivo.

**Figure 10.**
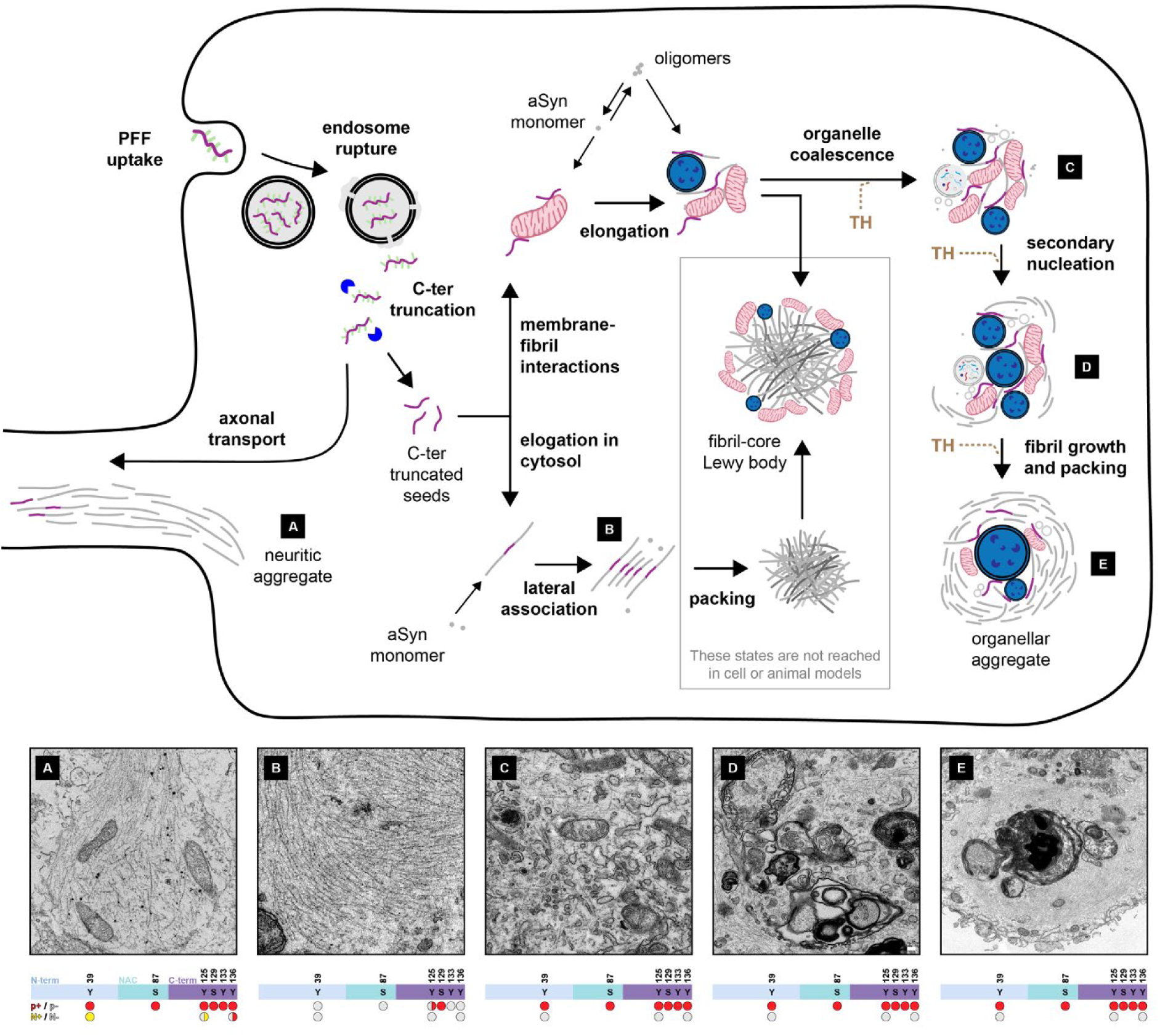
Mechanistic model of aSyn aggregation and inclusion diversity. Schematic representation of aSyn pathology formation, integrating data from this study with prior seeding models and recent EM analyses of LB from human postmortem brains in PD and LBD. Following internalization into iDA neurons, PFF undergo rapid C-terminal truncation, predominantly at residue 114. These cleaved seeds recruit endogenous monomeric aSyn, initiating fibril elongation and amplification. Two distinct pathways are proposed: a membrane/organelle-independent (**A-B**) and a membrane/organelle-dependent mechanism (**C-E**). The contribution of each pathway likely depends on local monomer availability and cellular context. In the membrane-independent route, fibrils elongate in the cytosol via monomer addition, undergo specific PTM, and may cluster into dense fibrillar assemblies. Neuritic aggregates **(A)** are predominantly formed this way, after transport of truncated seeds to neurites. In the soma, when fibril growth outpaces organelle recruitment, compact inclusions with laterally associated fibrils are formed. These purely fibrillar inclusions might be precursors of more densely packed fibril-core Lewy bodies with radiating fibrils and peripheral organelles. However, this is speculative given that inclusions with radiating fibrils have, to our knowledge, not been observed in seeding models. Conversely, in the membrane-dependent route (**C-E**), truncated seeds associate directly with intracellular membranes, where fibrils elongate and incorporate lipids. These lipid-rich fibrils can scaffold organelle accumulation, resulting in membrane- and organelle-dense inclusions. Variability in fibril morphology, PTM, packing, and organelle content likely reflects differences in seeding dynamics, maturation states, aberrant processing, and incorporation of tyrosine hydroxylase (TH) into the inclusion. Notably, fibrils within mature inclusions are consistently shorter than those observed at earlier time points or peripherally, suggesting fragmentation by organelle-derived proteases. While this may support degradation, impaired clearance could allow accumulation of seeding-competent fragments, promoting pathology propagation. LB-like inclusions may thus serve as reservoirs of such species under conditions of proteostatic failure. This model captures seeded fibril amplification and inclusion evolution but does not encompass early-stage oligomerization, which may represent parallel or precursor pathways. Further models bridging the full aggregation continuum, from oligomers to mature inclusions, will be critical to understanding the pathogenesis of synucleinopathies.

In this context, it is informative to consider prior reports of LB-like pathology in iPSC-derived neurons. To date, only two studies have described such structures following PFF treatment. Lam et al. used iPSC-derived cortical neurons expressing aSyn-GFP to study the ultrastructure of somatic inclusions^38^. These inclusions contained fibrils, clustered vesicles, lipid droplets, dysmorphic mitochondria, and filamentous material, features reminiscent of human LB. The authors classified multiple inclusion types, including neuroprotective p62-positive structures and lipid-rich, neurotoxic inclusions. While the use of aSyn-GFP fusion constructs facilitates the detection of aSyn aggregates and monitoring their fate, it is also known to alter fibril surface properties and cellular interactions, potentially affecting inclusion morphology and composition^94-96^. In previous neuronal seeding models expressing aSyn-GFP fusion constructs, the inclusions were found to be composed primarily of fibrillar aggregates^97^.

In parallel, Bayati et al. demonstrated that LB-like inclusions could form in iPSC-derived dopaminergic neurons treated with aSyn PFF, but only under immune challenge with interferon-γ^26^. In their model, lysosomal impairment facilitated PFF uptake and aggregate accumulation within membrane-bound compartments, proposed to result from autophagosome expansion due to chaperone-mediated autophagy (CMA) dysfunction^26,98^. However, these membrane-enclosed inclusions contrast with the membrane-less LB observed in human tissue^21,81,99-103^, except in rare neuritic cases^19^. These membrane-enclosed inclusions were identified primarily through immunogold labeling of the PFF, without molecular or ultrastructural confirmation of their correspondence to authentic LB pathology.

### PTM shape the biochemical architecture and identity of aSyn pathology

Our iDA seeding model reveals that different types of aSyn PTM are selectively enriched across distinct aggregate subtypes and neuronal compartments. Moreover, our results reveal that neuritic and somatic aSyn pathology exhibit distinct aSyn PTM patterns. Mature LB-like inclusions displayed the most complex PTM signatures, whereas early-stage or filamentous aggregates showed minimal modification. This suggests either a progressive accumulation of PTM during aSyn aggregate formation and maturation, or that specific PTM patterns may influence the kinetics of pathology formation and the types of aggregates that emerge. Interestingly, pS129 was the only PTM consistently detected across all aggregate morphologies and compartments, including dot-like, speckled, filamentous, neuritic, and LB-like structures. This positions pS129 as a central hallmark of fibrillar aSyn pathology and the most reliable marker for capturing the full spectrum of seeded aggregates (excluding non-fibrillar oligomers). Its ubiquitous presence suggests that pS129 is deposited early, potentially during or immediately after fibrillization, while other PTM appear selectively and correlate more strongly with inclusion maturation and subcellular context.

Previous STED microscopy of postmortem human PD tissue has shown that PTM-modified aSyn species, such as pS129 and N- and C-terminal truncations, form concentric, layered “onion-skin” architectures within inclusions^88^. This structural stratification suggests that PTM are not merely post hoc consequences of aggregation but may actively regulate the progression of inclusion assembly. These patterns imply the existence of distinct biochemical microenvironments within aggregates, potentially governed by local kinase/phosphatase activity, protein quality control mechanisms, or oxidative stress. In line with this, prior studies have found that early-stage aggregates are enriched in PTM, such as pS129 and nY39, while later-emerging modifications, such as pS87 and pY136, are associated with more mature inclusions^104^. Our data mirror this progression, with complex PTM profiles accumulating in compact, late-stage aggregates and early inclusions remaining largely unmodified (Figure 10). The association of multiple PTM with different types of aSyn pathology presents challenges to elucidating their relative contribution to modulating aSyn pathology formation and pathogenicity. It suggests that crosstalk between these PTM should be further investigated and accounted for when developing therapeutic strategies targeting aSyn PTM. Recent studies from our group demonstrate that PTM can also modify the biochemical behavior and pathogenic potential of aSyn aggregates^40,75,77^, with modifications such as O-GlcNAc modification or nitration nearly abolishing the seeding activity of aSyn fibrils in vivo^75,77^.

Neuritic aggregates, which appear early in the aggregation process, already exhibit a full panel of disease-relevant PTM, including pS87, pY125, pY133, pY136, and nitration at Y39, Y125, and Y136, alongside pS129. The early and broad incorporation of PTM in neuritic aggregates suggests that they undergo biochemical maturation soon after their formation, potentially reflecting an environment that favors early modification. In contrast, the majority of somatic aggregates remain largely PTM-negative despite being strongly positive for pS129, except for dense or ring-like LB-like inclusions, which consistently show strong PTM colocalization. Early-stage aggregates, such as filamentous, dot-like, or speckled structures, displayed minimal PTM labeling, indicating either early-phase aggregates prior to PTM accumulation or inclusion subtypes with reduced modification capacity. Interestingly, in neurons that simultaneously contain LB-like inclusions and adjacent speckled aggregates, only the LB-like structures are PTM-positive, while speckled aggregates remain negative, demonstrating selective PTM incorporation within the same cellular environment. These findings support the idea that PTM profiles not only reflect the stage of aggregate maturation but also vary by subcellular localization and possibly by the structural identity of the aggregate itself. These compartment-specific PTM profiles likely have functional consequences.

Interestingly, pS129 was the only PTM consistently detected across all aggregate morphologies and compartments, including dot-like, speckled, filamentous, neuritic, and LB-like structures. This positions pS129 as a central hallmark of fibrillar aSyn pathology and the most reliable marker for capturing the full spectrum of seeded aggregates (excluding non-fibrillar oligomers). Its ubiquitous presence suggests that pS129 is deposited early, potentially during or immediately after fibrillization, while other PTM appear selectively and correlate more strongly with inclusion maturation and subcellular context. This distinction underscores the utility of pS129 for broad aggregate detection, but also highlights its limitations in resolving aggregate heterogeneity or pathogenic potential. Altogether, these observations suggest that aSyn aggregates can exist in at least two biochemical states: pS129-positive/PTM-negative and pS129-positive/PTM-positive (Figure 10). This dichotomy reflects compartment-specific processing and raises the possibility that distinct cellular environments shape PTM accessibility and dynamics. Whether neuritic pathology represents an independent route of pathology development or an early stage in a continuum that culminates in somatic LB formation remains unclear. It is well established that neuritic pathology can extend retrogradely toward the soma; however, our findings raise the question of whether full PTM acquisition occurs only upon inclusion maturation within the somatic compartment. One possibility is that, as LB evolve and begin to sequester organelles, they entrap the key modifying enzymes required for PTM deposition, creating localized conditions that facilitate further biochemical remodeling.

These ultrastructural observations raise a broader question in synucleinopathy research: whether LB serve primarily harmful or protective functions. Our findings support a dual perspective. On one hand, the accumulation of mitochondria, lysosomes, and endosomes within inclusions coincides with structural disruption and loss of organelle function, most clearly demonstrated by our lysosomal functional assays. Lysosomes trapped inside inclusions exhibit significantly reduced acidification and functional impairment, suggesting that their sequestration compromises degradation capacity and may exacerbate cellular stress. Likewise, mitochondrial damage was more pronounced in neurons with inclusions, with a subset of severely damaged mitochondria observed outside of aggregates, potentially representing organelles that had not yet been engulfed or processed into MLB.

On the other hand, the selective compartmentalization of dysfunctional organelles into inclusions could reflect a protective mechanism aimed at shielding the surrounding cytosol from further harm. This aligns with models proposing that LB function to sequester potentially harmful or damaged cellular components. However, as our data show, this sequestration comes at a cost: lysosomal function is compromised, organelle turnover is disrupted, and essential proteins and enzymes like TH become mislocalized and likely rendered non-functional due to their confinement within inclusions and reduced accessibility within the cytoplasm.

### A mechanistic model of aSyn pathology formation and diversity

On the basis of our observations, we propose a working model for the mechanisms of aSyn seed–mediated pathology formation that could partially explain the pathological diversity observed in human PD brains (Figure 10). Upon internalization of preformed fibril seeds into neurons, through the endolysosomal pathway, they undergo rapid cleavage of the last 25 C-terminal residues, which increases the hydrophobicity of the fibril surfaces and facilitates both primary and secondary nucleation events at the fibril ends and surfaces, respectively. At this stage, aSyn aggregation can proceed through at least two pathways: a membrane-independent mechanism and a membrane-dependent mechanism.

#### Membrane-independent pathway

Fibril elongation through monomer addition, combined with PTM incorporation and further C-terminal cleavages, generates fibrils with a strong propensity for lateral association and rapid clumping into dense fibrillar assemblies. These assemblies are distinguished by cores enriched in truncated aSyn species and peripheries enriched in full-length and phosphorylated aSyn. Membranous organelles, such as mitochondria, are subsequently recruited to the periphery. Notably, this process occurs in both neurites and soma; however, the PTM patterns of neuritic fibrils differ from those of organelle-free fibrillar accumulations in the soma, suggesting compartment-specific interactomes and differential fibril processing.

#### Membrane-dependent pathway

Growing evidence indicates that, upon internalization into neurons, aSyn fibrils associate with various membranous organelles, including mitochondria. Fibril elongation on organelle surfaces promotes the formation of longer fibrils that may act as scaffolds for the recruitment of additional membranous organelles. Thus, it is plausible that fibril interactions with different organelles may trigger distinct endogenous fibril seeds and pathways to fibrillization and LB formation.

Our CLEM studies suggest that organelle sequestration in the vicinity and center of LB is not a passive consequence of aggregation but appears highly selective and compartment-specific (Figure 10). Mitochondria were consistently recruited across inclusion types, but their distribution varied: they often accumulated at the periphery of pS129-positive inclusions, a pattern also reported in PD patient brain tissue^21^. Lysosomes, in contrast, were found exclusively in more mature, LB-like aggregates, implying that lysosomal engagement is a late-stage event, potentially reflecting an overwhelmed or dysfunctional autophagic response. The Golgi apparatus and endosomes were recruited more broadly, including to earlier-stage aggregates, suggesting that secretory and trafficking pathways may be disrupted from the onset of seeded pathology. A particularly compelling feature of mature LB inclusions is the presence of electron-dense MLB in their cores, consistent with previous reports in Alzheimer’s disease^82^ and PD models^26,66^ where they arise under sustained mitochondrial and lysosomal stress^26,66,79,80^. Their selective restriction to the inclusion core suggests that MLB represent a late-stage hallmark of LB maturation, reflecting combined mitochondrial and lysosomal dysfunctions. Taken together, we propose that the differential recruitment and fate of organelles contribute to the distinct morphologies of aSyn pathological aggregates observed across disease contexts (Figure 10).

Moreover, it is plausible that fibrils formed through the membrane-dependent pathway differ in structure, biochemical composition (PTM and lipids), and secondary nucleation properties from those formed via the membrane-independent pathway. Previous studies have shown that fibril growth on membrane surfaces can disrupt membranes and sequester lipid molecules, which then become incorporated into the fibrils. Lipid incorporation could alter fibril structure and biochemical properties, potentially explaining why speckled-like aggregates are devoid of PTM (e.g., ubiquitination) and do not bind ThS. A consistent observation across our CLEM studies is that fibrils located in the core of membranous organelle–enriched inclusions are markedly shorter than those at the periphery of inclusions or in organelle-free neuritic and somatic fibril bundles. This suggests that core fibrils may undergo fragmentation driven by proteolytic enzymes released from disrupted lysosomes. Since fibril size likely influences their ability to be secreted and internalized, and smaller fibrils exhibit higher internalization and seeding activity, these findings further link LB formation to the generation of seeding-competent small fibrils. Altogether, this supports the idea that blocking the transition from fibrils to LB could hinder the propagation of aSyn pathology.

Taken together, our mechanistic model (Figure 10) provides a framework for testing how different pathways of fibril growth, remodeling, and fragmentation contribute to the diversity of aSyn pathology. This framework can be refined and expanded through future studies that integrate high-resolution imaging, biochemical analysis, and structural characterization of both model-derived and human brain–derived fibrils. In this context, it will be particularly important to address how the exposed N- and C-terminal domains of fibrils regulate interactions with cellular proteins and organelles, and how PTM within these domains influence pathology formation and spreading. Developing new neuronal models that express, at physiological levels, N- and C-terminal variants designed to alter fibril–membrane or fibril–organelle interactions could provide critical insights into the mechanisms that shape aSyn aggregation, inclusion maturation, and toxicity.

### Conclusion

Our study establishes the iDA seeding model as a physiologically relevant human neuron system that resolves the sequential stages of Lewy pathology formation without genetic manipulation or aSyn overexpression. In this model, inclusions emerge through progressive incorporation of PTM, selective recruitment of mitochondria, lysosomes, and other organelles, and the late appearance of multilamellar bodies, together defining a stepwise maturation process. These observations indicate that inclusion remodeling is an active process that can both compartmentalize damaged components and, at the same time, compromise essential cellular functions. By capturing these dynamic and compartment-specific events, the iDA model provides a framework for dissecting the mechanisms that shape aSyn pathology and for testing targeted approaches to modulate its maturation and impact on neuronal health.

## Material and Methods

### Purification of human WT aSyn

Recombinant overexpression and purification of human WT aSyn were carried out using E. coli BL21 (DE3) cells transformed with pT7-7 plasmids encoding WT aSyn. After growing the cells in Luria broth (LB) medium with ampicillin, protein expression was induced with 1 mM 1-thio-β-d-galactopyranoside when the optical density at 600 nm reached 0.4–0.6. The culture was incubated for 4–5 hours before the cells were harvested by centrifugation. Cell pellets were lysed using ultrasonication, and the lysate was subjected to heat treatment followed by centrifugation to remove impurities. The resulting supernatant was purified using a HiPrep Q Fast Flow column, and fractions containing pure aSyn were analyzed by SDS-PAGE, pooled, and further purified using reverse-phase HPLC as previously described^105^. The final product was confirmed to be highly pure through UPLC and ESI-MS, then snap-frozen and lyophilized for storage.

### Preparation of WT aSyn monomers

The lyophilized WT aSyn was resuspended in PBS buffer and processed according to a previously described filtration method^105^. The resuspended aSyn was adjusted to a pH of approximately 7.2–7.4 and filtered using 100 kDa spin filters. The filtrate containing the monomeric form of aSyn was quantified by measuring the absorbance at 280 nm using a Nanodrop 2000 Spectrophotometer (ThermoFisher) and an extinction coefficient of 5960 M−1 cm−1, as predicted from the aSyn sequence (ProtParam, ExPASy). Monomeric WT aSyn was prepared for use in subsequent experiments.

### Preparation of WT aSyn PFF

WT aSyn PFF were prepared by dissolving 4 mg of lyophilized recombinant aSyn in 600 μL of 1x PBS^105^. The pH was adjusted to approximately 7.2–7.4, and the solution was filtered through 0.2 μm filters (MERCK, SLGP033RS). The filtrate was transferred to black screw-cap tubes, which were incubated at 37 °C and shaken at 1000 rpm for five days. Fibril formation was verified through transmission electron microscopy (TEM) and Coomassie staining, as previously described^105^. For sonicated seed preparation, WT aSyn PFF were sonicated for 20 seconds at 20% amplitude, with 1-second pulses on and off (Sonic Vibra Cell, Blanc Labo, Switzerland). Following established protocols, the amount of monomers and oligomers released from the sonicated fibrils was quantified through filtration. TEM confirmed the fibril structure, and Coomassie staining was used to quantify the release of monomers and oligomers following sonication^105^.

### Preparation of WT aSyn PFF fluorescently labelled

Fluorescent labelling of human WT aSyn PFF was performed as previously described^60^ by diluting PFF to a concentration of 250 μM in 500 μL of PBS, with the pH adjusted to 7.5. An equivalent amount of Atto488 or Atto647 maleimide (Atto-Tec, Switzerland) was added, and the mixture was incubated overnight at 4 °C. After incubation, the labelled PFF were ultracentrifuged at 100,000 g for 1 hour at 4 °C. The supernatant was discarded, and the pellet was resuspended in PBS, repeating the wash steps until the dye in excess was removed. Labelling was confirmed by running the fibrils on SDS-PAGE and scanning the gel using the Typhoon FLA 7000 (GE Healthcare) with excitation and emission wavelengths of 400 and 505 nm, respectively. The labelled fibrils were fragmented through sonication, using four cycles of 5 seconds at 20% amplitude (Sonic Vibra Cell, Blanc Labo, Switzerland). Finally, the PFF were snap-frozen in liquid nitrogen and stored at −80 °C. The structural and biophysical properties of the fibrils were evaluated using electron microscopy, SDS-PAGE with Coomassie staining, and thioflavin T (ThT) assays.

### Differentiation of NGN2-iPSCs into dopaminergic neuronal culture (iDA)

The optimized protocol for generating functional iDA cultures has been fully described by Sheta et al. ^54,55^. Briefly, the NGN2-expressing human iPSCs (AIW002-02, from The Early Drug Discovery Unit (EDDU), McGill University, Canada) were first differentiated into iNeurons. On day −1, cells were plated in mTESR plus media (STEMCELL Technologies, Switzerland) containing Y27632 (Tocris, Switzerland). The next day (DIV0), the mTESR plus media was fully replaced by the complete DIV0/DIV1 media composed of DMEM/F12 (Life Technologies, Switzerland), N2 (Life Technologies, Switzerland), B27 (Life Technologies, Switzerland), NEAA (Life Technologies, Switzerland), BDNF (10ng/mL) (450-02; Peprotech, United Kingdom), GDNF (10ng/mL) (450-10; Peprotech, United Kingdom), mouse laminin (23017015; Life Technologies, Switzerland), and doxycycline (2 µg/mL) (Sigma-Aldrich, Switzerland) in order to induce NGN2 expression. On DIV 1, the media was fully changed again, using the complete DIV0/DIV1 media as described above. At DIV2, the cells were dissociated with Accutase and replated on dishes (96-well or 48-well black clear bottom plates) or on coverslips coated with poly-L-ornithine (Sigma-Aldrich, Switzerland) and laminin (23017015; Life Technologies, Switzerland) in a Neurobasal medium supplemented with N2, B27, GlutaMax, NEAA, BDNF (10ng/mL), GDNF (10ng/mL), mouse laminin and doxycycline (2 µg/mL). The day after (DIV3), the NGN2-expressing iNeurons were exposed to differentiation media from the STEMdiff Midbrain Neuron Differentiation Kit (STEMCELL Technologies, Switzerland), supplemented with doxycycline (2 µg/mL) and Sonic Hedgehog (200 ng/mL). At DIV 6, the media was fully changed with the complete STEMdiff Midbrain Neuron Differentiation supplemented with doxycycline (2 µg/mL) and Sonic Hedgehog (200 ng/mL). At DIV 9, the media was replaced with maturation media from the STEMdiff Midbrain Neuron Maturation Kit (STEMCELL Technologies, Switzerland). From DIV9 onward, half of the media was replaced once a week with fresh maturation media.

### PFF treatment

The iDA cultures were plated in 35 mm fluorodishes (Millian, France), either with or without coverslips (CS) (VWR, Switzerland), at a density of 500,000 cells/mL. iDA cultures were plated in black, clear-bottom 96-well plates (Falcon, Switzerland) at a density of 100,000 cells/mL. On DIV9, the differentiation media was completely replaced with maturation media. This media either contained aSyn PFF at a final concentration of 500 nM, or an equivalent volume of PBS, which matched the amount of PFF added to the extracellular media.

| **PFF treatment**<br>(500 nM final) | **Cell density (cells/mL)** | **Final volume of media per well** |
| --- | --- | --- |
| 35 mm fluorodishes | 500,000 | 1.5 mL |
| Black/clear flat-bottom 96-well plates | 100,000 | 100 μL |

### Hippocampal primary neuronal culture

Pregnant C57BL/6Jrj mice (Janvier Labs, France) were imported on demand. Neonatal mice (postnatal days 0–1) were euthanized by decapitation immediately prior to brain dissection and primary hippocampal neurons were isolated following established protocols^57^. All procedures were performed ex vivo. In alignment with the 3Rs principle, dams were, when appropriate, redirected to the EPFL Organ/Tissue Sharing Program (Optimice). All animal protocols received approval from the Swiss Federal Veterinary Office (authorization numbers VD 3392 and VD 4029). Cells were plated in 6-well plates pre-coated with 0.1% (w/v) poly-L-lysine in water (Brunschwig, Switzerland), at a density of 300,000 cells/mL.

### Total cell lysis and WB analyses

iDA cultures were lysed using 2% SDS in Tris-buffered saline (TBS) (50 mM Tris, 150 mM NaCl, pH 7.5) containing the protease inhibitor cocktail (Roche, Switzerland), 1 mM phenylmethane sulfonyl fluoride (PMSF), and phosphatase inhibitor cocktails 2 and 3 (Sigma-Aldrich, Switzerland). The cell lysates were boiled for 10 minutes, and protein concentration was determined using a BCA protein assay (Thermo Fisher Scientific, Switzerland). Laemmli buffer (4% SDS, 40% glycerol, 0.05% bromophenol blue, 0.252 M Tris-HCl pH 6.8, and 5% β-mercaptoethanol) was added to the lysates, and 30 μg of total protein was loaded onto 16% tricine gels for 2 hours at 125 V. Proteins were then transferred to 0.2 μm nitrocellulose membranes (GE Healthcare, Switzerland) using a semidry transfer system (Life Technologies, Switzerland) at 25 V and 0.5 A. Membranes were blocked for 1 hour at room temperature (RT) in Odyssey blocking buffer (Li-COR Biosciences, Germany), followed by overnight (O/N) incubation at 4°C with primary antibodies (listed in the tables in Figure S1). After three washes in PBS with 0.1% Tween 20 (PBS-T), the membranes were incubated with secondary antibodies (Alexa Fluor 680 or 800, Li-COR Biosciences). The membranes were washed again with PBS-T and scanned using a Li-COR scanner (Li-COR Biosciences, Germany). Total aSyn levels were quantified by analyzing the WB band intensity using Image Studio software (RRID:SCR_015795, Li-COR Biosciences, Germany) and normalized to actin levels. All experiments were performed in triplicate independently.

### Immunocytochemistry (ICC)

iDA cultures were fixed with 4% formaldehyde (Sigma-Aldrich, Switzerland) for 20 minutes at RT, followed by immunostaining as previously described^106^. Details regarding the antibodies used, including their source and dilution, are provided in each figure legend and Figure S1. In brief, the iDA cultures were blocked with 3% bovine serum albumin (BSA, Sigma-Aldrich, Switzerland) in PBS containing 0.1% Triton X-100 (PBS-T) for 30 minutes at RT. After blocking, the cells were incubated overnight at 4 °C with primary antibodies. Following five washes in PBS-T, the cultures were incubated with secondary antibodies and stained with DAPI (Life Technologies, Switzerland). The cells were again washed five times with PBS-T before being mounted using polyvinyl alcohol mounting medium with DABCO (Sigma-Aldrich, Switzerland). Cells plated on coverslips were examined using a confocal laser scanning microscope (LSM 700, Carl Zeiss) with a 40x objective, and images were analyzed with Zen software (RRID:SCR_013672). For iDA cultures grown in black, clear-bottom 96-well plates, imaging was performed using the IN Cell Analyzer 2200, as previously described^57^.

### Assessment of PFF uptake in iDA neurons by live cell imaging

At DIV10, iDA neurons were stained with 1 µM LysoTracker and NeuroFluor NeuO (StemCell, cat. no. 01801) diluted 1:400 in BrainPhys Imaging Optimized Medium (StemCell, cat. no. 05796) for 45 minutes. Following staining, the medium was replaced with fresh BrainPhys Imaging Optimized Medium containing 300 nM of labelled PFF^647^). Cells were placed in a stage-top incubator (Tokai Hit) on the microscope, with CO₂ maintained at 5% and temperature at 37°C. After 20 minutes of incubation, time-lapse imaging was initiated using an inverted Leica TCS SP8 STED 3X microscope in confocal xyzt mode, equipped as previously described, with the following modifications: HC PL APO CS2 63x/1.40 OIL objective, 4.5x numerical zoom, 1024x1024 pixel resolution, bidirectional scanning at 1400 Hz with 3-line averaging. Two sequential z-stacks were acquired with 1 µm steps (5 µm total thickness) to capture all images for both sequences within 17.389 seconds (8.69 seconds per stack for each sequence). Continuous imaging was performed after each stack acquisition for up to 10 minutes. Fluorescence was recorded using HyD detectors with these settings: First sequence (dual-color acquisition): NeuO (3% laser power at 480 nm, fluorescence collected at 485-550 nm, gain 80V, 0-3.50 ns time gating) and PFF-AF647 (3% laser power at 645 nm, fluorescence collected at 651-720 nm, gain 130V, 0-3.50 ns time gating). Second sequence (single-color acquisition): LysoTracker (2.5% laser power at 570 nm, fluorescence collected at 578-650 nm, gain 80V, 0-3.50 ns time gating).

### Quantification of total aSyn endogenous level in iDA cultures at DIV9

iDA cultures were fixed at DIV9 in 4% formaldehyde (Sigma-Aldrich, Switzerland) for 20 minutes at RT, immunostained as described in the section ICC above and acquired using IN CellAnalyser 2200 (GE Healthcare). For each independent experiment, three wells were acquired per condition, and in each well, nine fields of view were imaged. Each independent experiment was reproduced at least 3 times. The identification of TH-positive or negative neuronal cells and the quantification of aSyn intensity were performed using a custom-built Macro in the software Fiji^107^. In brief, cells were thresholded based on their TH fluorescence signal, and nuclei were thresholded based on DAPI staining. Then, cells were separated from each other using a marker-controlled watershed with the centre of the nuclear detections as seeding points and the binarized TH channel as a mask. Cells with incorrectly differentiated neurites were manually excluded from the analysis. Within each cell, the median gray value of the aSyn fluorescence channel was used as a representative measure of the aSyn level in that cell. Mean fluorescence intensity per cell was divided by the corresponding cell area and normalized to the population mean of each independent experiment, then visualized as a violin plot.

### Correlative Light Electron Microscopy (CLEM)

iDA cultures were grown in 35 mm fluorodishes (Millian, France) with manually engraved alphanumeric grids etched into the bottom glass. The cells were treated with either PBS (as a negative control) or 500 nM of human WT PFF or PFF^488^. At the specified time point, cells were fixed for 2 hours using 2.0% paraformaldehyde (PFA) in 0.1 M phosphate buffer (PB) at pH 7.4. Instead of using detergent for permeabilization, a freeze-thaw method was employed to preserve cellular membranes and organelles better. Briefly, cells were incubated for 20 minutes in a cryoprotectant solution (20% DMSO and 2% glycerol in 0.1 M phosphate buffer, pH 7.4). Then, each dish was filled with liquid nitrogen (LN2) for about 7 seconds; the LN2 was removed, and cryoprotectant was added immediately afterward. This freeze-thaw process was repeated twice. After washing with PBS, ICC was performed (refer to the Materials and Methods ICC section for more details). TH-positive neurons containing LB-like inclusions (positive for pS129 staining) were selected using a confocal fluorescence microscope (LSM700, Carl Zeiss, Germany) for ultrastructural analysis. The precise location of selected cells was recorded using the alphanumeric grid on the dish. Cells were then processed for electron microscopy following a previously established protocol^57^. Briefly, cells were fixed again with 2.5% glutaraldehyde and 2.0% paraformaldehyde in 0.1 M PB at pH 7.4 for an additional 2 hours, post-fixed in 1% osmium tetroxide, stained with uranyl acetate, dehydrated through a graded ethanol series, and embedded in Durcupan resin. After polymerization, the resin-embedded cells were trimmed, sectioned (50–60 nm), and mounted on copper grids for ultrastructural analysis. Sections were contrasted with uranyl acetate and lead citrate, and imaged using a Tecnai Spirit transmission electron microscope (FEI) operated at 80 kV.

#### Immunogold staining

To validate the presence of fibrillar alpha-synuclein within aggregates, pre-embedding immunogold labeling was performed. After PFA-fixation and incubation in primary antibodies (see above), cells were incubated with FluoroNanogold conjugated secondary antibody (Nanoprobes) for 2 hours at RT, and immunofluorescence microscopy, resin embedding, ultramicrotome sectioning and TEM imaging were performed as described above. On the resulting EM micrographs, gold particles were detected using a difference-of-gaussian filter implemented in FIJI^107^.

#### Tomography

To measure fibril properties, tilt series from –60 to 60 degrees were acquired on resin-embedded slices using a transmission electron microscope (Tecnai T12, FEI) at 120 kV. Tomograms were reconstructed using IMOD software^108^. Small training sets of gold-associated, thick filaments (class 1) and thin filaments without gold particles attached to it (class 2) were segmented manually, and two convolutional neural networks were trained in EMAN2^109^ to segment both classes separately in the whole tomogram. In FIJI, the thickness of the fibrils was measured by making a 3D distance map of the fibril segmentations which was evaluated at the pixels that constitute the skeletonization of the segmentations.

#### Super-resolution STED imaging

For super-resolution STED microscopy, iDA neurons on coverslips were fixed 52 days post-PFF exposure using 3% PFA and 0.1% glutaraldehyde for 15 min at RT. After fixation, neurons were washed three times in PBS (5 min each) and permeabilized with 0.3% Triton X-100 (Sigma-Aldrich, Canada) for 5 min at RT. Following a PBS wash, neurons were incubated in blocking solution (5% normal goat serum, 1% BSA, 0.1% Triton X-100 in PBS) for 1 h, then with primary antibodies in the blocking solution overnight at 4°C (refer to antibody table). The next day, neurons were washed three times with blocking solution (5 minutes each) and incubated with secondary antibodies for 1 hour (STAR635P for LAMP2 and AF594 for pS129; see the antibody table). After three additional PBS washes (each 5 min) and a brief rinse in Milli-Q water, coverslips were mounted using Prolong Diamond Antifade reagent (refractive index 1.47; Thermo Fisher, Canada).

Imaging was performed on an inverted Leica TCS SP8 STED 3X microscope, featuring a motorized stage, a tunable white-light laser (470-780 nm), and a 405-nm diode laser (Leica Biosystems, Concord, ON, Canada). Imaging parameters are reported following the guidelines provided by Llopis et al.^110^. Sequential scanning of each channel was done with a 405-nm laser (TH, Alexa Fluor 405), 590-nm laser (pS129, Alexa Fluor 594), or 656-nm laser (LAMP2, Abberior STAR 635P). Fluorescence depletion for Alexa Fluor 594 (pS129) and Abberior STAR 635P (LAMP2) was achieved using a 775-nm pulsed laser set at 30% input power, with tuning at 9% and 4.5% for pS129 and LAMP2, respectively. Laser intensities and gain settings were optimized to maximize the signal-to-noise ratio and prevent saturation using the QLUT Glow range indicator. Point scanning was conducted using a bidirectional scanner set to 700 Hz, with 4-line averaging applied. A 100x HC PL APO CS2/1.40 NA STED white objective (Leica, Canada) with immersion oil type F (refractive index 1.5180, Leica, Canada) and a 2x numerical zoom was used. Hybrid detectors (HyD) collected the emitted wavelengths: 655-750 nm for LAMP2 (Abberior STAR 635P), 600-630 nm for pS129 (Alexa Fluor 594), and 410-500 nm for TH. The pixel size was set to 15 nm (2056 x 2056 pixels), and the pinhole size and z-step were optimized for resolution and oversampling, supporting later deconvolution. Deconvolution was performed using Huygens Professional (Scientific Volume Imaging, Hilversum, the Netherlands) with a theoretical point spread function, custom background settings, default signal-to-noise ratio, and the CMLE algorithm. Image adjustments (color balance, contrast, and brightness) were made using Fiji ImageJ (RRID:SCR_002285).

#### Lysosomal functional assay

iDA neurons were exposed to PFF for 52 days, then stained with 1 µM LysoTracker DND-99 (ThermoFisher, Canada) for 45 min in maturation medium (STEMcell, Canada). After staining, neurons were rinsed once with 1x PBS containing Ca²⁺ and Mg²⁺ and fixed with 3% PFA and 0.1% glutaraldehyde for 15 min at RT. Neurons were then immunostained as described in the previous section for super-resolution STED microscopy, with adjustments to the primary and secondary antibody concentrations (LAMP2/AF405, 1:100; pS129/AF488, 1:400; TH/AF594, 1:400).

Lysosomal clustering was analyzed using Fiji ImageJ (RRID:SCR_002285). Two regions within each neuron were delineated to define: 1) the area containing α-syn inclusions and 2) the cytosolic region of the soma without inclusions. The corrected total cell fluorescence (CTCF) of LAMP2 (representing total lysosomes) was measured within both regions to evaluate lysosomal density inside and outside the inclusions. CTCF values were normalized to the surface area of each region (n=3, total of 37 neurons). To assess lysosomal function, fluorescence intensity of LysoTracker and LAMP2 was measured along a line crossing lysosomes (LAMP2) inside α-syn inclusions and in the cytosolic region (n=3, total of 19 neurons). The area under the curve for LysoTracker and LAMP2 fluorescence was calculated and normalized by distance using GraphPad Prism (RRID:SCR_002798).

#### Statistics

Statistical analysis was conducted using GraphPad Prism (RRID:SCR_002798) on data from a minimum of three independent experiments. One-way ANOVA was used to evaluate the results, followed by Tukey’s HSD post-hoc test for multiple comparisons (details on the compared groups are provided in the relevant figure legends). A p-value of less than 0.05 was considered statistically significant. One-way ANOVA followed by Dunnett’s multiple comparison test was used for Figure S3E, and an unpaired t-test for Figure 9.

## Acknowledgments

H.A.L were supported by EPFL. A.L.M.M and Y.J were supported by EPFL, Idorsia Pharmaceutical Ltd. and the Foundation Bru. M.L.N.B and C.U were supported by EPFL and the Foundation Bru. L.V.D.H, A.T, G.O, D.S, M.C, S.C.R, J.B, G.K, F.K were supported by EPFL. M.T, R.S, W.I and A.O were supported by the Canadian Institute of Health Research (CIHR) grant (FRN162400) and the Natural Sciences and Engineering Research Council (NSERC; RGPIN-2023-05581). We thank the CIME facility (EPFL) for the use of its electron microscopy facility and Dr. Galina Limorenko for her help with imaging some of the samples shown in Figure S9.

## Author contributions

H.A.L conceived the study. H.A.L and A.L.M.M supervised the study. H.A.L and A.L.M.M designed all the experiments and wrote the paper. A.L.M.M prepared the samples and performed the experiments in Figures 1-8 and Figures S3A-B, and S4-S10. A.L.M.M analyzed Figures 1-4, Figures 7P and Figures S3A-B, S4-S10, S13-S14 and prepared the final figures. L.V.D.H analyzed Figures 1H-I, Figure 5-7A-O, Figure 8 and Figures S4, Figures S10-11 and drew Figure 10. M.L.N.B prepared the samples, performed the experiments and analyzed Figures 1E-G, 4, S3A-B and S8. A.T prepared the samples, performed the experiments and analyzed Figures 1E-G and S3A-B. G.O prepared the samples, performed the experiments and analyzed Figure 1B-C. D.S prepared the samples, performed the experiments and analyzed Figure S5. Y.J prepared the samples, performed the experiments and analyzed Figure S2. U.C prepared the samples, performed the experiments and analyzed Figure S13C. M.T, R.S and W.I prepared the samples shown in Figures 9 and S3C. M.T, R.S and W.I confirmed Figures 1B, 2, and 3 in independent experiments (data not shown). M.T performed and analyzed the experiments shown in Figures 9 and S3C. M.C. prepared the samples for CLEM analysis and acquired EM images in Figures 1H, 6, 7A-O, 8, and S4 and S10-S11. S.C.R prepared the samples for CLEM analysis and acquired EM images in Figure 5A-G. J.B acquired EM images in Figure 6P. G.K supervised the experiments shown in Figures 1H, 6, 7A-O and contributed to the interpretation of the data. A.O supervised the study for Figures 9 and S3C and contributed to the interpretation of the data.

