## Supplemental Figures for "Recapitulating Parkinson’s pathology in human iPSC dopaminergic neurons reveals new mechanistic insights into Lewy body formation and heterogeneity"

**Running title:** Seeded alpha-synuclein aggregates in iPSC-derived neurons replicate key features of Lewy body pathology

**Keywords:** Parkinson, alpha-synuclein, aggregation, Lewy body, iPSC, post-translational modification, heterogeneity.

**Supplemental information - Figures titles and Legends**

**Figure S1**

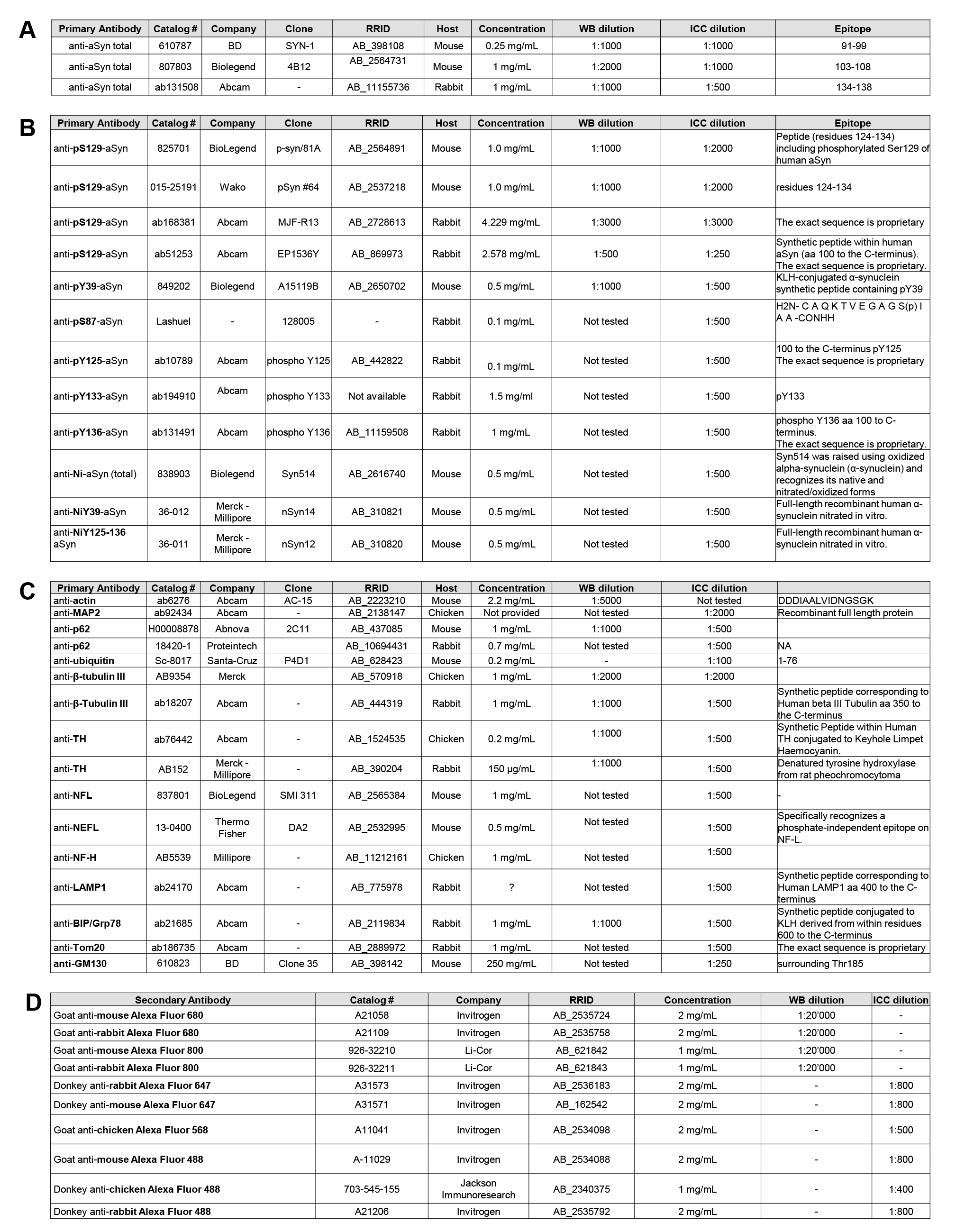

**Figure S1. List of the antibodies used in this study**

**A**. Antibodies used for the detection of total aSyn.

**B**. Antibodies used for the detection of aSyn PTM.

**C**. Other antibodies used in the study.

**D**. Secondary antibodies used for immunoblotting or confocal imaging.

**Figure S2**

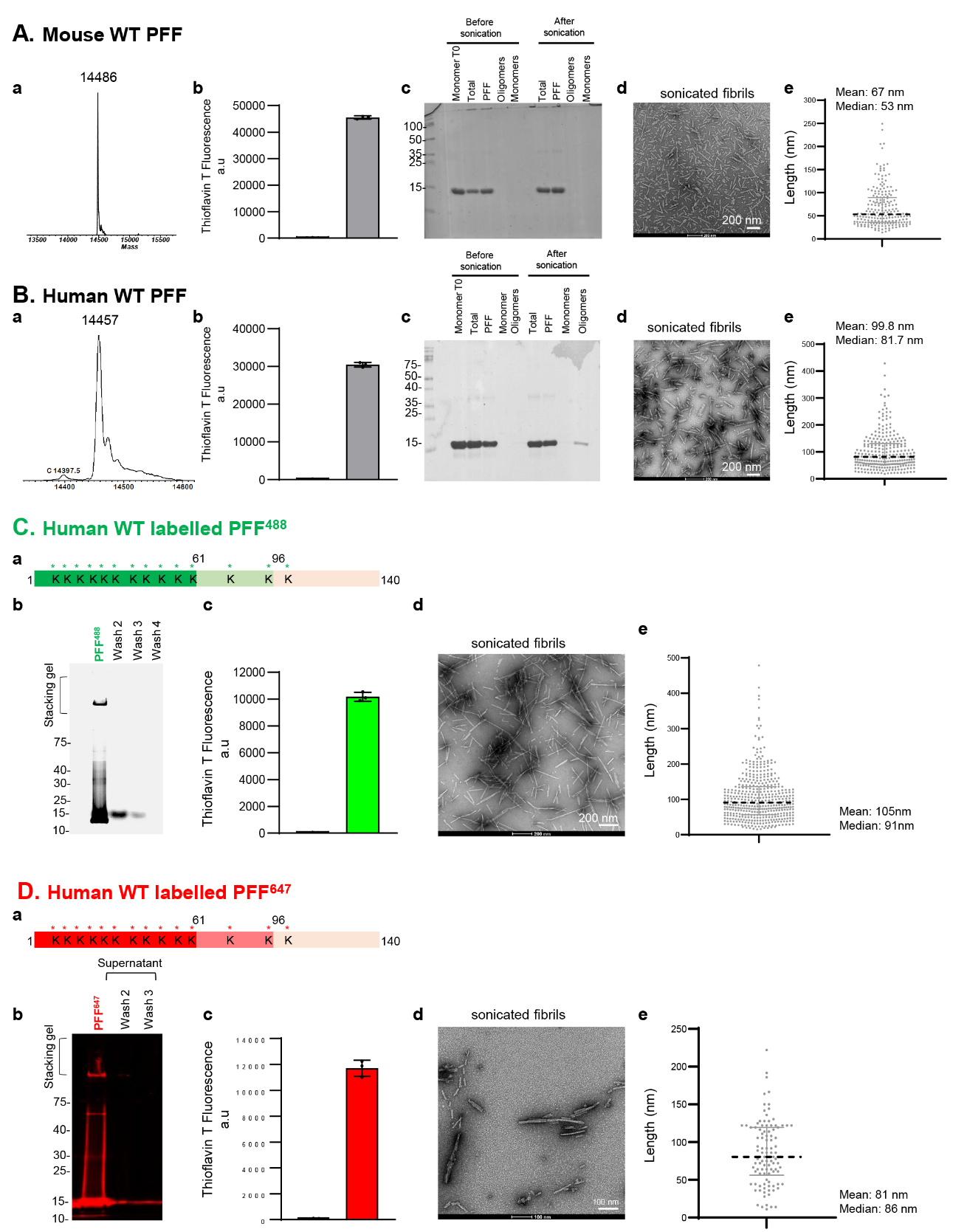

**Figure S2. Preparation and characterization of recombinant monomeric and PFF aSyn species.**

**A-D.** Mouse aSyn (**A**) and human (**B-D**) aSyn wild-type (WT) were produced in *E. coli* and purified by anion exchange chromatography and size-exclusion chromatography, followed by a final chromatographic step using reverse-phase HPLC, as previously described^1^. **a.** Purity and characterization of aSyn monomers. The purity of recombinant monomeric aSyn after purification was assessed by ESI-LC/MS, which showed the expected mass. **b-e.** Purity and characterization of aSyn PFF. aSyn PFF were formed by incubation of monomeric aSyn for 5 days at 37°C under constant agitation at 1000 rpm. After fibrilization, part of human aSyn PFF were labelled with Atto488 maleimide (**C**) or Atto647 (**D**). **b**. After sonication, PFF formation was assessed by ThT fluorometry. All data represent the average ± SD (n=3). **c**. The purity of aSyn PFF was verified by SDS-PAGE gel and Coomassie blue staining. After sonication, PFF preparations were centrifuged, and the presence of the PFF was verified in the pellet fraction, while the absence of monomer release after the sonication step was assessed in the supernatant fraction or after filtration through a 100 kDa filter (filtration). **d-e**. aSyn PFF were characterized by transmission electron microscopy (TEM) imaging. (**d**) Representative images of negatively stained aSyn PFF before and after sonication. All aSyn PFF showed the characteristic rigid non-branched fibrillar morphology. Scale bars = 200 nm (**Ad** and **Bd**) and 100 nm (**Dd**). **E**. The average length of the PFF after sonication.

**Figure S3**

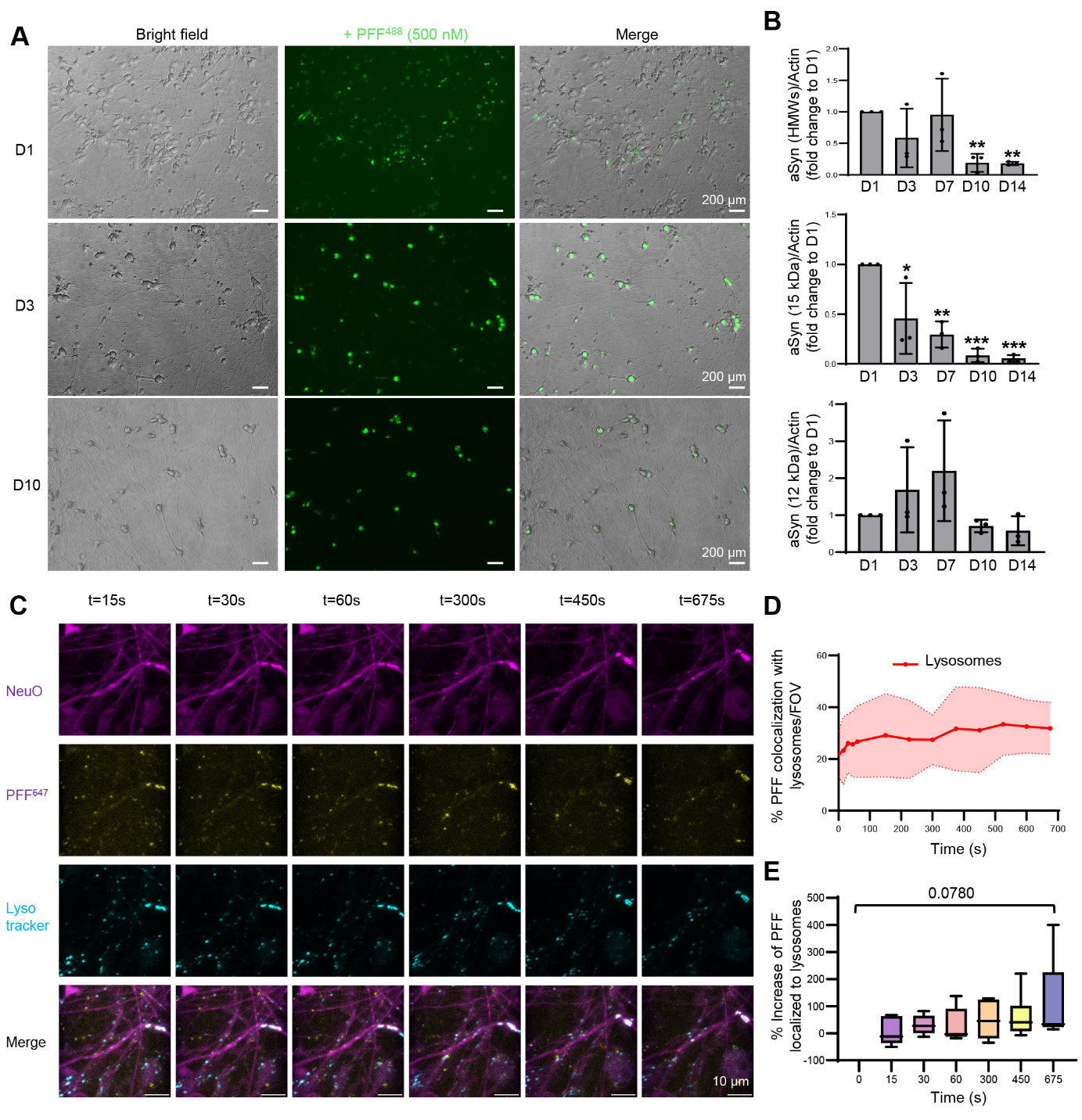

**Figure S3. Internalisation, processing, and clearance of aSyn PFF seeds over time in human iPSC-derived dopaminergic neurons.**

**A.** Individual channels of the images showing fluorescently labelled PFF (PFF^488^) internalised by iDA neurons at D1, D4, and D10 post-treatment in **Figure 1E**. Scale bars = 200 µm. **B.** WB analyses of immunoblots shown in **Figure 1G**. Levels of full-length aSyn (15 kDa) or truncated aSyn (12 kDa) or HMWs were quantified by densitometry and normalized to actin (see immunoblots in Figure 1G). Graphs represent the mean +/- SD of 3 independent experiments. *p<0.01, **p<0.001, ***p<0.0001 (ANOVA with Tukey HSD post-hoc test, D1 vs. others time-points). **C.** Representative live-cell confocal images showing iDA (NeuO, purple) at DIV10 exposed to labelled PFF^647^ (yellow) for 20 min and showing colocalization in real time with active lysosomes (lysotracker, cyan) right after the PFF settles. **D.** Graphical representation of the colocalization between PFFs and lysosomes over time, showing a stable interaction between both entities (20 to 30% colocalizing). **E.** Box plots are displaying the increase of colocalization signal between lysotracker and PFF^647^ over time.

**Figure S4**

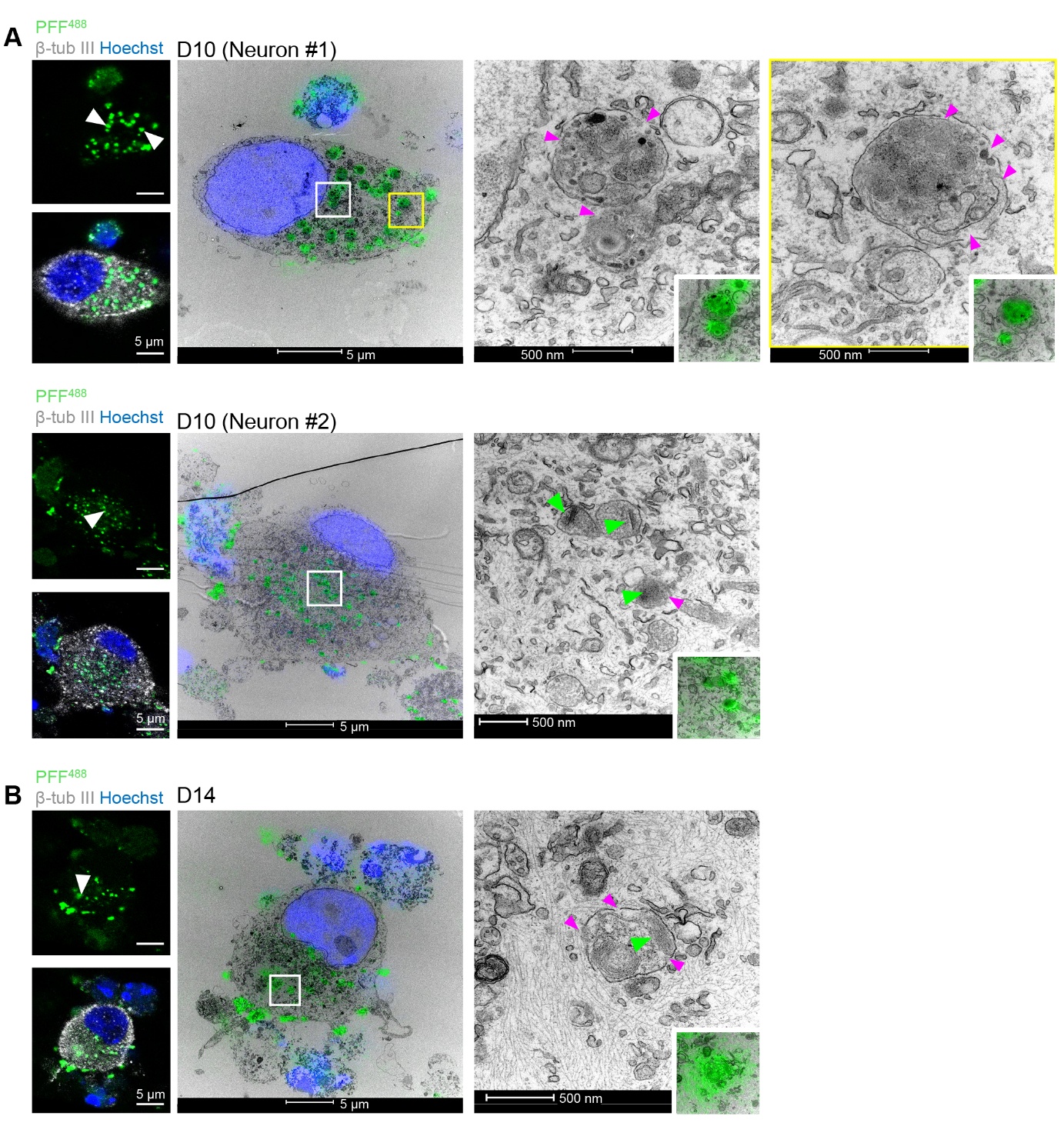

**Figure S4. Correlative light and electron microscopy of endolysosomal structures in neurons challenged with fluorescently labelled PFF (PFF^488^).**

**A.** Ten days after fibril addition (D10), PFF^488^ accumulate in large (>500 nm), membrane-bound endolysosomal structures containing membranes and other subcompartments. Rupture of the endolysosomal membrane is indicated by the pink arrow in Neuron #1.In another example (Neuron #2), PFF^488^-positive structures contained laterally associated fibrils (green arrows). **B.** A late endolysosome with a ruptured membrane (pink arrows) containing membranes and fibrils (green arrow). Scale bars = 5 µm and 500 nm.

**Figure S5**

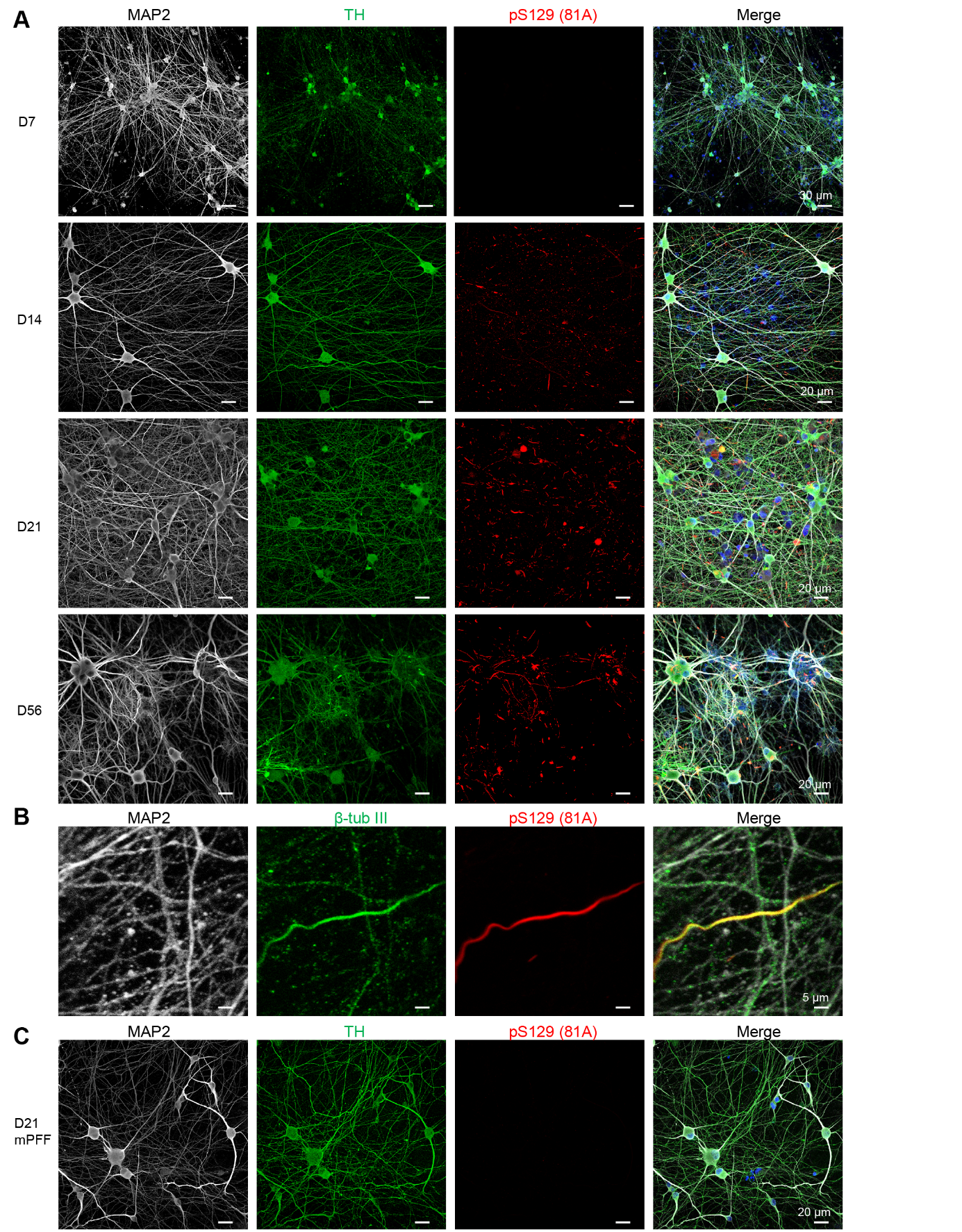

**Figure S5. Characterization of** **aSyn pathology formation** **in human iPSC-derived dopaminergic neurons over time.**

**A.** Individual channels of the images shown in Figure 1J illustrate the progression of pS129 pathology at D14, D21, and D56 following PFF treatment. pS129 aggregates were identified using the 81A antibody. Neurons were labelled with microtubule-associated protein 2 (MAP2) and Tyrosine Hydroxylase (TH) antibodies, while nuclei were counterstained with DAPI. Scale bars = 20 µm. **B.** pS129 neuritic pathology was colocalized with neurites positively stained by the β-tubulin III antibody but not with MAP2, indicating that the pS129 pathology localized predominantly in axonal neurites rather than dendrites. Scale bars = 20 µm. **C.** Mouse PFF failed to induce pS129 pathology (81A antibody staining) in iDA 21 days (D21) after PFF treatment. Neurons were labelled with microtubule-associated protein 2 (MAP2) and Tyrosine Hydroxylase (TH) antibodies, while nuclei were counterstained with DAPI. Scale bars = 20 µm.

**Figure S6**

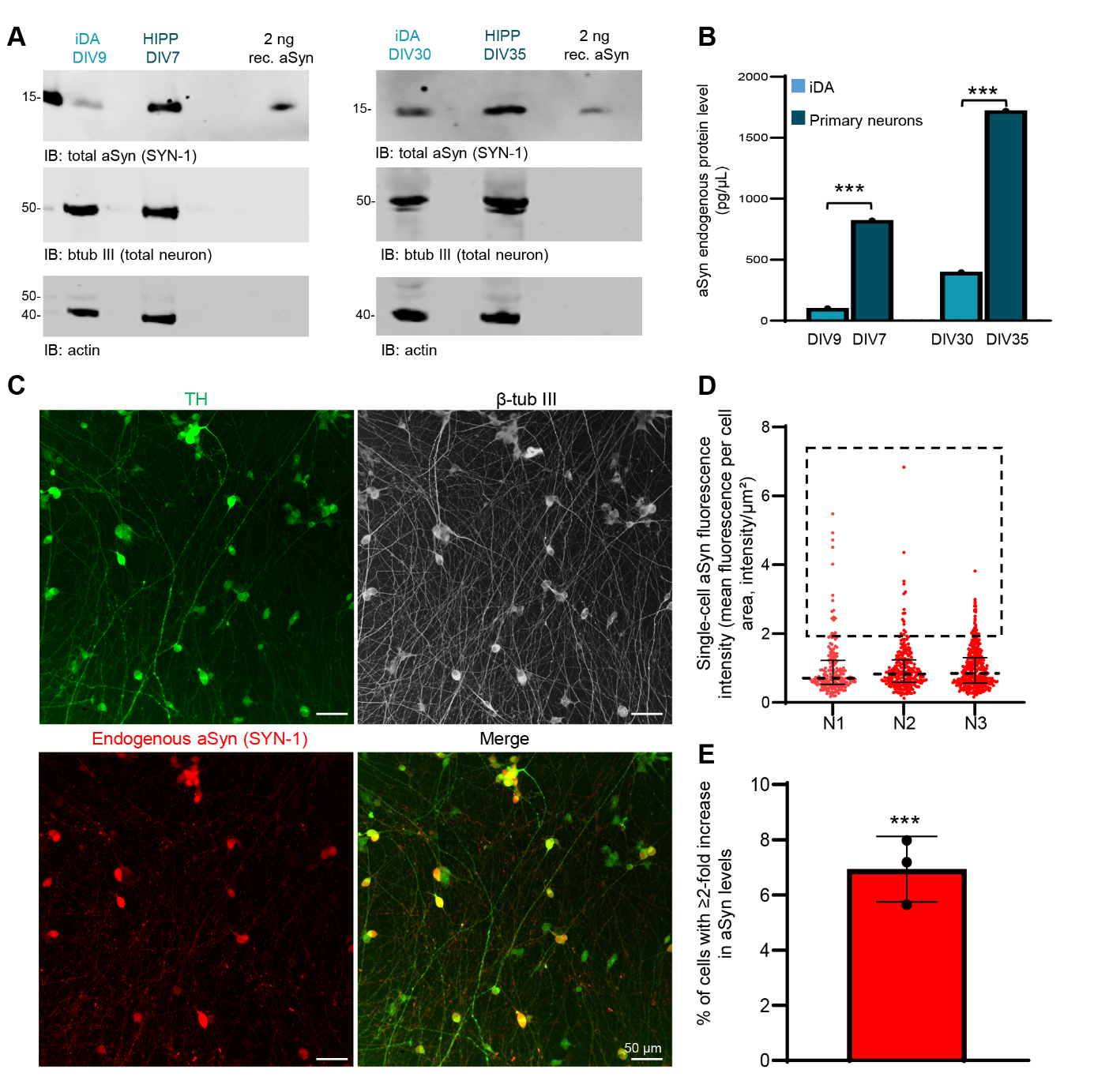

**Figure S6. Endogenous aSyn levels in iDA cultures.**

**A–B.** Western blot analysis comparing endogenous aSyn levels in iDA cultures and hippocampal primary neurons at two key time points: DIV9, when both cultures are treated with aSyn preformed fibrils (PFF) in the seeding model, and DIV30, when initial somatic aggregates begin to appear in iDA cultures following PFF addition. Total aSyn was detected using the SYN-1 antibody. Actin and β-tubulin III were used as loading controls, while TH staining was included to confirm dopaminergic differentiation of iPSC-derived neurons. **A.** Representative immunoblots show substantially lower aSyn expression in iDA cultures relative to primary neurons. **B.** Densitometric analysis reveals aSyn concentrations of ~104 pg/µL in iDA cultures at DIV9 versus ~825 pg/µL in primary neurons at DIV7, and ~400 pg/µL in iDA cultures at DIV30 versus ~1723 pg/µL in primary neurons at DIV35. Quantification was performed using the signal intensity of 2 ng of recombinant aSyn protein loaded on the same blot as a reference standard. **C-E**. Quantification of endogenous aSyn levels in iDA cultures using high-content imaging analysis to assess single-cell expression. iDA neurons were plated in 96-well plates and fixed at DIV9. Cells were immunostained for total aSyn (SYN-1 antibody), TH, and MAP2 to identify dopaminergic neurons and neuronal processes, respectively (**C**). For each independent experiment, three wells per condition were imaged, with nine fields of view acquired per well. Each experiment was independently replicated at least three times. For each detected cell, the median gray value of the aSyn fluorescence channel was used as a representative measure of intracellular aSyn levels. Mean fluorescence intensity per cell was normalized by the corresponding cell area and further normalized to the population mean of each independent experiment. The resulting data were visualized as violin plots (**D**). Graph **E** shows the percentage of cells exhibiting a ≥2-fold increase in α-synuclein levels, derived from the three independent experiments presented in Graph C. The quantified values correspond to the regions highlighted by the dashed boxes in Graph C. Statistical significance was determined using one-way ANOVA followed by Tukey’s HSD post hoc test. *p < 0.0001. Scale bars = 50 µm.

**Figure S7**

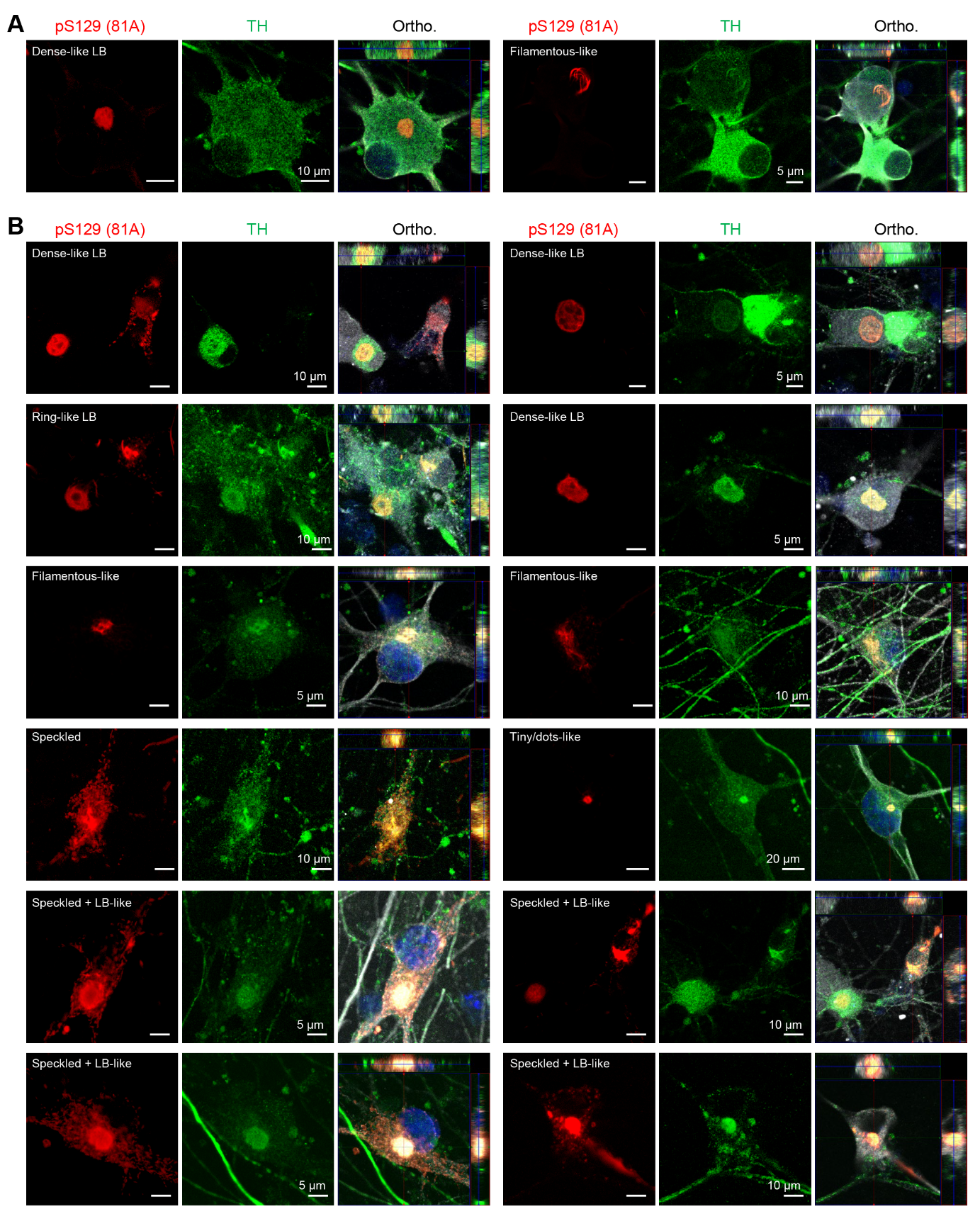

**Figure S7. Tyrosine Hydroxylase relocalization in pS129-positive aggregates.**

TH immunoreactivity was observed within pS129-positive aggregates (detected using the 81A antibody) at both D21 (**A**) and D56 (**B**). At D21, approximately 10–20% of aggregates showed TH relocalization, increasing to ~20–30% by D56. This relocalization occurred irrespective of aggregate morphology. Notably, TH signal intensity was significantly higher within the seeded aggregates than in the surrounding cytoplasm, which appeared nearly devoid of TH staining, indicating redistribution of TH into the aggregates. Nuclei were counterstained with DAPI. Scale bars = 5 or 10 µm.

**Figure S8**

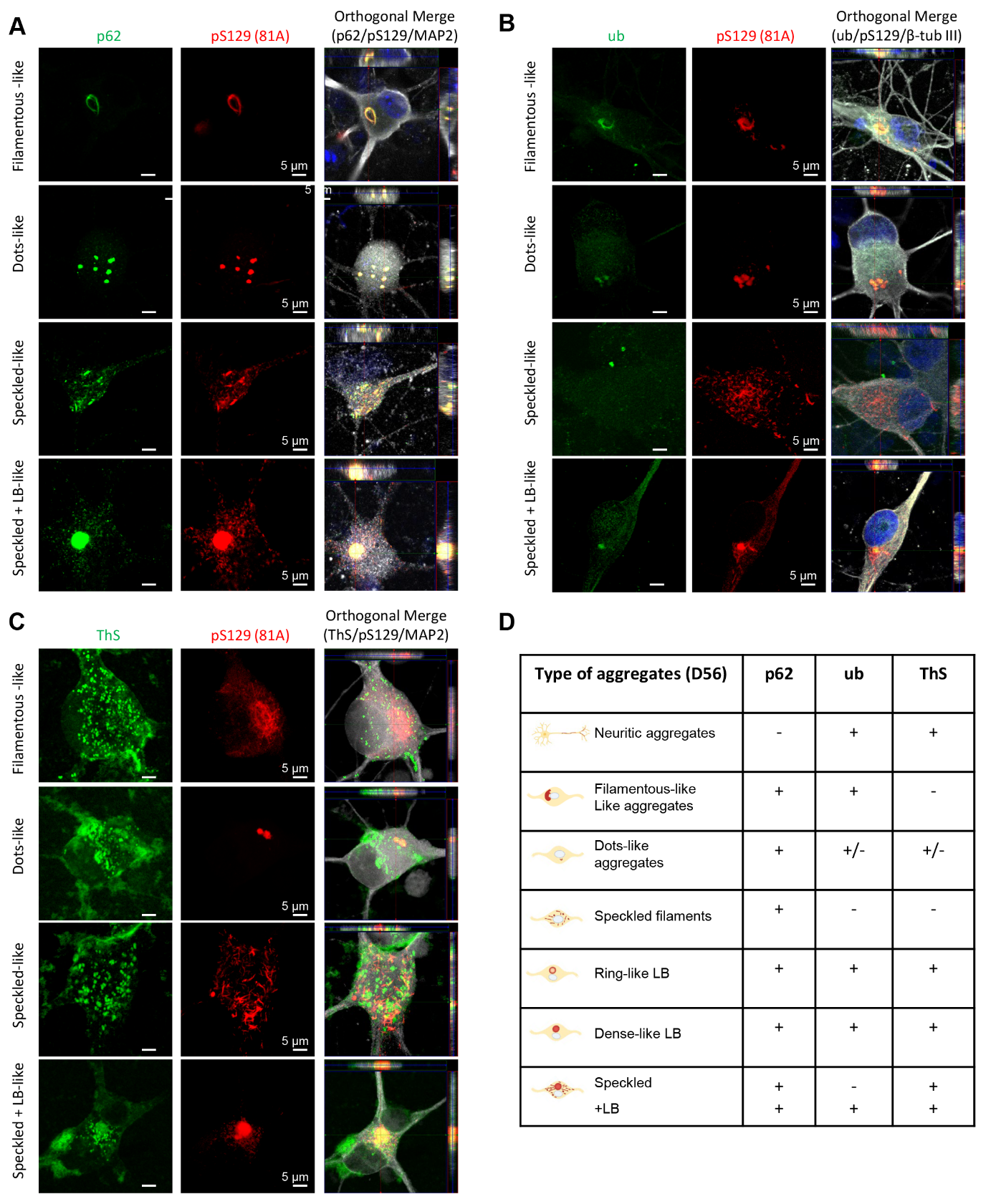

**Figure S8. Seeded aggregates in iDA neurons exhibit the LB pathological markers similar to human brain pathology.**

Representative images showing the differential sequestration of p62 (**A**), ub (**B**) and ThS (**C**) with pS129 pathology (81A antibody) in somatic pathology at D56. Neurons were stained with MAP2, while nuclei were counterstained with DAPI. Orthogonal projections (Ortho.). **Scale bars** = 5 µm. **D.** Summary table showing the differential sequestration of p62, ubiquitin, and ThS in different types of seeded aggregates at D56 (see additional staining in Figure 3).

**Figure S9 – Part I**

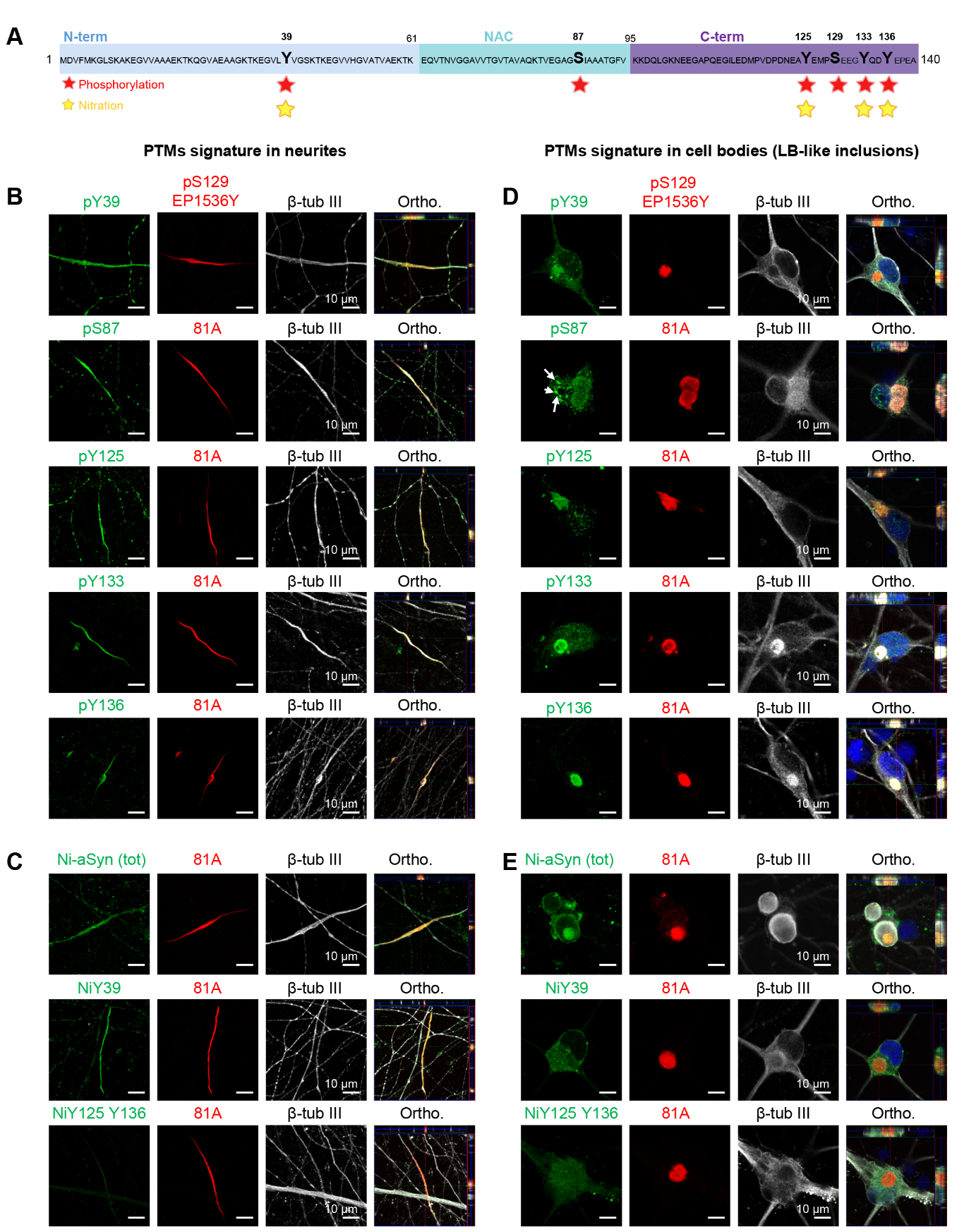

**Figure S9 – Part II**

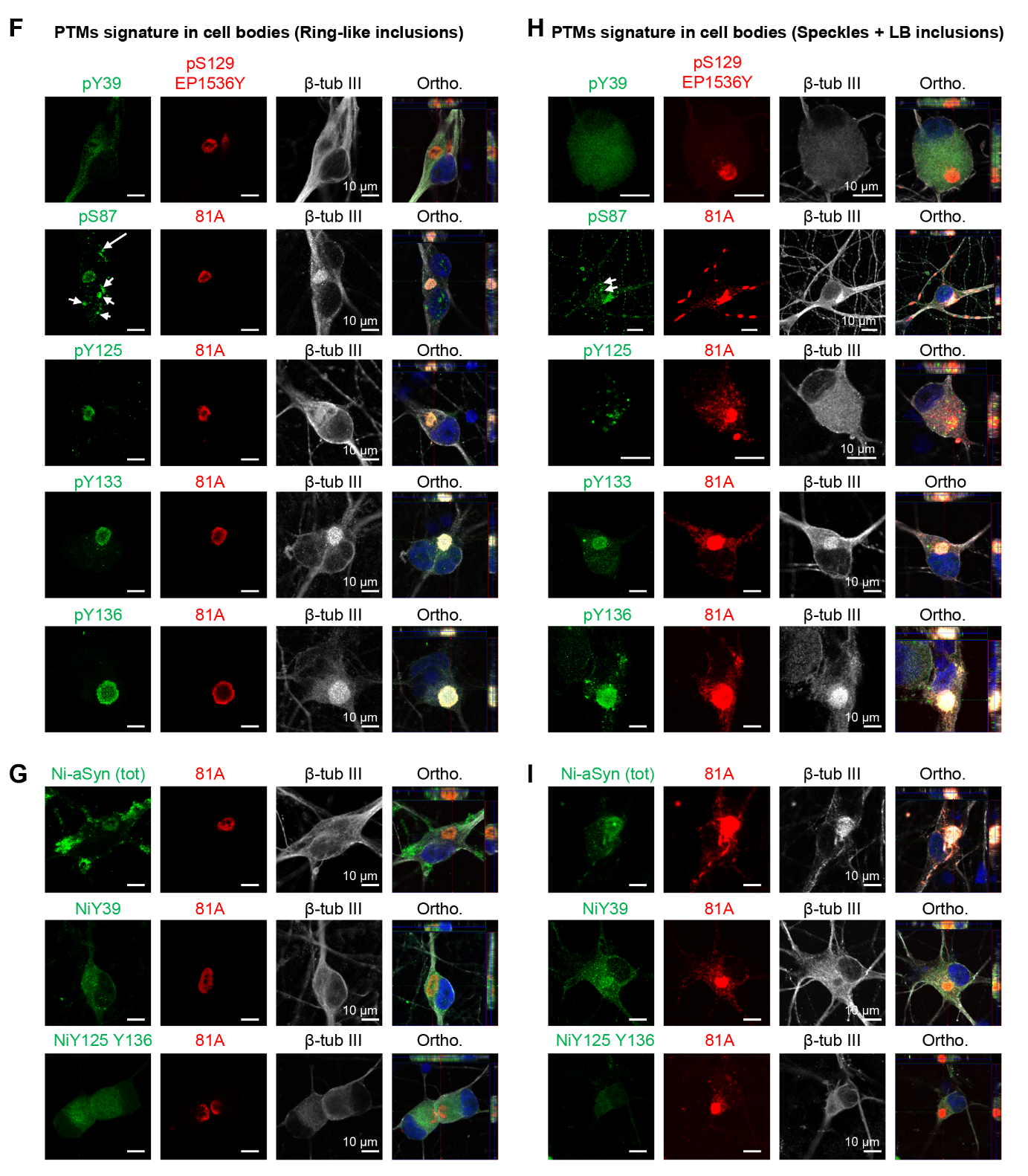

**Figure S9 - Part III**

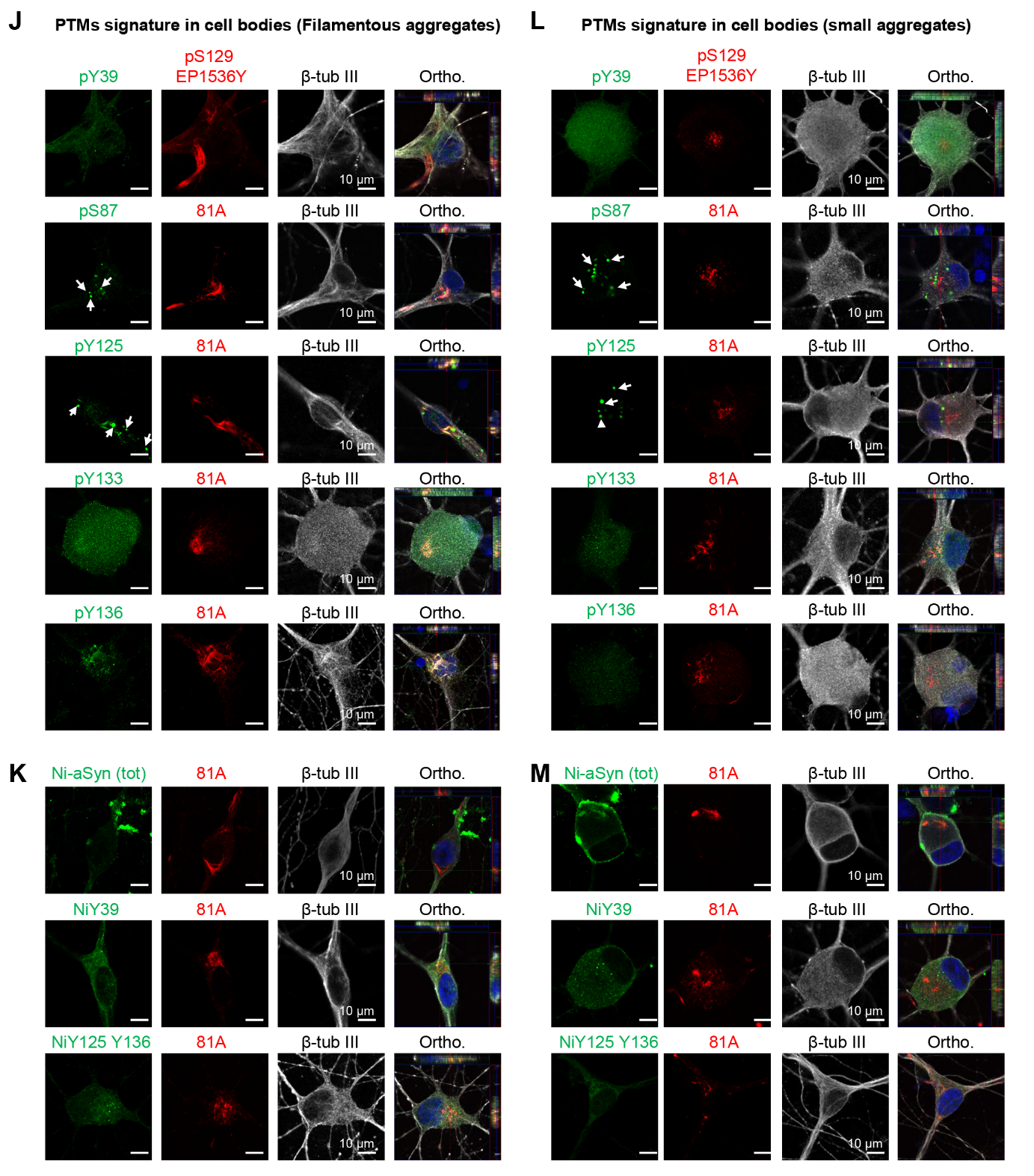

**Figure S9. The iDA seeding model replicates the PTM signature associated with human pathology.
A.** Schematic representation of PTM analyzed in this study. These include phosphorylation at residue Y39, S87, Y125, S129, Y133 and Y136, and nitration at Tyrosine residues (nY39, nY125, nY133, nY136). All of these PTM have been previously identified in aSyn pathology in post-mortem brain tissues from patients with PD and MSA. **B-M**. Representative images of pS129 pathology at D56 (B-C, neuritic pathology; D-E, dense LB inclusions; **F-G**, ring-like LB inclusions; **H-I**, Speckles and LB-like inclusions; **J-K**, filamentous aggregates and **L-M**, small aggregates) colocalizing with various PTM, including pY39, pS87, pY125, pY133, and pY136 phosphorylation (B, D, F, H, J and L) or with antibodies specific for nitration at Y39 (nY39) or Y125/Y136 (nY125/nY136) (C, E, G, I, K and M). White arrows indicate additional dot-like structures detected by some PTM antibodies, which are not pS129-positive. Neurons were stained with the β-tubulin III antibody, while nuclei were counterstained with DAPI. Orthogonal projections (Ortho.). Scale bars = 10 µm. The merged images shown in Figure S9 are also shown in the main Figure 4.

**Figure S10**

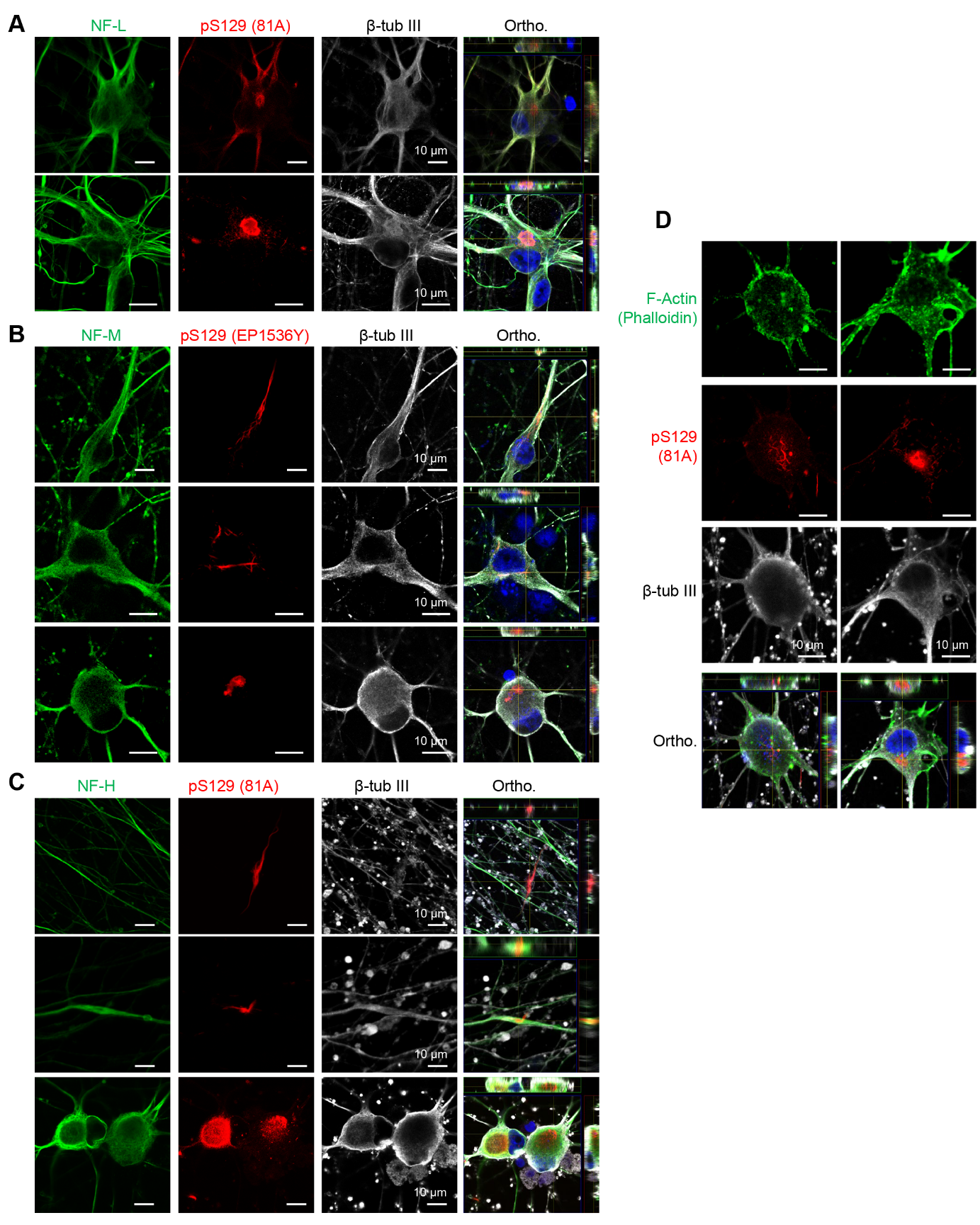

**Figure S10. pS129 pathology does not colocalize with major cytoskeletal proteins.**

ICC demonstrates the absence of colocalization between pS129 pathology and filamentous cytoskeletal proteins. **A.** Neurofilament light chain (NF-L, green) does not colocalize with pS129 pathology detected by 81A antibody (red) in somatic or neuritic regions. β-tubulin III (gray) staining highlights neuronal morphology, with orthogonal (Ortho.) projections confirming the lack of overlap. **B.** Similar results were observed for neurofilament medium chain (NF-M, green), with pS129 detected using EP1536Y antibody (red). **C.** Neurofilament heavy chain (NF-H, green) also showed no colocalization with pS129 pathology detected by 81A antibody (red). **D.** Phalloidin staining of F-actin (green) confirms no colocalization with pS129 pathology (81A antibody, red) in neuritic or somatic regions. Neurons were stained with the β-tubulin III antibody, while nuclei were counterstained with DAPI. Orthogonal projections (Ortho.). Scale bars = 10 µm.

**Figure S11**

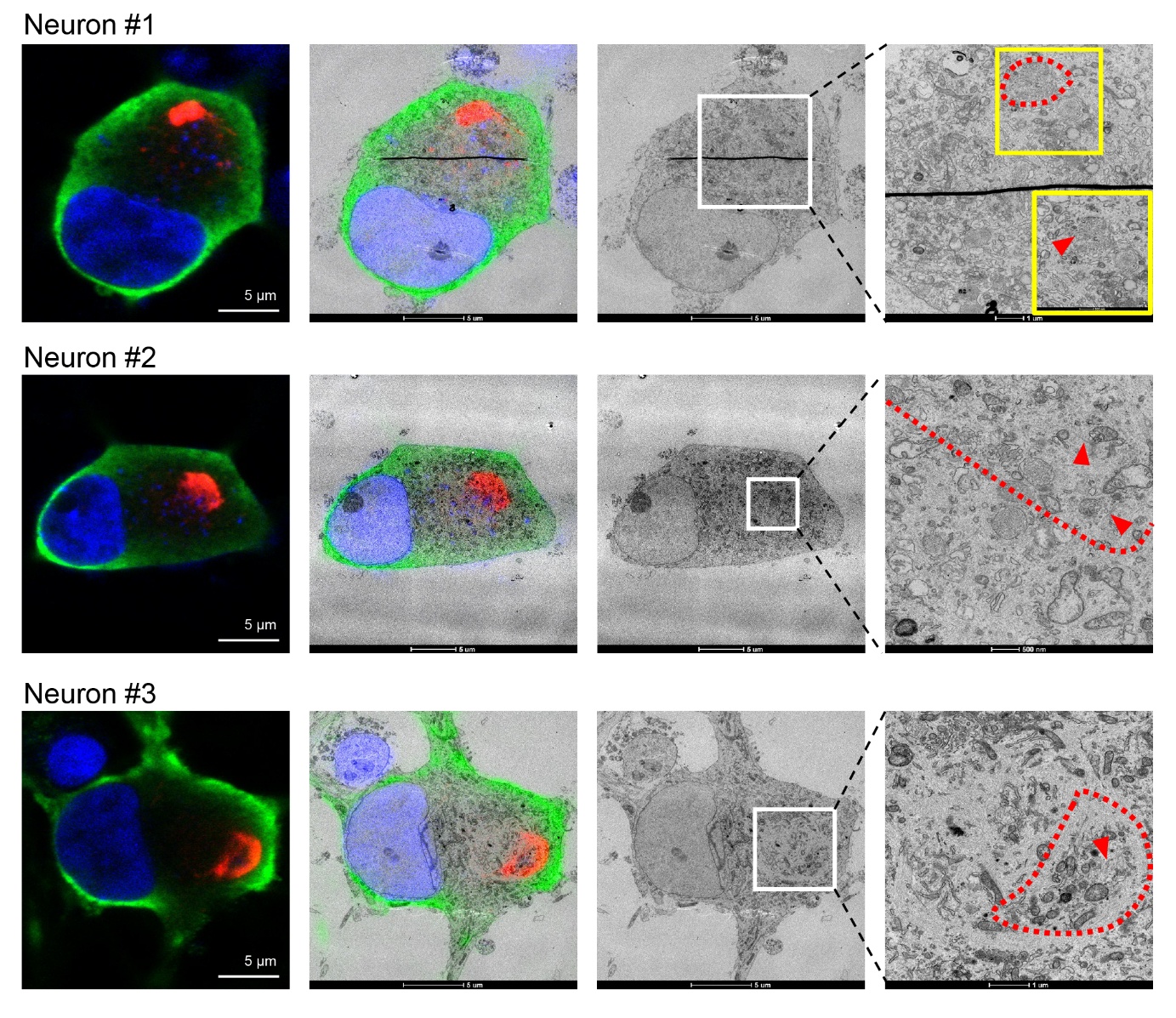

**Figure S11. Ultrastructure of dense D21 aggregates surrounded by speckled cytoplasmic staining**. The aSyn-pS129 immunofluorescence profile of the neurons #1, #2 and #3 consists of a dense centre surrounded by a punctate pattern of aSyn-pS129 immunoreactivity. The dense immunostaining colocalizes with a mixture of small vesicles, membranous organelles (notably mitochondria and lysosomes) and thick (13 ± 2.5 nm) fibrils that are randomly oriented (red arrows). The surrounding regions, with a speckled immunopositivity, correlate with thin (8 ± 2nm) filaments that curve throughout the cytosol. Scale bars = 500 nm or 1 and 5 µm.

**Figure S12 – Part I**

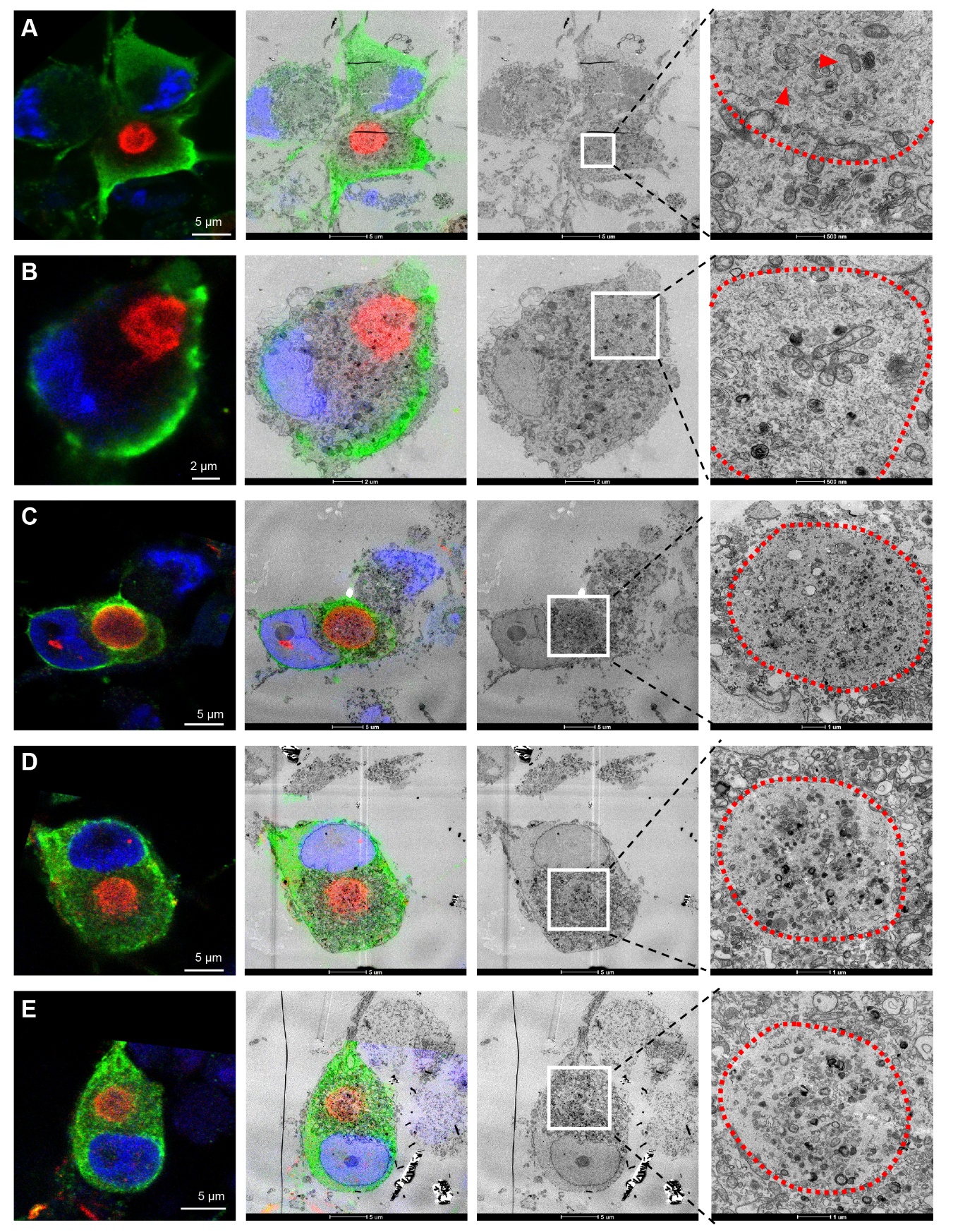

**Figure S12 – Part II**

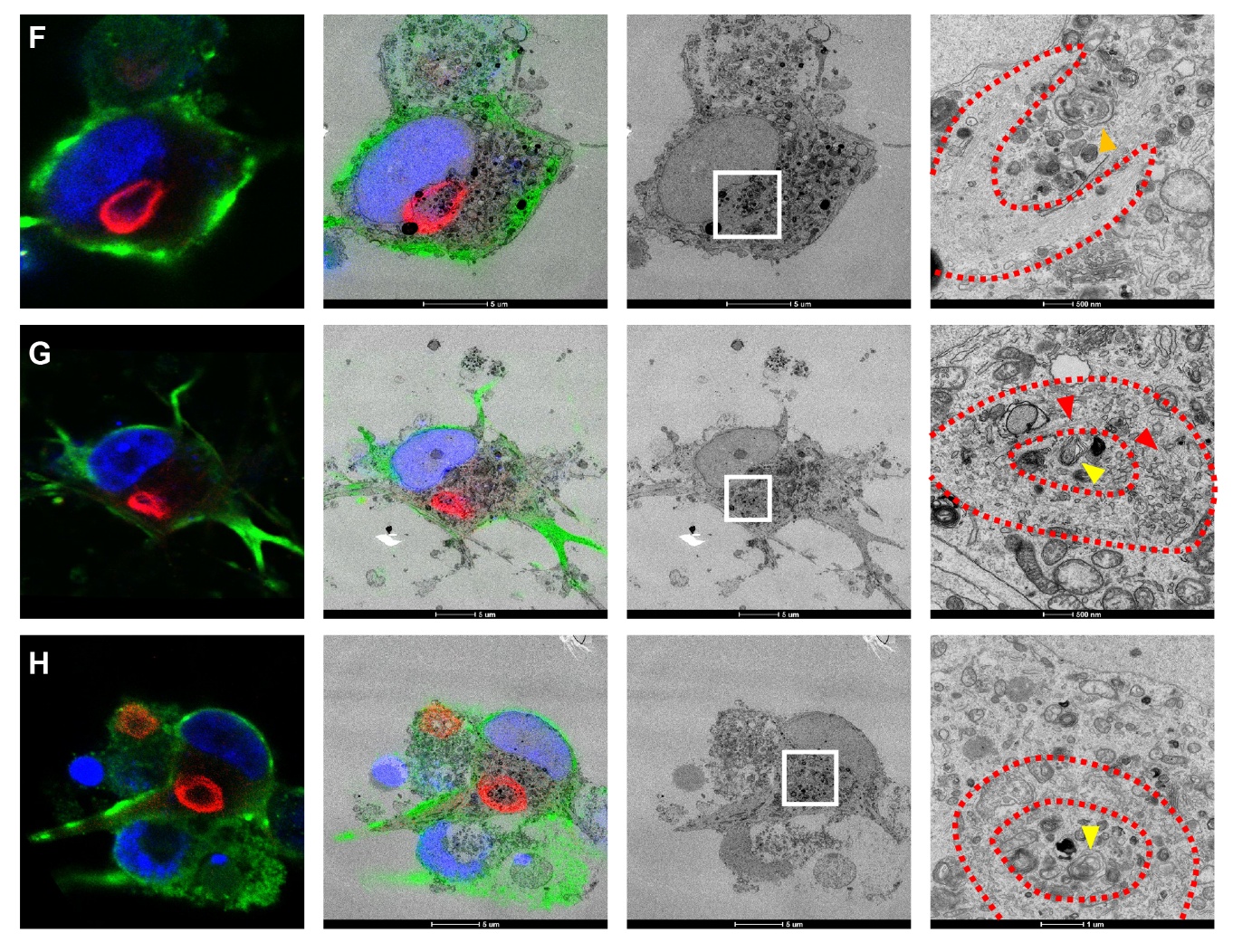

**Figure S12. Ultrastructure of the different types of LB at D56.** **A-E.** Ultrastructure of the dense LB. These aggregates share a membranous ultrastructure rich in small vesicles, a few thick fibrils (13.5 ± 2.5 nm, as indicated by the red arrow in **A**), clustered mitochondria (notably in **B**) and thin filaments (8 ± 2nm, notably in **C**, **D** and **E**). Note that the amount of thick fibrils in dense D56 aggregates is lower than in dense D21 aggregates, suggesting degradation of newly formed aSyn fibrils over time. Scale bars = µm. **F-H.** Ultrastructure of the ring-like LB. The centre of the ring, which is low in aSyn-pS129 immunoreactivity, correlates with the presence of autophagosomes (orange arrow), multilamellar bodies (yellow arrows) and other lipid-rich organelles. In **F**, the ring itself correlates with an ultrastructure that is rich in parallel filaments, whereas in **G** and **H,** the ring is a membranous region consisting of small vesicles, organelles, and thick filaments (13 ± 2.5 nm, red arrows). Scale bars = 500 nm or 1, 2 and 5 µm.

**Figure S13 – Part I**

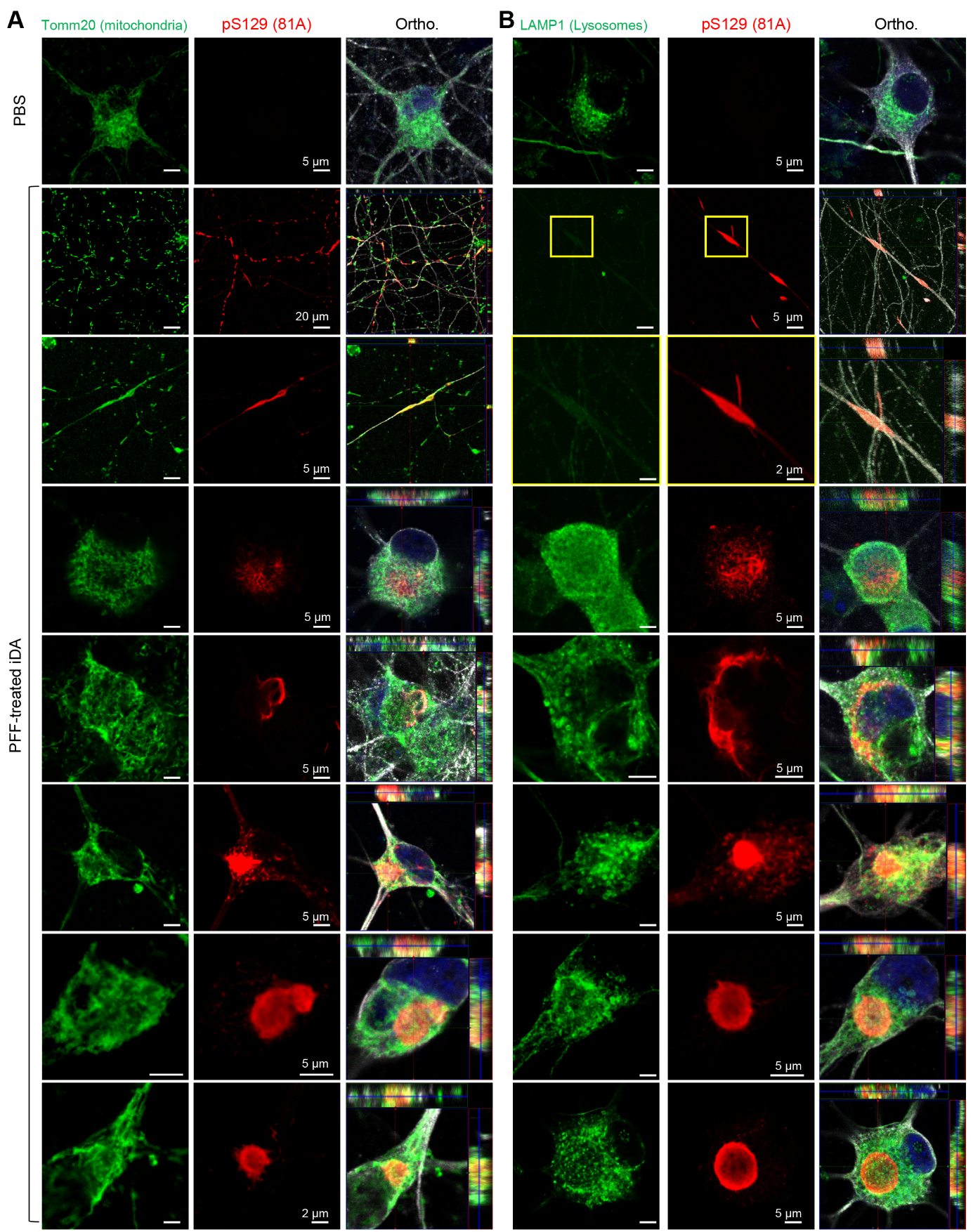

**Figure S13 – Part II**

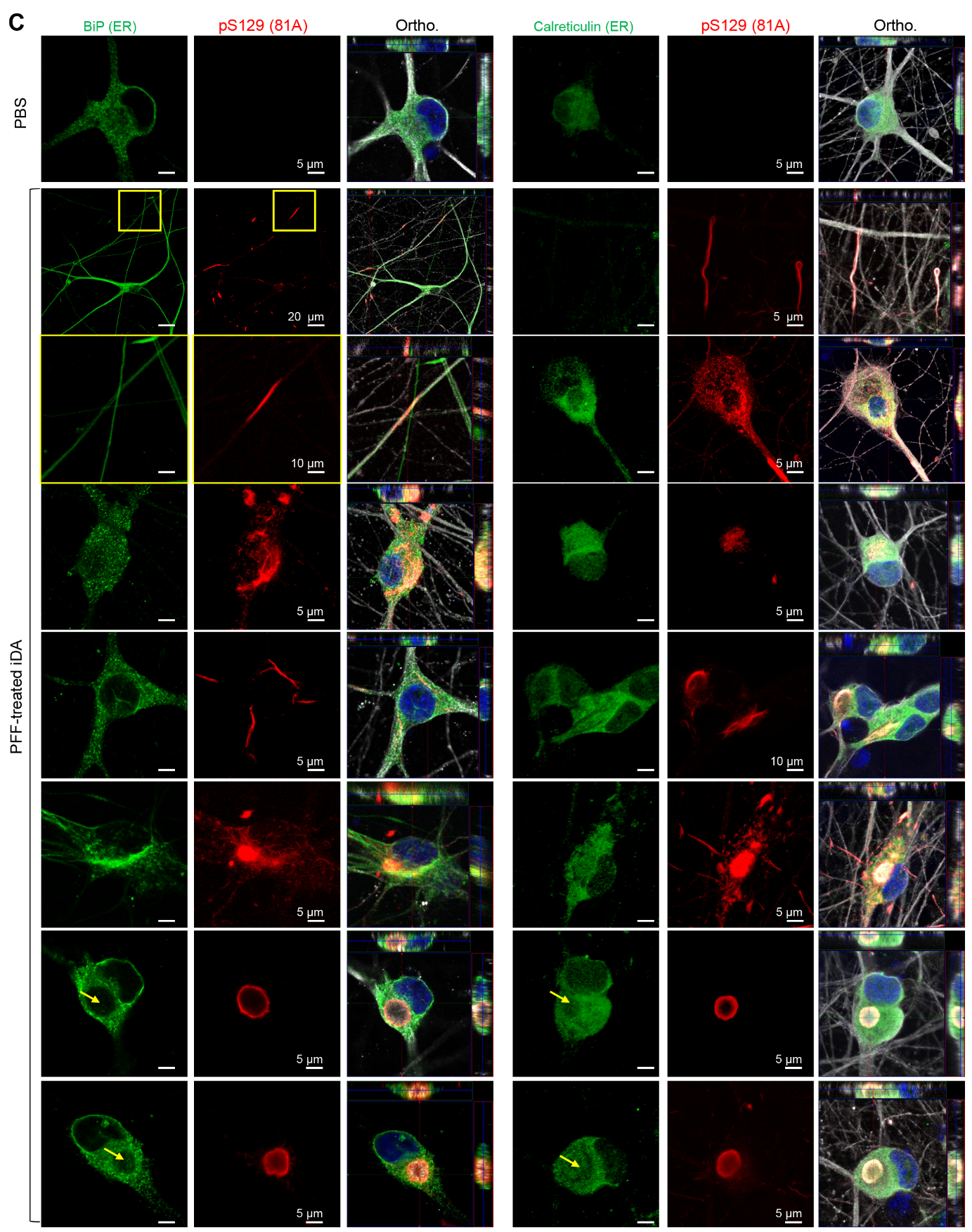

**Figure S13 – Part III**

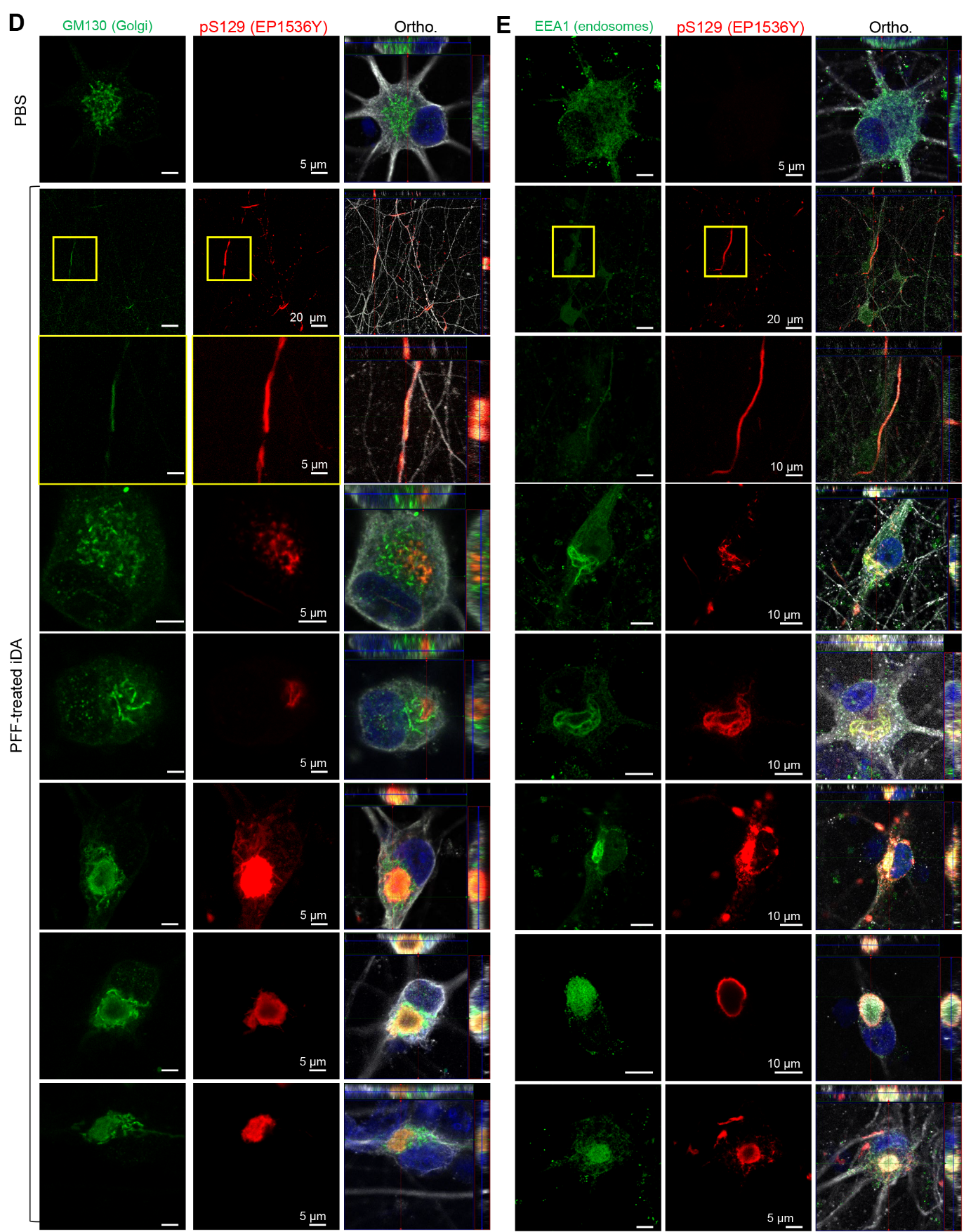

**Figure S13 – Part IV**

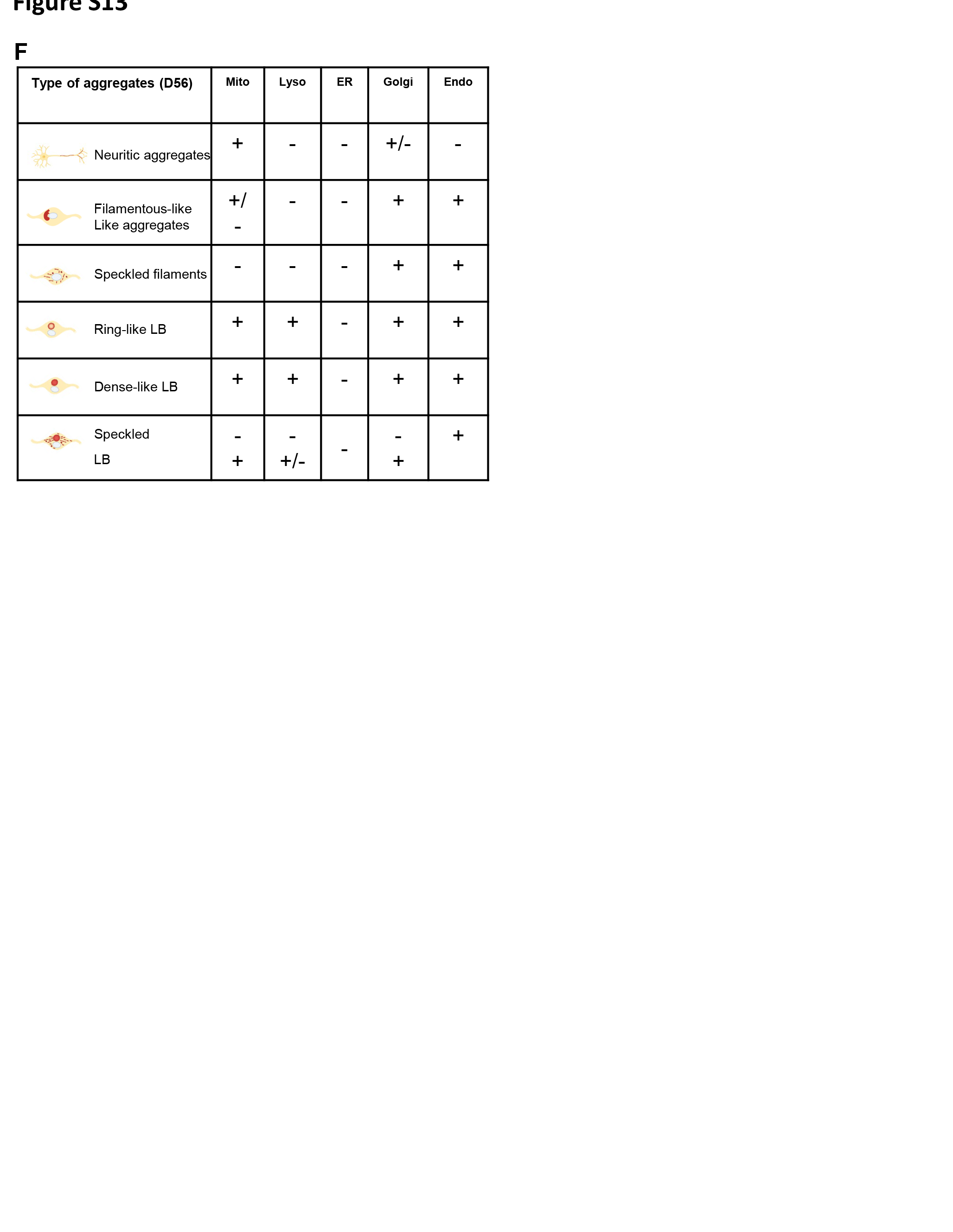

**Figure S13. Sequestration of key organelles into pS129 positive aggregates at D56 A-D.** Representative images show pS129 pathology (red, 81A antibody) alongside markers for key organelles: Tomm20 for mitochondria (**A**), LAMP1 for lysosomes (**B**), Bip for the endoplasmic reticulum (ER) (**C**) and GM130 for the Golgi apparatus (**D**). Neurons were stained with the β-tubulin III antibody, while nuclei were counterstained with DAPI. Orthogonal projections (Ortho.). Scale bars = 5, 10 or 20 µm as indicated for each panel. E. Summary table showing the differential colocalization of key organelles within the different types of seeded aggregates at D56 (see additional staining in Figure 7P).

**Figure S14**

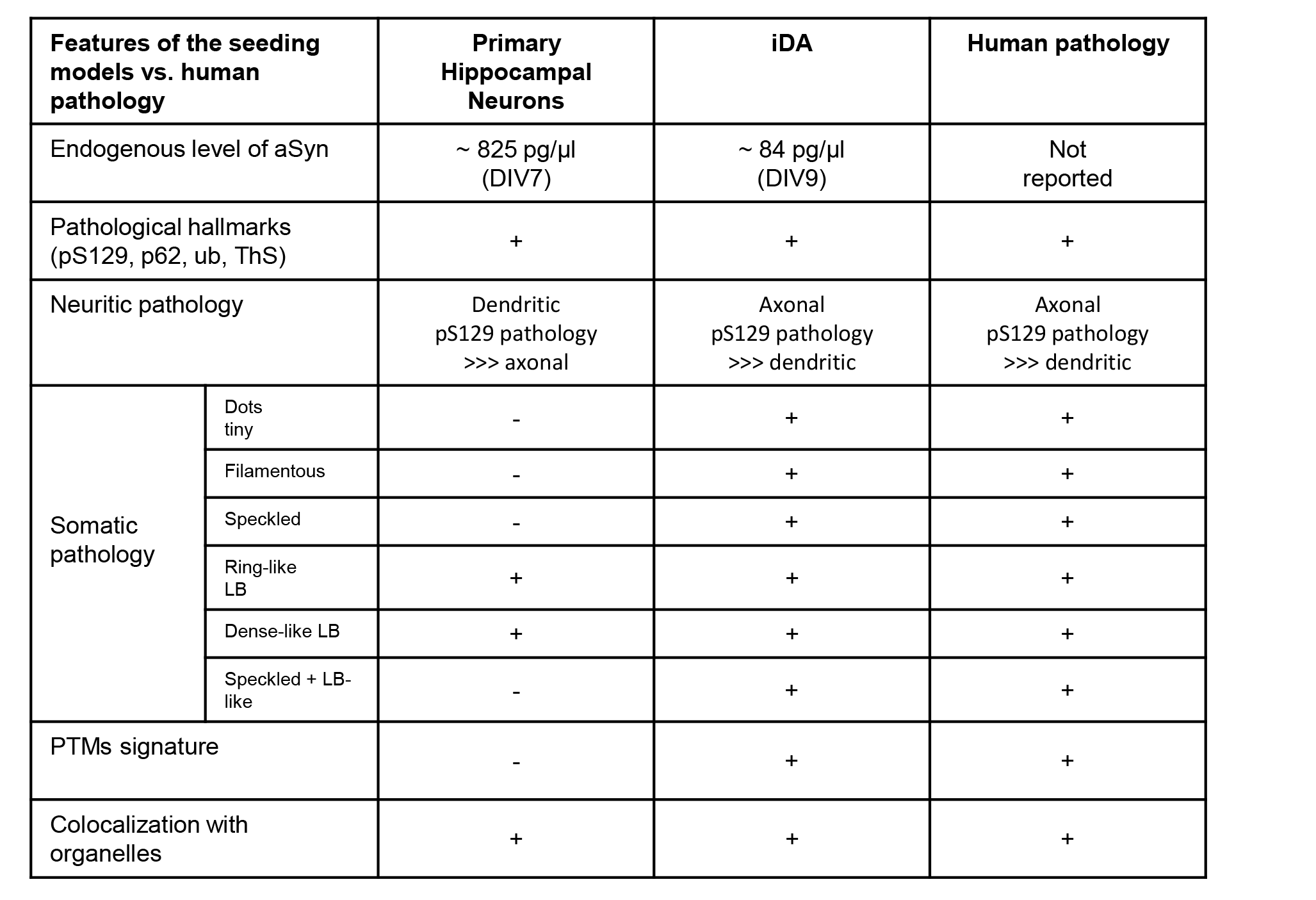

**Figure S14. Comparative analysis of aSyn seeding responses in primary hippocampal cultures and iDA models, and their resemblance to human pS129 pathology.**

This table summarizes key features of aSyn pathology observed in hippocampal primary neurons^2^ and induced dopaminergic (iDA) neurons following seeding, and compares these with the LB pathological hallmarks described in human brain tissue^3-5^. Parameters assessed include: **endogenous aSyn levels**, **neuritic distribution of pS129 pathology** (axon vs. dendrite)^6^, **aggregate morphological spectrum^7-9^**, **PTM signatures**, and **organelles sequestration^10-16^**.
